# Spatial Dependence and Heterogeneity in Molecular Imaging: Moran Quadrant Maps Enable Advanced Spatial-Statistical Analysis

**DOI:** 10.1101/2025.10.27.684518

**Authors:** Léonore E. M. Tideman, Felipe A. Moser, Lukasz G. Migas, Jacquelyn Spathies, Katerina V. Djambazova, Cody R. Marshall, Matthew S. Schrag, Eric P. Skaar, Jeffrey M. Spraggins, Raf Van de Plas

## Abstract

Multiplexed molecular imaging, such as imaging mass spectrometry, provides spatially-contextualized insights that are transforming biomedical research by advancing our understanding of tissue organization and disease mechanisms. However, the spatial-statistical properties of molecular imaging data are often underutilized, and scalable computational tools to analyze them are lacking. Here, we demonstrate how local and global spatial autocorrelation (SAC) metrics can be used to quantify spatial dependence and spatial heterogeneity, properties that violate the common assumption of independent and identically distributed measurements. We develop mathematically rigorous methods for SAC-based exploratory analysis. Furthermore, we introduce a novel spatial feature extractor, the Moran quadrant map, and develop two advanced workflows based on it: Moran-Felsenszwalb segmentation for tissue domain segmentation and Moran-HOG clustering for colocalization-based image analysis. Finally, our open-source Moran Imaging toolbox (https://github.com/vandeplaslab/Moran_Imaging) provides scalable Python implementations, including a novel parallelized spatial lag algorithm, unlocking SAC-based analyses and biological insights for large-scale imaging.

## 1 Introduction

Studying the anatomical and cellular organization of tissues is key to understanding the structural and functional complexity of the human body, in health and in disease [1, 2]. With tissue function regulated by molecular exchanges between cells, accounting for spatial context across different scales (*e*.*g*., cellular micro-environments or niches, multicellular functional units, anatomical regions) is essential. Multiplexed molecular imaging technologies, such as imaging mass spectrometry (IMS), imaging mass cytometry (IMC), multiplexed ion beam imaging (MIBI), multiplexed error-robust fluorescence in situ hybridization (MERFISH), multiplexed immunofluorescence (MxIF) and co-detection by indexing (CODEX) microscopy, address this need for spatially-contextualized molecular profiling, for instance in the Human Biomolecular Atlas Program (HuBMAP) [3, 4], Kidney Precision Medicine Project (KPMP) [5], and Human Tumor Atlas Network (HTAN) [6]. However, the statistical relationships between spatially-localized measurements are often underutilized, or even disregarded. Characterizing and exploiting this spatial-statistical information could significantly improve our understanding of the relationship between tissue organization and biological function, helping to unlock the full potential of molecular imaging.

Computational analysis for multiplexed molecular imaging encompasses many tasks, including tissue domain segmentation, image clustering and classification, and differential expression testing [7–9]. Remarkably, many approaches consider measurements within an imaging dataset (*e*.*g*., mass spectra in IMS) as separate measurements, disregarding their spatial coordinates and thus failing to account for their potential spatial dependencies. There are many examples of such spatially-agnostic machine learning and deep learning algorithms, for example in IMS analysis: random forests and neural networks for mass spectra classification [10, 11], matrix factorization methods (*e*.*g*., principal component analysis, non-negative matrix factorization) for dimensionality reduction [8, 9], and centroid-based or density-based clustering (*e*.*g*., *k*-means and HDBSCAN clustering algorithms) for spatial segmentation [8, 9, 12]. Using spatially-agnostic algorithms may be problematic because it implicitly assumes independent and identically distributed (i.i.d.) pixel content. However, the i.i.d. assumption is violated by two fundamental properties of molecular imaging data: spatial dependence and spatial heterogeneity. We will demonstrate the importance of explicitly accounting for these two properties in the context of exploratory analysis, tissue domain segmentation, and colocalization analysis.

Our work translates and extends spatial-statistical metrics from spatial econometrics and geospatial information science to molecular imaging analysis [13–15]. The first property to account for, spatial dependence, refers to the correlation between spatially-localized measurements that is determined by their spatial arrangement [16–18]. We use spatial autocorrelation (SAC) statistics to quantify the degree of spatial dependence between measurements by combining a measure of molecular similarity (or, more generally, content similarity) with a measure of spatial similarity. The overall spatial dependence within an image can be captured by a global SAC statistic, such as Moran”s *I* [19]. Positive global SAC is indicative of clustering (*i*.*e*., spatial proximity of similar values), whereas negative global SAC indicates dispersion (*i*.*e*., spatial proximity of dissimilar values) [18, 20]. In molecular imaging, positive global SAC implies that measurements from nearby locations are more likely to report similar content than distant measurements. Positive global SAC has been observed in IMS data [21–23], potentially arising from the acquisition process (*e*.*g*., pixel size) and the sample”s histological structure. Experimental factors, such as matrix application and re-crystallization in matrix-assisted laser desorption/ionization (MALDI) IMS, or analyte diffusion during fixation, tissue mounting, or washes, may also introduce dependencies. Since spatial dependence violates the independence assumption of two-sample distribution comparison tests (*e*.*g*., Wilcoxon”s rank-sum test), it can pose a problem for differential expression testing. In IMS, intra-sample comparisons that ignore spatial dependence are known to result in high false discovery rates [21]. In spatial transcriptomics, spatial-dependence-aware statistical tests and machine learning models have been developed to identify spatially-variable genes that differentiate tissue domains [24–27]. The second property, spatial heterogeneity, indicates that relationships between variables vary over space. It is the opposite of spatial stationarity, and therefore violates the identical distribution assumption[16, 28, 29]. In molecular imaging, spatial heterogeneity can be caused by varying spatial regimes across tissue domains (*e*.*g*., multicellular functional units, anatomical regions) or between healthy and diseased tissue. For example, in IMS studies of bacterial infection, the spatial heterogeneity of abscessed tissue can help identify molecular markers specific to the host-pathogen interface [30, 31].

Studying spatial heterogeneity involves relating local patterns to global trends, necessitating local SAC metrics to complement global SAC metrics [28, 32]. In spatial transcriptomics, software tools have been developed to promote the use of spatial statistics in gene expression analysis [33, 34]. However, these tools primarily target profiling-style modalities rather than rasterized ones [35], and their functionality is largely limited to exploratory analysis and the identification of spatially-variable genes. The importance of modeling spatial heterogeneity for understanding and predicting spatial processes has also been recognized by the geospatial AI community [36, 37]. Given that the robustness and generalizability of convolutional neural networks can be compromised by their assumption of spatial stationarity [38–40], spatial-heterogeneity-aware neural network architectures have been developed for geospatial data [39, 40]. Although such customized models can outperform spatially-agnostic approaches, they are often application-specific. We take a different approach: instead of integrating spatial-statistic-awareness into the model, we extract spatial statistics as separate features, allowing their combination with traditional SAC-agnostic computer vision methods, enabling new modular workflows and re-use across different signsal processing and machine learning tasks, and facilitating their use also in human interpretation.

This work first adapts local and global SAC metrics for rasterized molecular imaging. This enables efficient quantification of spatial dependence and heterogeneity, facilitating SAC-based exploratory analysis. We then introduce a novel univariate SAC-informed feature extraction method, the Moran quadrant map (MQM), on the basis of which we develop two workflows for multiplexed molecular imaging data. Our first workflow is Moran-Felsenszwalb tissue domain segmentation. By partitioning a sample into spatially-coherent regions, segmentation defines regions-of-interest for further analyses (*e*.*g*., differential expression analysis between anatomical regions) [41]. In IMS, multivariate segmentation is usually done by clustering pixels with similar mass spectra using, for example, *k*-means clustering or spatially-informed extensions thereof [8, 42]. In spatial transcriptomics, many segmentation workflows are based on graph neural networks and/or self-supervised contrastive learning [43, 44]. However, these methods lack scalable implementations, limiting their applicability to large datasets, such as IMS datasets with thousands to millions of pixels. We compare Moran-Felsenszwalb segmentation to both non-spatial and spatial *k*-means clustering [42] and to the BANKSY algorithm [45]. Our second workflow is Moran-HOG colocalization-based clustering. Colocalization refers to quantifying the spatial similarity between images [46–49]. Clustering colocalized ion images has many applications in IMS, ranging from technological quality control to improved analyte identification and biological process characterization. Multiple deep-clustering workflows have been developed for colocalized ion image clustering and isotope detection [47–49]. We evaluate our Moran-HOG clustering workflow against noise-robust deep-clustering (NRDC) [48] and DeepION [49]. Beyond the analytical advances demonstrated in ion- and photon-based modalities (IMS and MxIF), the open-source Moran Imaging toolbox (https://github.com/vandeplaslab/Moran_Imaging/) provides scalable implementations that can be easily used in other modalities as well, such as IMC and MER-FISH. These scalable algorithms, including a novel parallelized spatial lag algorithm, effectively unlock SAC-based analyses and biological insights for large-scale imaging.

## 2 Results

The results are organized into three sections. First, we demonstrate how local and global SAC statistics enable spatially-informed exploratory analysis. The next two sections demonstrate new workflows that leverage our SAC-based feature extractor, the MQM, to deliver advanced tissue domain segmentation and colocalization analysis. We benchmark them against state-of-the-art methods, evaluating both analytic performance and computational efficiency. To emphasize the broad utility of our methods, the results cover six distinct multiplexed molecular imaging case studies. These include ion- and photon-based modalities (*i*.*e*., MALDI-IMS, MxIF-microscopy), different molecular classes (*i*.*e*., proteins, lipids), and diverse tissue types (*i*.*e*., murine brain, human brain, human kidney, and zebrafish). Overviews of our case studies and datasets are provided in section 4.7 and Supplementary section 4, respectively. Supplementary section 3.3 provides hardware specifications used in the benchmarks. Although our focus is on utilizing spatial-statistical properties of molecular imaging data, in Supplementary sections 2.3 and 3.2 we also demonstrate how to remove their effect using Moran eigenvector spatial filtering. The Moran Imaging Python package and MALDI-IMS data necessary to reproduce our results are available on GitHub (https://github.com/vandeplaslab/Moran_Imaging) and Zenodo (https://zenodo.org/records/17399931).

### 2.1 Exploratory spatial analysis

Exploratory spatial analysis is a form of exploratory data analysis that explicitly accounts for spatial dependence and spatial heterogeneity. Using the Moran Imaging package, we demonstrate how to perform univariate exploratory spatial analysis of molecular imaging data. We quantify SAC at both local (*i*.*e*., pixel-wise) and global (*i*.*e*., image-wide) levels by adapting established metrics - Moran statistics and Moran”s *I* - to rasterized imaging. As discussed in sections 4.1, 4.2, and 4.3, this adaptation involves nuanced, yet important, modifications to standard (geo)spatial statistics metrics because molecular imaging datasets often have a large number of isolates, which must be accounted for in the spatial weights matrix and the SAC metrics, and a high pixel count, which requires computationally efficient implementations. Furthermore, to facilitate discovery of local patterns of spatial association and anomalies, we propose casting the Moran scatterplot to the spatial domain as a MQM.

The first case study quantifies the global SAC of all ion images in MALDI-IMS dataset n^*o*^1, which measures a coronal rat brain section of a Parkinson”s disease model [50]. Each of its 809 ion images report 21,971 pixels, of which 4,007 are isolates (18% off-tissue). The global SAC of each ion image is estimated using Moran”s *I* (section 4.2), with Queen-contiguity and a row-standardized 1^st^-order spatial weights matrix (section 4.1). Out of 809 ion images, 806 report a statistically significant positive Moran”s *I*, indicative of positive global SAC. None exhibit a statistically significant negative Moran”s *I*, aligning with previous observations that negative global SAC is rarely observed [18]. Figure 1 presents six example images with corresponding Moran”s *I* statistics, and illustrates how images can be compared using Moran”s *I*, for example, to discard molecular species with noisy distributions. Each ion image is identified by its mass-to-charge ratio (*m/z*). We observe that spatially-coherent (*i*.*e*., structured) images tend to exhibit a high Moran”s *I*, whereas noisy images tend to have a lower Moran”s *I*. To determine whether a given Moran”s *I* is statistically significant, we compare the observed Moran”s *I* to the reference distribution obtained by randomly permuting intensity values across pixels (section 4.2). A pseudo-*p*- value is computed (Equation 8) with *M* = 999 spatial permutations, and is compared to the significance threshold *α* = 0.002. The ion image for *m/z* 5485.974 (Figure 1a) has a Moran”s *I* of 0.822 and a pseudo-*p*-value of 0.001 (*i*.*e*., the minimum achievable pseudo-*p*-value). With 0.001 *< α*, we conclude that this Moran”s *I* is statistically significant and that the distribution of *m/z* 5485.974 exhibits positive global SAC. Similarly, the ion image for *m/z* 4111.242 (Figure 1b) appears both spatially-coherent and biologically-informative. However, the ion image for *m/z* 12790.625 (Figure 1f) is not biologically-informative. With its Moran”s *I* = − 0.003 close to zero and pseudo-*p*-value 0.12 ≫ 0.002, its Moran”s *I* is not statistically significant, so we conclude that the distribution of *m/z* 12790.625 is spatially random. While spatial-coherence alone does not guarantee biological relevance, the observed correlation suggests potential for automated SAC-based differentiation between biologically-informative patterns and noise. This is valuable in massively multiplexed imaging modalities that measure hundreds to thousands of molecular species concurrently. Furthermore, measuring the global SAC of each ion image allows us to determine for which molecular species the i.i.d. assumption is not verified.

**Fig. 1.**
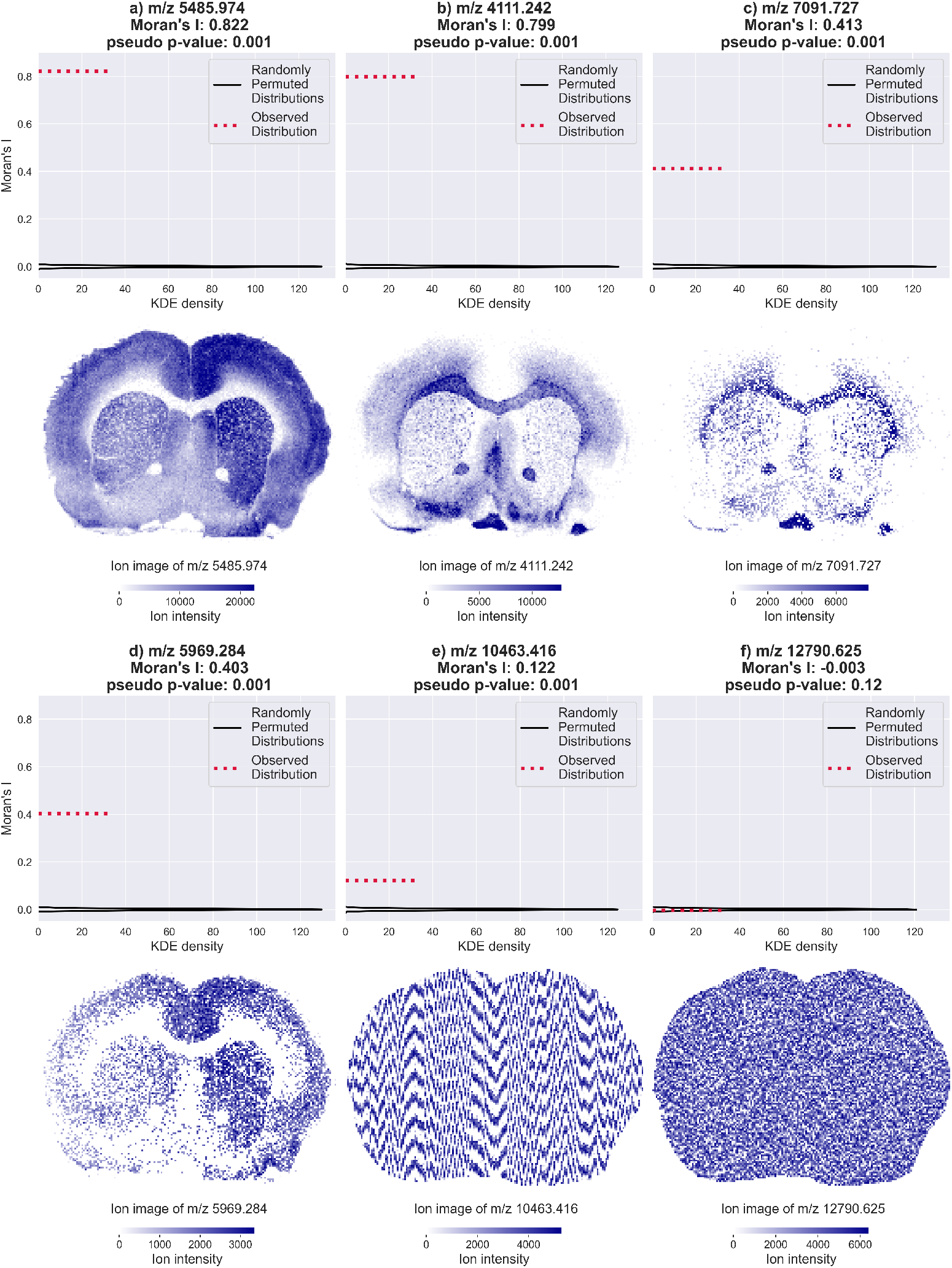
Quantification of global spatial autocorrelation (SAC) in ion images using Moran”s *I* and statistical assessment of significance through permutation testing. Moran”s *I*s are computed for all 809 ion images in IMS dataset n^*o*^1 (six shown). **a**, Ion image *m/z* 5485.974 exhibits a statistically significant Moran”s *I* of 0.822. **b**, Ion image *m/z* 4111.242 exhibits a statistically significant Moran”s *I* of 0.799. **c**, Ion image *m/z* 7091.727 exhibits a statistically significant Moran”s *I* of 0.413. **d**, Ion image *m/z* 5969.284 exhibits a statistically significant Moran”s *I* of 0.403. **e**, Ion image *m/z* 10463.416 exhibits a statistically significant Moran”s *I* of 0.122. **f**, Ion image *m/z* 12790.625 exhibits a statistically insignificant Moran”s *I* of -0.003. Noisy images exhibit lower Moran”s *I* statistics than spatially-coherent images, demonstrating that Moran”s *I* could be used to identify molecular species whose spatial distributions are more likely to report biological structure. Furthermore, the statistically significant positive global SAC of these ion images illustrates that the i.i.d. assumption is often unwarranted in molecular imaging.

The second case study focuses on local SAC quantification and demonstrates the MQM feature extractor on MALDI-IMS dataset n^*o*^1. Figure 2 shows how Moran”s *I* (section 4.2) and local Moran statistics (section 4.3) measure global and local SAC, respectively, and how the latter can be visualized as a spatial autocorrelation map (SAM). Although ion images map molecular distributions, local intensity variation - especially in noisy, low-intensity regions - can hinder threshold-based delineation of histological structures. Instead of local ion intensity values, the SAM reports local intensity correlations, making it helpful for discerning regional differences in spatial patterning (*e*.*g*., varying tissue architecture across anatomical regions). Subsequently, we demonstrate how the four quadrants of the Moran scatterplot can be recast as a MQM (section 4.4). In addition to Moran”s *I* quantifying spatial dependence in an image, we introduce the Moran scatterplot”s coefficient of determination *R*^2^ as a means of quantifying spatial heterogeneity (section 4.4). The ion image for *m/z* 4111.242 (Figure 2a) yields a Moran”s *I* of 0.799, indicative of positive global SAC (*i*.*e*., image-wide clustering). However, Moran”s *I* does not allow for localizing spatial clusters and outliers within the image. This necessitates local Moran statistics. Specifically, in the SAM (Figure 2b), we observe spatial clustering (*i*.*e*., positive Moran statistics) in the corpus callosum, anterior commissure, optic chiasma, and olfactory nerve tracks. Spatial outliers (*i*.*e*., negative Moran statistics) are observed in the corpus striatum. The magnitude of the positive Moran statistics exceeds that of the negative Moran statistics. The *R*^2^=0.8198 (Figure 2c) reports the image as spatially stationary, and the majority of pixels (88%) belong to the high-high or low-low quadrants (*i*.*e*., they are members of spatial clusters). Based on these clusters, the MQM (Figure 2d) clearly delineates histological structures. The ion image for *m/z* 12790.625 (Figure 2e) has a Moran”s *I* of -0.003, indicating spatial randomness. Its SAM (Figure 2f) shows randomly distributed positive and negative Moran statistics, with a similar magnitude range for both. The *R*^2^=0.0001 (Figure 2g) indicates spatial heterogeneity, and only half of the pixels (49%) belong to the high-high or low-low quadrants, with the other half being spatial outliers. The MQM (Figure 2h) also reflects this lack of spatial-coherence. Figures S9-S10 present analyses for *m/z* 5485.974 and 10463.416.

**Fig. 2.**
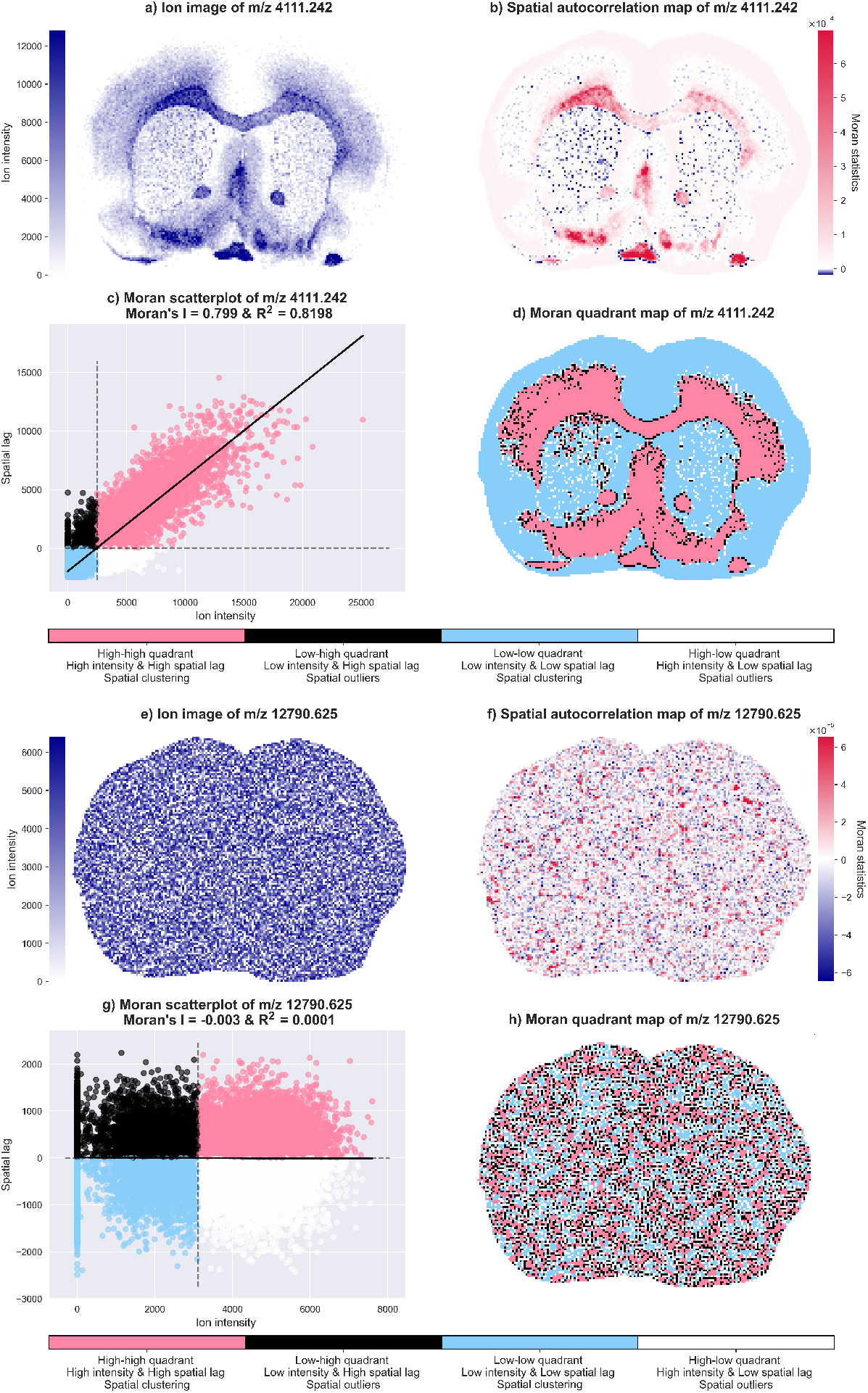
Quantification of local spatial autocorrelation (SAC) in ion images and demonstration of spatial autocorrelation map (SAM) and Moran quadrant map (MQM) feature extractors. Exploratory spatial analyses of two ion images from IMS dataset n^*o*^1 are shown, one with high and one with low global SAC. **a**, Ion image measured for *m/z* 4111.242 with a global Moran”s *I* of 0.799. **b**, SAM for *m/z* 4111.242, showing this molecular species” local Moran statistics. The SAM reports local correlation while the ion image reports ion intensity values. **c**, Moran scatterplot for *m/z* 4111.242 with *R*^2^ = 0.8198. The scatterplot ties the global SAC of an ion image to its local SAC. The coefficient-of-determination *R*^2^ quantifies the spatial heterogeneity in the image, its high value suggesting spatial stationarity for *m/z* 4111.242”s distribution. **d**, MQM for *m/z* 4111.242. The MQM feature extractor yields a four-label image, further categorizing the local SAC values reported by the SAM. The pink and blue regions exhibit spatial clustering, the former capturing high intensity clustering and the latter low intensity clustering. The black and white pixels in the MQM report spatial outliers, often found at borders between spatially homogeneous areas or reporting histological structures that are too small to be adequately captured by the used neighborhood size. **e**, Ion image of *m/z* 12790.625 with a global Moran”s *I* of -0.003. **f**, SAM for *m/z* 12790.625. **g**, Moran scatterplot for *m/z* 12790.625 with *R*^2^ = 0.0001. **h**, MQM for *m/z* 12790.625.

The third case study demonstrates, on an MxIF-microscopy cohort imaging healthy and diseased human brain, how MQMs can be used to segment histological structures consistently across images with substantial inter-sample and intra-sample intensity variability. Vasculature architectures can inform on diseases such as cerebral amyloid angiopathy, a neuropathology seen in Alzheimer”s disease, characterized by amyloid plaque deposition in the walls of brain vasculature. In Figure 3, MQMs identify two important regions of human brain vasculature: the high-high quadrant (pink) delineates meningeal vessels within the frontal cortex, whereas the low-low quadrant (blue) reports perivascular space. Figure 3 demonstrates that the MQM can serve as an effective standalone tool for univariate image segmentation.

**Fig. 3.**
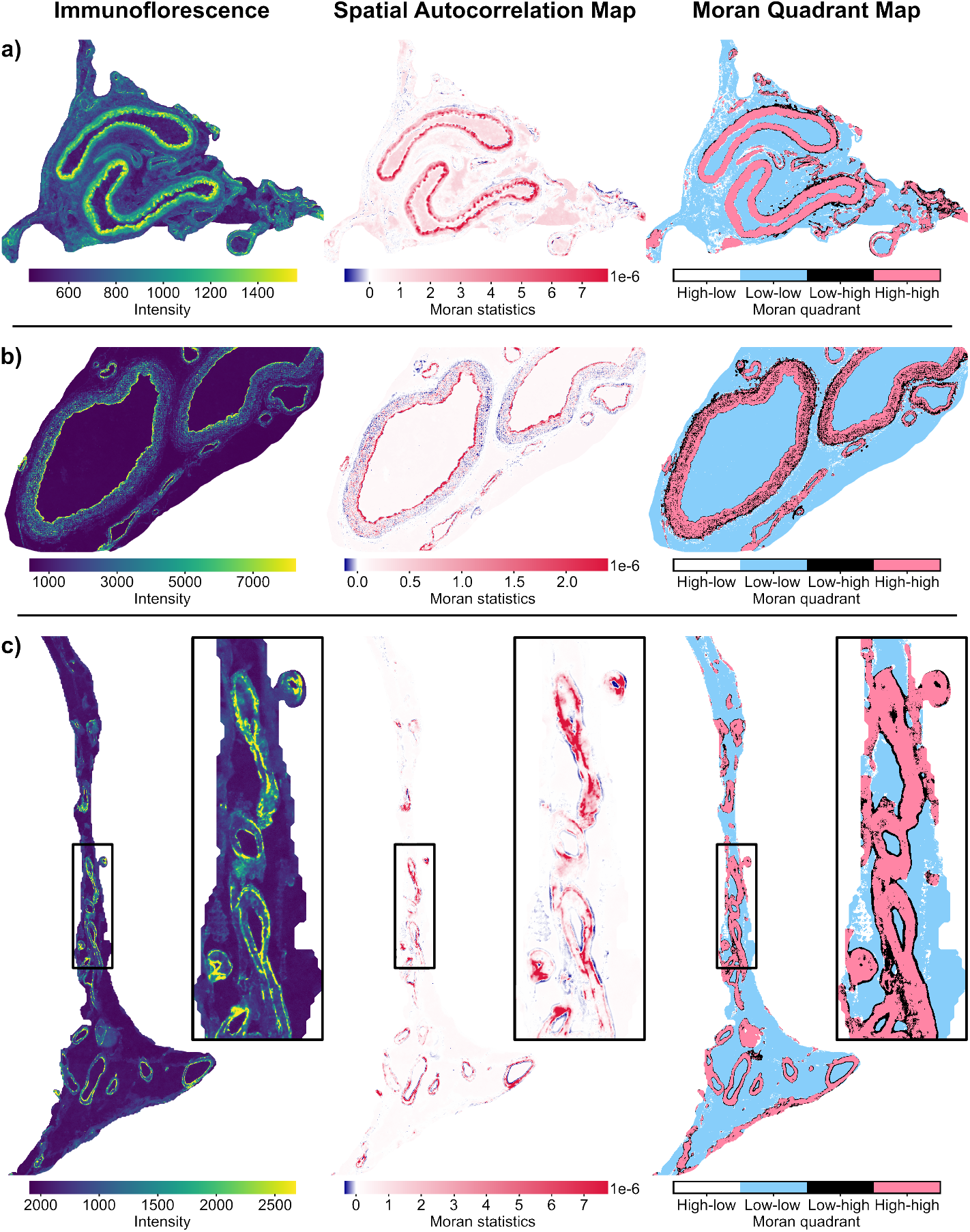
Spatial autocorrelation maps (SAMs) and Moran quadrant maps (MQMs) for three samples of human brain tissue imaged by MxIF-microscopy. Note that we consider only one channel of the MxIF datasets in our analysis. **a**, Sample acquired from a control subject. **b**, and **c**, Samples acquired from subjects suffering from Alzheimer”s disease, presenting severe cerebral amyloid angiopathy. Each sample was stained with a Collagen IV antibody marker post-MALDI-IMS imaging. The SAMs and MQMs of all three samples were obtained using a cumulative 10^th^-order Queen-contiguity spatial weights matrix, with no additional pre-processing. Delineating vasculature walls using only measured (photon) intensity, directly from the immunofluorescence images (left), is difficult with differences in absolute intensity response between experiments (compare **a** through **c**”s immunofluorescence images) and experiment-specific intensity perturbations (*e*.*g*., grid pattern in **b** due to MALDI-IMS measurements). While the SAMs” focus on spatial correlation amplifies certain borders and attenuates other intensity variations, they do not quite solve the delineation of vasculature walls. The MQMs, however, deliver (in pink) clear and consistent vasculature walls that are robustly characterized despite substantial differences in vasculature wall intensity variation between the experiments.

### 2.2 Moran-Felzenszwalb segmentation

The fourth case study demonstrates tissue domain segmentation on MALDI-IMS dataset n^*o*^2. It compares our novel MQM-based approach, namely Moran-Felzenszwalb segmentation (section 4.6), to state-of-the-art multivariate image segmentation approaches (Supplementary section 1.1): spatial *k*-means clustering algorithm [42] and the BANKSY algorithm [45]. Dataset n^*o*^2 measures a whole-body adult male zebrafish section at 10-*µm* pixel size. Figure 4a provides an autofluorescence image of the sample for reference, identifying anatomical regions such as the brain, eye, spinal cord, muscle, intestine, liver, gills, and heart. The aim of tissue domain segmentation is to partition the dataset into the aforementioned organs. However, the large size of dataset n^*o*^2, comprising 1,720,392 pixels (of which 1,005,524 on-tissue pixels) and 1,636 *m/z*-bins, poses a challenge. Indeed, many graph-based workflows developed for spatial transcriptomics do not scale to large IMS datasets due to their reliance on an adjacency matrix with quadratic space-complexity. For example, applying GraphST [43] to dataset n^*o*^2 would have required 3.68 TiB of RAM to store a 1,005,524 × 1,005,524 matrix. Our workflow addresses the need for efficient segmentation.

**Fig. 4.**
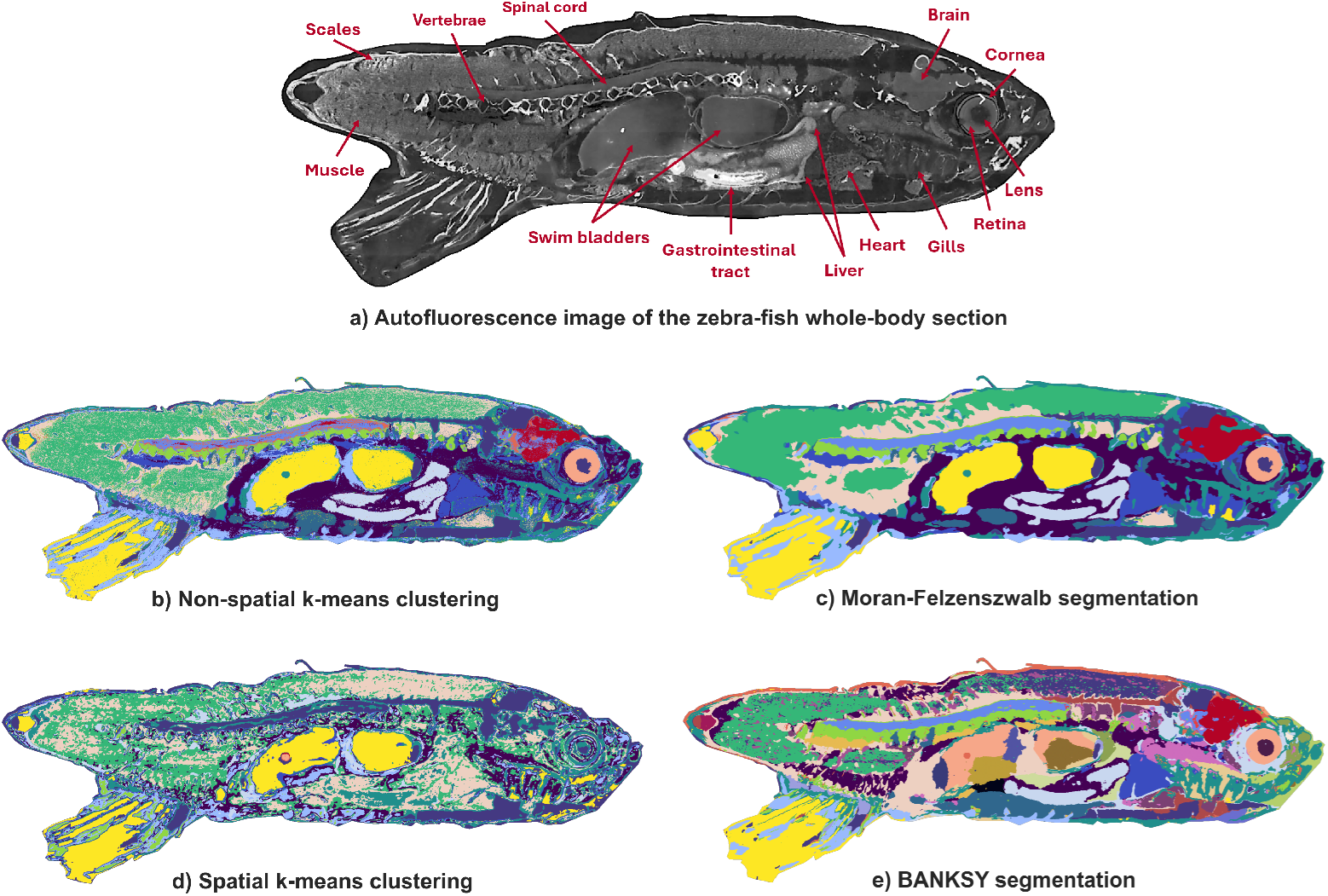
Tissue domain segmentation of MALDI-IMS dataset n^*o*^2. **a**, Autofluorescence image of the whole-body adult male zebrafish section. **b**, Results from (non-spatial) *k*-means clustering with *k* = 15. **c**, Results from our Moran-Felsenszwalb segmentation workflow: organ-delineating super-pixel boundaries are obtained by applying the Felsenszwalb segmentation to the Moran quadrant maps (MQMs) of the top-5 UMAP features, and clustering into 15 clusters. **d**, Results from spatial *k*-means clustering, using *k* = 15 clusters. **e**, Results from the BANKSY algorithm, which determined 41 clusters. Despite **d** and **e** coming from spatially-informed algorithms, the MQM-based results of Moran-Felsenszwalb segmentation are much less noisy than **d** and avoid the over-segmentation of **e**. Also, these workflows have following runtimes: 11.5 hours for BANKSY, 10.2 hours for spatial *k*-means clustering, and 13.7 minutes for Moran-Felsenszwalb segmentation, demonstrating that Moran-Felsenszwalb segmentation is both more scalable and more accurate than the other algorithms.

Figure 4 shows the results obtained by a spatially-agnostic method, non-spatial *k*-means clustering (Figure 4b), and three spatially-informed methods: our Moran-Felsenszwalb workflow (Figure 4c), spatial *k*-means clustering (Figure 4d), and BANKSY (Figure 4e). Our approach applies the Felsenszwalb segmentation algorithm to MQMs of the top-5 latent features provided by a non-linear dimensionality reduction algorithm, namely uniform manifold approximation and projection (UMAP) (Figure S11). The MQMs are computed using a cumulative 5^th^-order Queen-contiguity spatial weights matrix, resulting in a 5-pixel neighborhood size. For comparison, Figure 4b”s *k*-means clustering results with *k* = 15 are color-matched to Figure 4c”s Moran-Felsenszwalb results (Figure S12). Notably, Moran-Felsenszwalb segmentation delivers a less noisy, more homogeneous and interpretable partitioning of the zebrafish body, with muscle, vertebrae, spinal cord, brain, and heart as separate clusters (Figure S13). Moreover, the two swim bladders are assigned to the same cluster despite being spatially disjoint. The boundaries of the gastrointestinal tract and liver are correctly captured. In the eye, retina and cornea are distinguished. Despite spatial *k*-means clustering with *k* = 15, also using a 5-pixel-sized spatial neighborhood, its segmentation is far more noisy than Moran-Felsenszwalb segmentation. Given that BANKSY is based on the Leiden clustering algorithm, we cannot specify the number of clusters. Instead, we obtain 41 clusters and observe over-segmentation in the swim bladders and brain. Importantly, when segmenting dataset n^*o*^2, spatial *k*-means, BANKSY, and Moran-Felsenszwalb segmentation required 10.2 hours, 11.5 hours, and only 13.7 minutes, respectively. Therefore, we can conclude that Moran-Felsenszwalb segmentation is both more accurate and more scalable than the other algorithms.

The fifth case study performs tissue domain segmentation on MALDI-IMS dataset n^*o*^3, measuring a sagittal rat brain section at 20-*µ*m pixel size [51]. With 71,940 pixels and 92 *m/z*-bins, this dataset is far smaller than dataset n^*o*^2. However, its layered tissue with mixed cell populations makes its segmentation more difficult. Figure 5a shows both homogeneous and highly heterogeneous tissue regions. A spatially-agnostic partition into 9 clusters is obtained by non-spatial *k*-means clustering (Figure 5b). For Moran-Felzenszwalb segmentation, we compute MQMs for the top-12 UMAP features, using a cumulative 2^nd^-order Queen-contiguity spatial weights matrix (Figure S22), obtaining 9 clusters also. As per previous observations, Moran-Felzenszwalb segmentation (Figure 5c) is less noisy and discerns larger anatomical features, such as the midbrain, thalamus, and isocortex, as well as smaller structures within the hippocampus, differentiating the CA1-CA2-CA3 fields of Ammon”s horn, the pyramidal layer of CA3, and dentate gyrus layering. In comparison, spatial *k*-means segmentation (Figure 5d) is very detailed, but less interpretable, and BANKSY results (Figure 5e) are noticeably less detailed in the dentate gyrus. The spatially-informed workflows have the following runtimes: 12.8 minutes for spatial *k*-means clustering, 9.5 minutes for BANKSY, and only 1.3 minutes for Moran-Felsenszwalb segmentation.

**Fig. 5.**
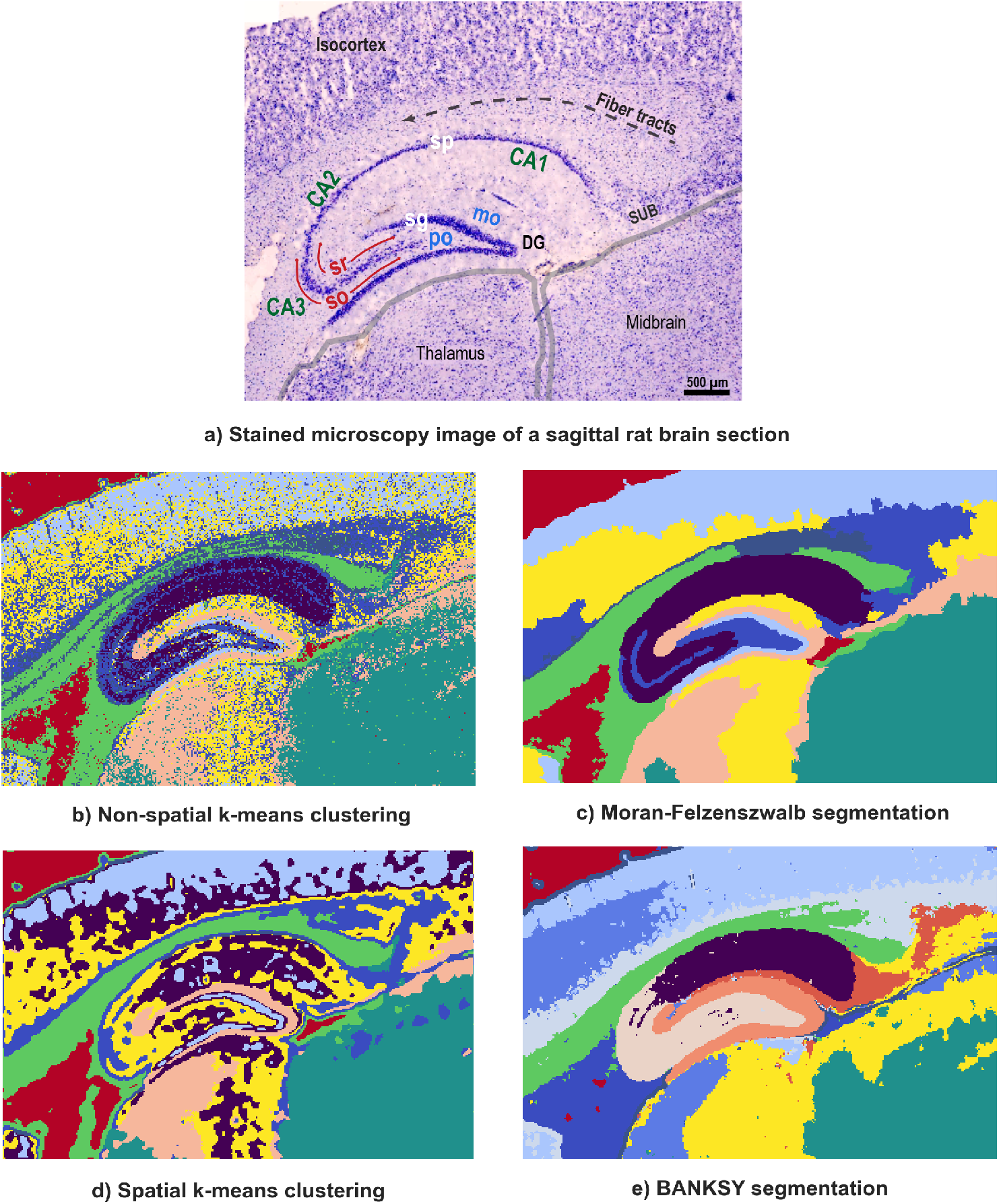
Tissue domain segmentation of MALDI-IMS dataset n^*o*^3. **a**, Stained microscopy image of the sagittal rat brain section, identifying anatomical regions within the hippocampus: Ammon”s horn (CA1, CA2, CA3), dentate gyrus (DG), the molecular layer (mo), polymorph layer (po), pyramidal layer (sp), granule cell layer (sg), stratum radiatum (sr), stratum oriens (so), and the subiculum (sub). **b**, Results from (non-spatial) *k*-means clustering with *k*=9. **c**, Results from our Moran-Felsenszwalb segmentation workflow: super-pixel boundaries are obtained by applying the Felsenszwalb segmentation to the Moran quadrant maps (MQMs) of the top-12 UMAP features, and clustering into 9 clusters. **d**, Results from spatial *k*-means clustering, using 9 clusters. **e**, Results from the BANKSY algorithm, which determined 15 clusters. The Moran-Felsenszwalb segmentation results are more interpretable than **d** and deliver more detailed layering within the dentate gyrus than **e**.

### 2.3 Moran-HOG clustering

The sixth case study demonstrates on MALDI-IMS dataset n^*o*^2 how MQMs can facilitate colocalization-based image clustering. IMS datasets often contain ion images that report similar spatial distributions at different *m/z*-values. For example, different *m/z*-bins may be co-expressed within specific tissue regions because they represent distinct forms of a given molecular species (*e*.*g*., isotopes, adducts, fragments), or correspond to different molecular species that share a precursor or biosynthetic pathway. Here, our novel Moran-HOG clustering workflow (section 4.6) is tested against two state-of-the-art self-supervised deep-clustering workflows (Supplementary section 1.2): NRDC [48] and DeepION [49]. Our workflow differs from NRDC and DeepION in that we do not use deep learning for spatial feature extraction. Rather, we use the Histogram-of-Oriented-Gradients (HOG) of the MQMs of the images. Table 1 compares the partitions obtained from all three workflows to manually-defined true labels using the adjusted Rand index (ARI), adjusted mutual information (AMI), balanced adjusted Rand index (bARI), and balanced V-measure [52].

**Table 1.**
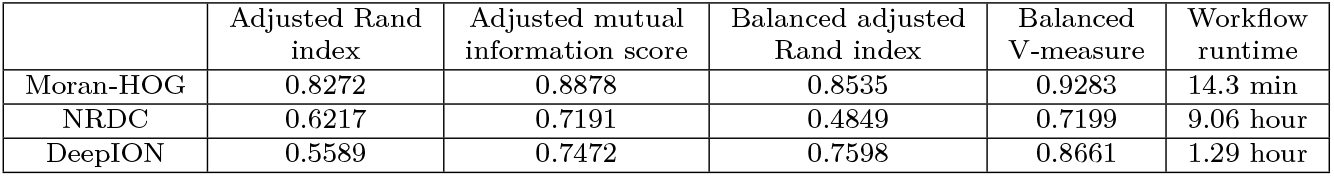
Clustering of 174 ion images from MALDI-IMS dataset n^*o*^ 2 into 8 groups of colocalized images. The performance and scalability of three colocalization-based clustering workflows is compared: Moran-HOG, NRDC, and DeepION. The clusters have varying sizes: the smallest cluster is made up of only 6 ion images, whereas the largest cluster is made up of 45 ion images. Some clusters are associated with a specific anatomical structure of the zebrafish; others are not. For all four metrics, larger values indicate better clustering. When comparing predicted clustering labels to true labels, the ARI and AMI equal 1 for identical partitions and approach 0 for random partitions. The bARI and the balanced V-measure are new clustering metrics that account for cluster size imbalance. The bARI is an extension of the ARI that weighs each cluster equally. The balanced V-measure is the harmonic mean of the balanced homogeneity and balanced completeness, both of which are entropy-based scores. The Moran-HOG clustering workflow outperforms the other two workflows across all clustering performance metrics as well as in terms of computational efficiency.

|  | Adjusted Rand index | Adjusted mutual information score | Balanced adjusted Rand index | Balanced V-measure | Workflow runtime |
| --- | --- | --- | --- | --- | --- |
| Moran-HOG | 0.8272 | 0.8878 | 0.8535 | 0.9283 | 14.3 min |
| NRDC | 0.6217 | 0.7191 | 0.4849 | 0.7199 | 9.06 hour |
| DeepION | 0.5589 | 0.7472 | 0.7598 | 0.8661 | 1.29 hour |

To establish a baseline, 174 ion images from dataset n^*o*^2 were manually assigned to 8 clusters (Figure S3). Moran-HOG colocalization analysis first obtains a MQM for each ion image, using a cumulative 5^th^- order Queen-contiguity spatial weights matrix. Then, we apply *k*-means clustering to the HOG of these MQMs. Figures S14-S21 illustrate how each image is characterized by its Moran-HOG features, also providing (non-MQM-based) HOG features for comparison. Table 1 demonstrates that, with ARI=0.8272, AMI=0.8878, bARI=0.8535, and balanced V-measure=0.9283, Moran-HOG clustering outperforms both NRDC and DeepION. Furthermore, Moran-HOG clustering is computationally efficient: its runtime is only 14.3 minutes, whereas other approaches require hours. To quantify the added-value of utilizing SAC, we also applied *k*-means clustering to HOG features extracted directly from the (scaled) images, rather than from their MQMs. This resulted in poor performance (ARI=0.482; AMI=0.166), demonstrating that MQMs facilitate HOG extraction of descriptive spatial features. These results suggest that deep learning does not necessarily outperform traditional statistical approaches on multiplexed molecular imaging data. Despite the AI community”s focus on scale (*e*.*g*., bigger models, bigger datasets, more compute), bigger is not necessarily better [53].

## 3 Discussion

This paper addresses the need for scalable analysis of the spatial-statistical properties of multiplexed molecular imaging data. It provides metrics and workflows that characterize but also exploit this information source to advance machine learning tasks and biological insight. We formalize the notion of spatial neighborhoods and demonstrate how local and global SAC statistics can be used to quantify spatial dependence and heterogeneity, which are two properties of molecular imaging data that violate the i.i.d. assumption. Furthermore, we propose a novel spatial feature extractor, the MQM, that can be used to inject spatial-statistical information into otherwise spatially-agnostic workflows. Our two MQM-based workflows, Moran-Felsenszwalb segmentation and Moran-HOG clustering, outperform state-of-the-art approaches on tissue domain segmentation and colocalization-based clustering case studies. Computational efficiency is central to the development and implementation of our methods, and our results can be reproduced using the open-source Moran Imaging package supplied. Although demonstrated on MALDI-IMS and MxIF-microscopy, our methods are equally applicable to other spatial-omics and molecular imaging/profiling modalities such as IMC and CODEX-microscopy, and potentially also in other areas of computer vision. Besides facilitating spatial-statistical analyses for large tissue atlasing initiatives, such as HuBMAP, KPMP, and HTAN, this paper”s methodological contributions and toolbox help unlock spatial statistics for large-scale molecular imaging, hence advancing spatial biology.

## 4 Methods

Consider one image from a multiplexed imaging measurement, for example, an ion image from an IMS dataset. For our analysis, each image is a rasterized rectangular grid of *n* pixels, and we flatten the image into a vector of pixel values ***z*** ∈ ℝ^*n×*1^. In Equation 1, mean-centering ***z*** yields a vector ***x*** ∈ ℝ^*n×*1^, where entry *x*_*i*_ reports the intensity value *z*_*i*_ observed at the *i*^th^ pixel minus the mean intensity *µ*_*z*_ of the whole image.

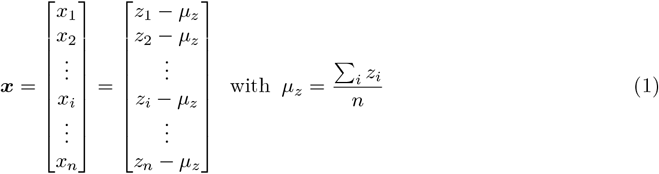

### 4.1 Spatial weights matrix

Defining spatial relationships between pixels is an essential first step towards quantifying spatial dependence and heterogeneity. Since most molecular imaging modalities deliver rasterized images, we employ contiguity-based weighting to define these spatial relationships. Contiguity criteria are generally defined in analogy to the game of chess [54, 55]: Rook-contiguity, which requires neighboring pixels to have a common border; Bishop-contiguity, which requires neighboring pixels to have a common vertex; and Queen-contiguity, which requires neighboring pixels to have a common border and/or vertex. In our work, we employ the Queen-contiguity criterion. We use a spatial weights matrix to formalize the notion of pixel neighborhood and to capture the spatial relationships present in a given image. Initially, we define a binary spatial weights matrix ***C*** [54–56] as a square, sparse, and symmetric matrix that has one spatial weight *c*_*i,j*_ per pair of pixels (Equation 2). For two different pixels (*i* ≠ *j*), *c*_*i,j*_ = 1 if pixels *i* and *j* are neighbors and *c*_*i,j*_ = 0 otherwise. By convention, spatial weights on the diagonal of ***C*** are zero (*c*_*i,i*_ = 0) because a pixel is not considered a neighbor of itself.

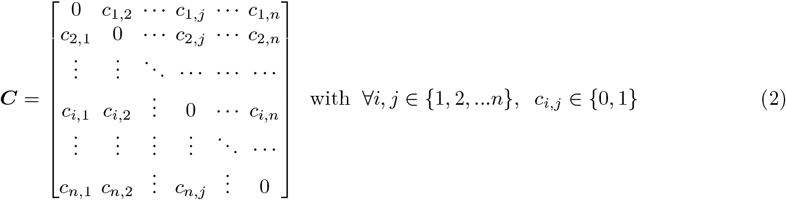

Although each ion image is modeled as a rectangular grid of pixels, the tissue sample under study is not necessarily rectangular. The ion images of an arbitrarily-shaped sample often consists of many off-tissue pixels, which do not encode biological information and should therefore be omitted from our analysis. We exclude off-tissue pixels by assigning them zero neighbors. For each off-tissue pixel *u*, we set the elements of the corresponding row (and column) of the spatial weights matrix ***C*** to zero: ∀*j, c*_*u,j*_ = 0. A pixel without neighbors is called an isolate [55, 56]. Note that explicitly accounting for isolates is uncommon in traditional (geo)spatial statistics literature and software, yet it is important for molecular imaging modalities that produce rasterized data, such as IMS and MxIF-microscopy. The spatial weights matrix ***C*** encodes the spatial connectivity of adjacent pixels and is referred to as a 1^st^-order contiguity matrix. To quantify the spatial dependence between a given pixel and the neighbors of its neighbors, we need a 2^nd^-order contiguity matrix. Assuming a binary and symmetric first-order contiguity matrix ***C***, we define the spatial weights matrix of order *k* as ***C***^*k*^ [54]. For example, ***C***^3^ is a 3^rd^-order spatial weights matrix. There is a difference between pure higher-order contiguity, which excludes lower-order neighbors, and cumulative higher-order contiguity, which includes lower-order neighbors (Figure S1). Section 2.1 in the Supplementary Information provides more details on *n*^th^-order contiguity matrices.

To facilitate interpretation and inter-experiment comparison, row-standardizing the spatial weights matrix is necessary. Row-standardization is the row-wise scaling of the spatial weights as per Equation 3 [54, 55]. Row-standardized weights *w*_*i,j*_ are obtained by dividing the 1^st^-order binary weights *c*_*i,j*_ (or the *k*^th^-order binary weights 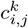) by their row sum. Each row-standardized weight is between 0 and 1, and they sum to one across each row (∑_*j*_ *w*_*i,j*_ = 1). The sum of all weights, *S*_0_ = ∑_*i*_ ∑_*j*_ *w*_*i,j*_, is equal to (*n* − *q*) where *n* is the total number of pixels and *q* is the number of isolates [55, 56]. Note that the row-standardized spatial weights matrix, denoted ***W***, is no longer binary nor symmetric.

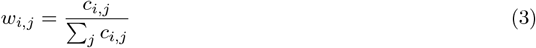

In addition to the observed value *x*_*i*_ at pixel *i*, we now introduce a spatial lag value for pixel *i*. The spatial lag, denoted **L**_***W***_ *x*_*i*_, is the weighted mean of the values observed at its neighboring pixels, not including *x*_*i*_. It is obtained by the spatial lag operation in Equation 4, which averages the observations (*e*.*g*., ion intensity measurements) at neighboring locations (*i*.*e*., pixels) into a value referred to as a spatially lagged variable [54, 57]. The spatial weights of pixel *i* determine which pixels belong to its neighborhood set 𝒩_*i*_, and weigh the contribution of each neighboring pixel”s observation *x*_*j*_ to the spatial lag value **L**_***W***_ *x*_*i*_ of pixel *i*. In matrix notation, the spatial lag expression corresponds to the matrix product of the *n* × *n* matrix ***W*** with the *n* × 1 vector of observations ***x***. The spatial weights matrix ***W*** can therefore be considered as a spatial lag operator applied to the flattened and mean-centered image ***x***. The spatially lagged variable **L**_***W***_ *x*_*i*_ is expressed in the same unit as the variable *x*_*i*_ under study, which facilitates interpretation. In section 4.5, we propose a more efficient approach to computing the spatial lag using convolution rather than matrix multiplication.

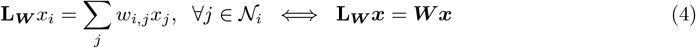

Row-standardization provides convenient bounds on the eigenvalues of the spatial weights matrix ***W***. The eigenvalues of the binary spatial weights matrix ***C*** do not fall within a set range, which is problematic because, as discussed in Supplementary section 2.3, the extreme eigenvalues of the spatial weights matrix determine the range of Moran”s *I* statistic [58–60]. If we were to use ***C***, we would therefore only be able to compare two Moran”s *I* statistics if they were computed using the same spatial weights matrix [20]. However, in molecular imaging, we need to compare the Moran”s *I* statistics obtained from different tissue samples, with distinct geometries that necessitate different spatial weights matrices. The largest eigenvalue of a row-standardized spatial weights matrix ***W*** is *λ*_1_ = +1 and the smallest eigenvalue satisfies −1 ≤*λ*_*n*_ ≤ − 0.5 [61]. Indeed, the Perron-Frobenius theorem states that the largest eigenvalue of a real, square, and non-negative matrix is contained in the interval defined by the minimum and maximum row sum, and that the sum of the eigenvalues equals the trace of the matrix [61]. Regarding matrix ***W***, its rows sum to one and it has a trace of zero.

### 4.2 Global Moran”s *I*

Moran”s *I*, defined in Equation 5, is a commonly used univariate measure of global SAC [19, 20]. It measures the coincidence of attribute similarity (*i*.*e*., content similarity) with locational similarity across an entire image. A positive Moran”s *I* indicates clustering (*i*.*e*., spatial proximity of similar values), a Moran”s *I* close to zero indicates spatial randomness, and a negative Moran”s *I* indicates dispersion (*i*.*e*., spatial proximity of dissimilar values) [17, 18, 20]. We refer to a spatially random pattern as noise. In Equation 5, the numerator is a measure of spatial autocovariance and the denominator is a measure of (general) variance. *S*_0_ is the sum of the weights of the row-standardized spatial weights matrix ***W***. Given that *n* is the total number of pixels and *q* is the number of off-tissue pixels or isolates, (*n* − *q*) is the number of pixels for which we have measurements.

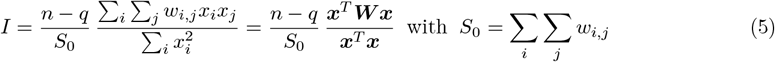

The sum *S*_0_ of all spatial weights is equal to (*n* − *q*) because ***W*** is row-standardized [20, 55]. Equation 5 therefore simplifies to Equation 6: Moran”s *I* is obtained by standardizing the spatial autocovariance. In other words, Moran”s *I* is formally equivalent to the regression coefficient in a linear least squares regression of the spatial lag ***W x*** on the mean-centered variable ***x*** [32, 54, 55].

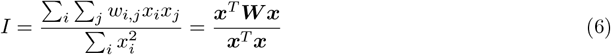

The expected value of Moran”s *I* under the null hypothesis of spatial randomness is given by Equation 7 [17, 58]. In images with a large number of pixels, Equation 7 simplifies to 𝔼 [*I*] ≈ 0. Note that although Moran”s *I* is defined similarly to the Pearson correlation coefficient, it is not necessarily bounded between − 1 and +1. As demonstrated in section 2.3 of the Supplementary Information, the upper and lower bounds of Moran”s *I* statistic are determined by the extreme eigenvalues of the spatial weights matrix [58–60]. However, in the case of a row-standardized spatial weights matrix, the range of Moran”s *I* statistic is [−1, +1].

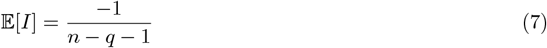

As demonstrated in section 2.1, Moran”s *I* can be used to reject the null hypothesis of spatial randomness in an image in favor of an alternative hypothesis of positive global SAC. We, therefore, need to determine whether the Moran”s *I* statistic computed for a given image is significantly larger than its 𝔼 [*I*] (*i*.*e*., *I >* E(*I*)). We choose a non-parametric hypothesis testing method called permutation testing [20, 62]. The idea of permutation testing is to compare the Moran”s *I* statistic of an image to a distribution of Moran”s *I* statistics that are obtained from spatially randomized versions of that image. The distribution of Moran”s *I* statistics under the null hypothesis, called the reference distribution, is obtained by randomly permuting the observed (ion) intensity values across spatial locations and recording the Moran”s *I* statistic corresponding to each permutation. The statistical significance of the Moran”s *I* of the measured (non-randomized) image is established by comparing its value to the reference distribution. The reference distribution is used to compute a pseudo-*p*-value, which is defined in Equation 8. The pseudo-*p*-value is equal to the fraction of statistics in the reference distribution that are equal to or more extreme than the observed Moran”s *I* statistic. It is a sampling-based approximation of the probability that the observed Moran”s *I* statistic could have been obtained if the null hypothesis were true. In Equation 8, *R* is the number of times that the statistic obtained from a randomized image is as extreme or more extreme than the Moran”s *I* statistic of the measured image, and *M* is the total number of permutations. A large number of permutations is recommended to obtain a robust estimate of the reference distribution and a reliable pseudo-*p*-value. In our case studies, we set *M* = 999, and as a result 0.001 is the minimum pseudo-*p*-value: *p* = 0.001 is obtained if none of the statistics from the randomized reference distribution exceed the Moran”s *I* of the measured (ion) image. If the pseudo-*p*-value is smaller than a user-determined threshold *α*, the null hypothesis is rejected, and the observed Moran”s *I* statistic is declared statistically significant. We set *α* to double the minimum pseudo-*p*-value (*α* = 0.002) to minimize the risk of false positives [63]. In conclusion, assuming statistical significance, a positive Moran”s *I* indicates positive global SAC.

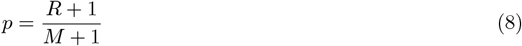

### 4.3 Local Moran statistics & Spatial Autocorrelation Map

Moran statistics are local SAC statistics developed by Anselin to enable a localized study of spatial patterns [28, 64]. As defined in Equation 9, the Moran statistic *I*_*i*_ of a pixel *i* measures the SAC within the pixel”s neighborhood 𝒩_*i*_, as specified by the *i*^th^ row of the row-standardized spatial weights matrix ***W***. In the numerator of Equation 9, the spatial lag **L**_***W***_ *x*_*i*_ = ∑_*j*_ *w*_*i,j*_*x*_*j*_ is the weighted mean of neighboring pixels” (ion) intensity values defined in Equation 4. In the denominator, the sum of all squared (ion) intensity values, 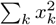, is a constant for all pixels.

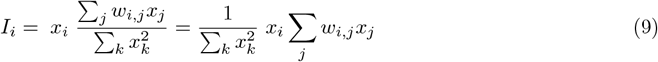

Moran statistics enable the localization of spatial clusters and outliers within an image [28, 64]. A positive Moran statistic *I*_*i*_ indicates positive local SAC for pixel *i*, indicating that pixel *i* is surrounded by similar valued observations. A Moran statistic near zero indicates spatial randomness within the pixel”s neighborhood. A negative Moran statistic *I*_*i*_ indicates negative local SAC, indicating that pixel *i* is an outlier with respect to its neighborhood. Given that we have one Moran statistic per pixel, these statistics can be visualized as an image. We refer to the visualization obtained by mapping the Moran statistics across a tissue sample as a spatial autocorrelation map (SAM). In multiplexed molecular imaging, the SAM of a molecular species can be viewed as a SAC-based image filter whose purpose is to facilitate the detailed characterization of local patterns of spatial association. As demonstrated by Equation 10, the local Moran statistics sum to the global Moran”s *I* from Equation 6 [28, 64]. The relative magnitude of a pixel”s Moran statistic is, therefore, indicative of its degree of influence on the image”s global spatial dependency.

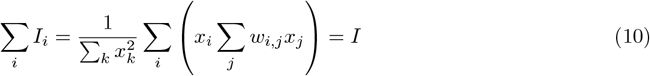

The significance of local Moran statistics can be assessed using the conditional permutation approach proposed by Anselin [28, 64, 65]. Similarly to the spatial permutation approach used to assess the significance of global Moran”s *I*, a reference distribution of local SAC statistics under the null hypothesis can be computed and the measured SAC statistic can be compared to the reference distribution using a pseudo-*p*-value. However, one difference is that we perform only one permutation test per pixel to simulate spatial randomness within the neighborhood of that pixel. For a given local Moran statistic *I*_*i*_, the conditional permutation consists of randomly relocating the pixel values within the neighborhood 𝒩_*i*_, while fixing the value *x*_*i*_ of pixel *i*. The significance of *I*_*i*_ could be assessed using the pseudo-*p*-value of pixel *i*. In practice, however, this would be unreliable because local pseudo-*p*-values suffer from multiple comparison issues and the potentially biasing effect of positive global SAC [64, 65]. In our case studies, many thousands of hypothesis tests (one per pixel) would need to be performed on overlapping subsets of pixels (*i*.*e*., the same data would be used in multiple tests). In the presence of positive global SAC, the local Moran statistics obtained from overlapping pixel neighborhoods would be correlated and, therefore, the tests would not be independent [64, 66]. Standard multiple comparison correction methods, such as the Bonferroni correction, would therefore result in potentially misleading claims about local clustering in molecular imaging data. Although these issues have been extensively discussed in the spatial statistics literature [64, 65, 67], we argue that none of the proposed solutions are effective on molecular imaging data. To avoid making potentially misleading claims about local clustering on the basis of pseudo-*p*-values that cannot provide a reliable bound on false positives, we choose not to report the significance of local Moran statistics. Instead, in this work, we propose using local Moran statistics in a descriptive manner.

### 4.4 Moran scatterplot & Moran quadrant map

After computing the SAM of an image, the spatial association of each pixel with its neighbors can be further classified into one of four SAC categories using the Moran scatterplot introduced by Anselin [20, 32]. It involves plotting each pixel”s spatial lag **L**_***W***_ *x*_*i*_ versus its observed mean-centered (ion) intensity *x*_*i*_. The Moran scatterplot is divided into four quadrants that represent four different types of local SAC (Figure S2). Since the plot is centered on the means of the observed (ion) intensities and spatial lag values, we can refer to an (ion) intensity value as high or low depending on whether the (*x*_*i*_, **L**_***W***_ *x*_*i*_)-pair is to the left or to the right of the vertical axis respectively, and we can refer to a spatial lag value as high or low depending on whether the (*x*_*i*_, **L**_***W***_ *x*_*i*_)-pair is above or below the horizontal axis respectively [20, 32, 66]. The upper-right and lower-left quadrants correspond to positive local SAC (*I*_*i*_ *>* 0): high (ion) intensity values that cluster together manifest in the upper-right (high-high) quadrant, and low (ion) intensity values that cluster together end up in the lower-left (low-low) quadrant. The lower-right and upper-left quadrants correspond to negative local SAC (*I*_*i*_ *<* 0): spatial outlier pixels that have a higher (ion) intensity than their neighbors are in the lower-right (high-low) quadrant, whereas spatial outliers that have a lower (ion) intensity than their neighbors are in the upper-left (low-high) quadrant. In this work, we expand on Anselin”s approach by mapping the classification of pixels onto one of the four Moran scatterplot quadrants. We refer to this spatial representation of Moran scatterplot quadrant membership as a Moran quadrant map (MQM). The MQM assigns each pixel to a specific Moran scatterplot category. In multiplexed molecular imaging, a unique MQM can be extracted for each molecular species that is measured. In our case studies, we show how a MQM can be used for direct human interpretation of a species” distribution (*e*.*g*., exhibiting less sensitivity to noise than the measured intensity pattern). We also demonstrate how the MQM can serve as a means of introducing SAC-awareness into an otherwise spatially-agnostic computer vision workflow.

The Moran scatterplot can also be used to assess the degree of spatial heterogeneity present in an image. If we perform a linear regression of the form ***y*** = *α* + *β****x*** where ***y*** = ***W x***, the linear least-squares minimization problem 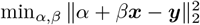 is solved by setting the gradient of the cost function with respect to *α* and *β* to zero. The least-squares estimates of the intercept *α* and slope *β* are given by Equation 11.

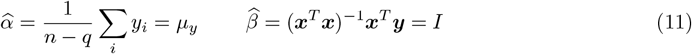

As seen by comparing Equations 6 and 11, the slope of the linear regression of the spatial lag (*y*-axis) on the mean-centered (ion) intensity (*x*-axis) is equal to Moran”s *I*. Therefore, the more the linear regression line reflects the global pattern of association between ***x*** and ***y***, the higher the degree of spatial stationarity [32]. Although this assessment is typically performed through visual inspection of the Moran scatterplot, such an approach is inherently subjective and not scalable to large sets of (ion) images. We therefore propose a more rigorous method based on the coefficient of determination *R*^2^. In Equation 12, *R*^2^ is defined as one minus the residual sum of squares divided by the total sum of squares and verifies 0 ≤ *R*^2^ ≤ 1. According to Equation 4, *y*_*i*_ = **L**_***W***_ *x*_*i*_ = ∑_*j*_ *w*_*i,j*_*x*_*j*_, the coefficient of determination can be interpreted as follows: a high *R*^2^ indicates that there is little deviation from the least squares regression line and that Moran”s *I* captures most of the spatial variance. In section 2.1, we report spatial stationarity for an image if *R*^2^ ≈ 1, and spatial heterogeneity if *R*^2^ ≈ 0. No prior work, to our knowledge, has used the coefficient of determination of the Moran scatterplot to quantify spatial heterogeneity.

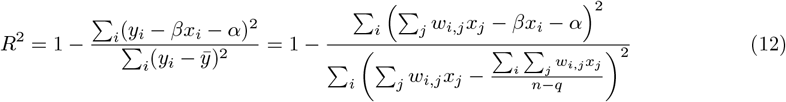

### 4.5 Novel parallelized spatial lag computation

Efficient spatial lag computation is critical for using spatial statistics in the context of molecular imaging, where datasets usually consist of thousands to millions of pixels. Although the traditional approach to spatial lag calculation described in section 4.2 works for any spatial arrangement of measurements, it proves insufficiently scalable in terms of runtime and memory requirements. We introduce a new approach that leverages the rasterized structure of molecular imaging data. Instead of computing spatial lag by matrix multiplication, we develop here a convolutional approach that facilitates parallel computation (*e*.*g*., via a graphics processing unit) and which is, therefore, scalable to very large imaging datasets. Our novel parallelized spatial-lag implementation is key for applying our Moran-Felsenszwalb and Moran-HOG workflows, which are presented in section 4.6, to MALDI-IMS and MxIF-microscopy datasets with thousands to millions of pixels.

Consider the 2-D mean-centered rasterized image *X*, obtained by reshaping the vector ***x*** in Equation 1 to the size *n*_row_ × *n*_col_ of the original image. With *i* ∈ {1, 2, …, *n*_row_ } and *j* ∈ {1, 2, …, *n*_col_ }, the meancentered (ion) intensity of the pixel with spatial coordinates (*i, j*) is denoted *x*_*i,j*_. Assuming row-major ordering, the mapping from *x*_*i,j*_ to *x*_*k*_ is the following: *k* = *I* · *n*_col_ + *j*. In analogy to the binary spatial weights matrix ***C***, we define the binary spatial weights kernel ***k***_***C***_ corresponding to a 1^st^-order Queen-contiguity neighborhood in Equation 13. What follows is an example of how to compute the spatial lag of a pixel using convolutional operations. We could use other contiguity criteria and/or define higher-order neighborhoods as well.

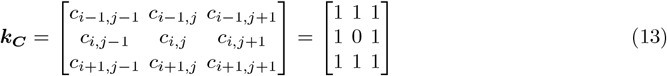

The row-standardized spatial weights kernel ***k***_***W***_ follows from ***k***_***C***_ in Equation 13. Equation 14 performs row-standardization, as defined in Equation 3, for a pixel whose neighborhood set consists of 8 pixels. When using a 1^st^-order Queen-contiguity neighborhood criterion, all on-tissue non-border pixels have 8 neighbors.

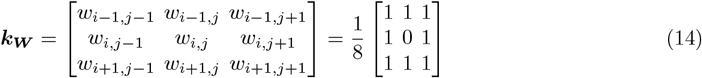

Equation 15 defines the patch ***P***_***i***,***j***_ of rasterized data surrounding the pixel with spatial coordinates (*i, j*).

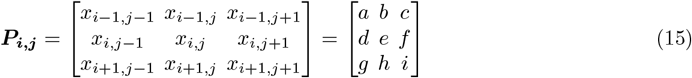

The spatial lag for pixel (*i, j*) can then be computed via convolution as described in Equation 16.

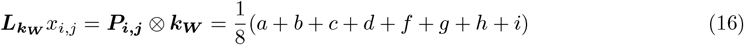

When a pixel is close to the border of the tissue sample under study (*e*.*g*., on the acquisition mask border), the convolutional approach described above risks accounting for off-tissue pixels. Therefore, the rasterized image is padded with zeros. In the example of Equation 17, pixel (*i, j*) is located in the top-left corner of the ion image and only has three neighbors (rather than eight). The row-standardization factor of the spatial weights kernel ***k***_***W***_ is adjusted accordingly as per Equation 17.

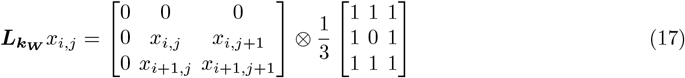

### 4.6 Novel segmentation & clustering workflows

In sections 2.2 and 2.3, we demonstrate how spatial statistics can be captured using the MQM as feature extractor, and how it can be provided to a spatially-agnostic computer vision method as an input. We target two important use cases in molecular imaging to deliver proofs-of-concept, namely tissue domain segmentation and colocalization-based image clustering. A brief description of these two unsupervised workflows is provided below.

#### Moran-Felzenszwalb tissue domain segmentation

Our first workflow focuses on the unsupervised segmentation of a tissue section into tissue domains (*e*.*g*., cell types, multicellular functional tissue units, anatomical regions). The Moran-Felzenszwalb workflow starts by extracting the localized SAC information of a set of latent features using MQMs. This demonstrates that the proposed feature extractor is also applicable to latent features derived through dimensionality reduction or feature engineering. Our workflow performs multivariate semantic image segmentation by providing the MQMs to the Felsenszwalb algorithm, which is a computationally efficient graph-based image segmentation algorithm [68]. The result is a partition of the dataset into a set of super-pixels, which are spatially contiguous and molecularly homogenous regions, that are assigned to a biological class on the basis of prior information or their molecular signature. The Moran-Felzenszwalb workflow consists of the following steps: (1) perform dimensionality reduction of the dataset (*e*.*g*., with UMAP); (2) define a spatial contiguity matrix and compute the MQMs of the latent features, using our Moran Imaging Toolbox; (3) partition the image into super-pixels by applying the Felzenszwalb segmentation algorithm to the MQMs; (4) perform pixel-level clustering using the (non-spatial) *k*-means or Leiden algorithm; (5) label each super-pixel obtained from step 3 according to the mode of the class labels obtained from step 4 of the pixels making up that super-pixel. The results depend on the following hyperparameters: the number of latent features to compute in step 1; the contiguity criterion and neighborhood order of the spatial weights matrix defined in step 2; the hyperparameter that determines the degree of latent feature map smoothing prior to Felzenszwalb segmentation and the two hyperparameters that determine the size of the super-pixels in step 3; and the hyperparameters of the pixel-level clustering algorithm used in step 4. The performance of our Moran-Felzenszwalb workflow is sensitive to the hyperparameters of the Felzenszwalb segmentation algorithm, in step 3, and to the hyperparameters of the pixel-level clustering algorithm, in step 4. It is robust with respect to the other hyperparameters.

#### Moran-HOG colocalization-based clustering

Our second workflow is the Moran-HOG clustering method for clustering of ion images with similar spatial patterns. It combines the MQM with a traditional feature descriptor from computer vision: the Histogram-of-Oriented-Gradients (HOG). HOG is a gradient-based local feature extraction method [69]. In this workflow, the MQM is applied directly to the ion images. The Moran-HOG workflow consists of the following steps: (1) define a spatial contiguity matrix and compute the MQMs for all ion images; (2) compute the HOG features from the MQMs; (3) perform image-level clustering of the mean-centered and unit-variance-scaled HOG features using the (non-spatial) *k*-means clustering algorithm. The results depend on the following hyperparameters: the contiguity criterion and neighborhood order of the spatial weights matrix defined in step 1; the hyper-parameters of the HOG feature descriptor (*e*.*g*., number of orientation bins, number of pixels per cell, number of cells per block) in step 2; and the number of clusters for the *k*-means clustering algorithm in step 3. Our Moran-HOG workflow is robust across a wide range of hyperparameter settings. Our work-flow”s modular design allows for flexibility. For example, if the number of clusters is not known a priori, the *k*-means clustering algorithm could easily be replaced by another algorithm, such as a density-based clustering algorithm (*e*.*g*., the HDBSCAN algorithm) or a graph-based clustering algorithm (*e*.*g*., the Leiden algorithm).

Note that, when computing MQMs in the context of the Moran-Felzenszwalb and Moran-HOG workflows, we use the efficient convolution-based approach to spatial lag computation presented in section 4.5.

### 4.7 Case study overview

**Table 2.** Overview of the different case studies.

| Case study | Used in section | Used for | Dataset | Tissue type | Molecular class | Figures |
| --- | --- | --- | --- | --- | --- | --- |
| 1 | 2.1 | global SAC quantification | MALDI-IMS dataset n°1 | rat brain (coronal section) | proteins | 1, S8 |
| 2 | 2.1 | local SAC quantification & SAM and MQM feature extractors | MALDI-IMS dataset n°1 | rat brain (coronal section) | proteins | 2, S9-S10 |
| 3 | 2.1 | SAM and MQM feature extractors for biological delineation | MxIF-microscopy dataset | (healthy & diseased) human brain | proteins | 3 |
| 4 | 2.2 | tissue domain segmentation | MALDI-IMS dataset n°2 | whole-body zebrafish | lipids | 4, S11-S13 |
| 5 | 2.2 | tissue domain segmentation | MALDI-IMS dataset n°3 | rat brain (sagittal section) | lipids | 5, S22-S27 |
| 6 | 2.3 | colocalization clustering | MALDI-IMS dataset n°2 | whole-body zebrafish | lipids | Table 1, S14-S21 |

## Nomenclature

*m/z*: mass-to-charge ratio
*AMI*: adjusted mutual information
*ARI*: adjusted Rand index
*bARI*: balanced adjusted Rand index
*CODEX*: co-detection by indexing
*HOG*: Histogram-of-Oriented-Gradients
*HTAN*: Human Tumor Atlas Network
*HuBMAP*: Human Biomolecular Atlas Program
*i.i.d*.: independent and identically distributed
*IMC*: imaging mass cytometry
*IMS*: imaging mass spectrometry
*KPMP*: Kidney Precision Medicine Project
*MALDI*: matrix-assisted laser desorption/ionization
*MERFISH*: multiplexed error-robust fluorescence in situ hybridization
*MIBI*: multiplexed ion beam imaging
*MQM*: Moran quadrant map
*MxIF*: multiplexed immunofluorescence
*NRDC*: noise-robust deep-clustering
*SAC*: spatial autocorrelation
*SAM*: spatial autocorrelation map
*UMAP*: uniform manifold approximation and projection

## Acknowledgments

Research reported in this publication was supported by the National Institutes of Health (NIH)”s Common Fund, National Institute Of Diabetes And Digestive And Kidney Diseases (NIDDK), and the Office Of The Director (OD) under Award Numbers U54DK120058, U54DK134302, and U01DK133766 (J.M.S. and R.V.), by NIH”s Common Fund, National Eye Institute, and the Office Of The Director (OD) under Award Number U54EY032442 (J.M.S. and R.V.), by NIH”s National Institute Of Allergy And Infectious Diseases (NIAID) under Award Numbers R01AI138581 and R01AI145992 (J.M.S., E.P.S., and R.V.), by NIH”s National Institute On Aging (NIA) under Award Number R01AG078803 (J.M.S., M.S.S., and R.V.), by NIH”s National Institute of Neurological Disorders and Stroke (NINDS) under Award Numbers 1RF1NS130334 and 1RF1NS129735 (M.S.S.), by NIH”s National Cancer Institute (NCI) under Award Number U01CA294527 (J.M.S. and R.V.), and by the National Science Foundation Major Research Instrument Program CBET - 1828299 (J.M.S.). The research was furthermore made possible in part by grant numbers 2021-240339 and 2022-309518 (L.G.M. and R.V.) from the Chan Zuckerberg Initiative DAF, an advised fund of Silicon Valley Community Foundation. The content is solely the responsibility of the authors and does not necessarily represent the official views of the National Institutes of Health.

## Notes

https://github.com/vandeplaslab/Moran_Imaging/

https://zenodo.org/records/17399931

http://doi.org/10.35079/HBM252.SRFF.799

### Competing Interest Statement

The authors have declared no competing interest.

