## Supplementary material for "Spatial Dependence and Heterogeneity in Molecular Imaging: Moran Quadrant Maps Enable Advanced Spatial-Statistical Analysis"

Supplementary Information
for

### Spatial Dependence and Heterogeneity in Molecular Imaging: Moran Quadrant Maps Enable Advanced Spatial-Statistical Analysis

Léonore E. M. Tideman[1], Felipe A. Moser[1], Lukasz G. Migas[1], Jacquelyn Spathies[2,3], Katerina V. Djambazova[2,4], Cody R. Marshall[2,5], Matthew S. Schrag[6,7], Eric P. Skaar[3], Jeffrey M. Spraggins[2,3,4,5,8,9], Raf Van de Plas[1,2,8*]

[1]Delft Center for Systems and Control, Delft University of Technology, Delft 2628 CD, Netherlands.
[2]Mass Spectrometry Research Center, Vanderbilt University, Nashville, TN 37232, USA.
[3]Department of Pathology, Microbiology, and Immunology, Vanderbilt University Medical Center, Nashville, TN 37232, USA.
[4]Department of Cell and Developmental Biology, Vanderbilt University, Nashville, TN 37232, USA.
[5]Chemical and Physical Biology Program, Vanderbilt University, Nashville, TN 37232, USA.
[6]Department of Neurology, Vanderbilt University Medical Center, Nashville, TN 37232, USA.
[7]Vanderbilt Brain Institute, Vanderbilt University, Nashville, TN 37232, USA.
[8]Department of Biochemistry, Vanderbilt University, Nashville, TN 37232, USA.
[9]Department of Chemistry, Vanderbilt University, Nashville, TN 37232, USA.

### Nomenclature

*m/z* mass-to-charge ratio
*FT-ICR* Fourier-transform ion cyclotron resonance
*HOG* Histogram-of-Oriented-Gradients
*HuBMAP* Human Biomolecular Atlas Program
*IMS* imaging mass spectrometry
*MALDI* matrix-assisted laser desorption/ionization
*MESF* Moran eigenvector spatial filtering
*MQM* Moran quadrant map
*MxIF* multiplexed immunofluorescence
*SAC* spatial autocorrelation
*TIMS* trapped ion mobility spectrometry

### 1 Supplementary background

Over the past decade, tailored machine learning workflows have been developed for the purpose of exploiting both the spatial and molecular information encoded in multiplexed molecular imaging data. In this supplementary section, we provide additional background for two molecular imaging data analysis tasks for which spatio-molecular machine learning algorithms have been developed previously, and for which our main paper provides novel advanced approaches on the basis of the Moran quadrant map (MQM).

#### 1.1 Tissue domain segmentation

Biological function is often regulated by complex molecular coordination between cells, localized in tissue domains that range from cellular neighborhoods to multicellular functional units and anatomical regions. Identifying spatial domains by spatial clustering is necessary when constructing atlases, such as the Human Biomolecular Atlas Program (HuBMAP). Tissue domain segmentation is a form of regionalization, or semantic segmentation, that partitions a univariate or multivariate image into spatial regions that are needed for subsequent analyses [1–3]. For example, segmentation is a prerequisite for the identification of molecular species with significantly different expression profiles between diseased and non-diseased regions of a tissue sample, which, in turn, may lead to the discovery of new biomarkers. Segmentation does not necessarily need to be spatially-informed: one traditional approach consists of applying unsupervised clustering methods such as the $k$-means or Leiden clustering algorithms to the spectral/molecular signatures of pixels without taking those pixels' spatial coordinates into account. However, incorporating spatial information along with the molecular information is generally known to produce more coherent domains.

Since our paper uses imaging mass spectrometry (IMS), a highly-multiplexed ion-based imaging technique, and photon-based multiplexed immunofluorescence (MxIF) microscopy as modalities representative of multiplexed molecular imaging, we will focus on those modalities. The following methods were developed specifically for IMS data, and two of them are used in the main paper as benchmarks against which to assess the performance of our MQM-based Moran-Felsenszwalb segmentation. Guo *et al.* have developed a univariate approach to segmentation with spatial Dirichlet Gaussian mixture models [4]. Alexandrov *et al.* [2] and Bemis *et al.* [5] have developed multivariate segmentation methods that account for spatial context. The segmentation maps obtained on IMS data by Alexandrov's spatial $k$-means clustering algorithm and by Bemis' spatial shrunken centroids algorithm tend to be less noisy and better correlated with morphological structures. Bemis' *et al.* [6] have made the R code for the spatial shrunken centroids algorithm openly available within the Cardinal package: https://cardinalmsi.org/. The spatial $k$-means clustering algorithm of Alexandrov *et al.* uses the Fastmap algorithm to map the mass spectra into an embedded feature space that encodes both spatial and spectral similarity [2]. It then performs clustering using the $k$-means clustering algorithm: rather than use squared Euclidean distances as a measure of within-cluster point-to-centroid distances, the $k$-means clustering algorithm uses a spatially-aware distance metric according to which the distance between two mass spectra depends on the squared Euclidean distances between the mass spectra localized in their respective spatial neighborhoods. Spangenberg *et al.* [7] have made an open-source Python implementation of the spatial $k$-means clustering algorithm by Alexandrov *et al.* available within the `msiFlow` package: https://github.com/Immunodynamics-Engel-Lab/msiflow/.

The BANKSY algorithm was developed by Singhal *et al.* for the cell typing and domain segmentation of spatial -omics data, such as spatial transcriptomics data [3]. BANKSY uses a pair of spatial kernels to encode the transcriptomic signature of the microenvironment around each cell, one constructed using the weighted mean of gene expression in each cell's neighborhood and the other using an azimuthal Gabor filter [3]. Singhal *et al.* have made the R and Python implementation of BANKSY openly available: https://github.com/prabhakarlab/Banksy and https://github.com/prabhakarlab/Banksy_py. In our benchmarks, we use the Python implementation.

Many tissue domain segmentation workflows have been developed specifically for spatial transcriptomics data. There are unsupervised learning approaches, such as BayesSpace [8], STAGATE [9] and CCST [10], and self-supervised learning approaches, such as GraphST [11], ConGI [12], and CAST [13]. Many of these workflows are based on graph convolutional neural networks (*e.g.*, STAGATE, GraphST, CCST,

CAST) and/or rely on contrastive learning (*e.g.*, GraphST, ConGI). The scalability of these workflows, in terms of both memory usage and runtime, varies significantly. Given the large number of workflows available to the spatial transcriptomics community, we refer the reader to the two following benchmark studies for a comprehensive overview: [14, 15].

In the main paper, we develop a novel multivariate tissue domain segmentation workflow, called Moran-Felzenszwalb segmentation, which takes advantage of the spatial statistics of multiplexed molecular imaging data through use of the MQM. We evaluate its performance against two other spatially-informed approaches, namely the spatial $k$-means clustering algorithm of Alexandrov *et al.* and the BANKSY algorithm of Singhal *et al.*.

#### 1.2 Colocalized image clustering

Colocalization refers to the quantification of the spatial similarity between images [16, 17]. In IMS, colocalized ion image clustering is the clustering of ion images with similar spatial distributions, such as those of isotopes. It is useful for identifying colocalized molecular species that may play complementary roles in physiological or pathological processes. By grouping ion images with similar spatial distributions, biologists can infer functional relationships between molecular species, such as those co-involved in metabolic pathways, signaling networks, or structural organization within tissues. Ion image clustering has many applications in IMS, ranging from technological quality control and background detection to improved analyte identification and biological process characterization [18–20].

We focus on unsupervised approaches to colocalized ion image identification and clustering. Three deep clustering workflows have been recently developed for colocalization analysis of IMS data [18, 21, 22]. The key idea of deep clustering is to use convolutional neural networks to extract spatial features on the basis of which to perform ion image clustering.

- Given that large IMS datasets are not readily available for model training, the deep clustering workflow proposed by Zhang *et al.* is based on transfer learning: a pre-trained Xception model is used to extract high-level features, termed neural ion images, from ion images. The neural ion images are then assigned to an isotope group using a spatially-agnostic clustering algorithm, such as $k$-means clustering [21].
- The noise-robust deep clustering (NRDC) workflow proposed by Guo *et al.* [18] uses a convolutional autoencoder for image denoising, a convolutional neural network for feature extraction, pairwise pseudo-labeling with $k$-nearest-neighbors-based consistency constraints for clustering, and adaptive self-paced training for ensuring robustness. The corresponding Python implementation of NRDC has been made openly available on GitHub: https://github.com/DanGuo1223/mzClustering. However, in our comparison in the main paper, we have adapted the improved implementation by Tim Daniel Rose: https://github.com/tdrose/deep_mzClustering.
- The DeepION workflow of Guo *et al.* [22] is based on BYOL (bootstrap your own latent), which is a self-supervised deep learning student-teacher architecture [23]. DeepION has four modules: one for data augmentation, one for spatial feature encoding (*i.e.*, representation learning), one for projection and prediction, and one for dimensionality reduction. In COL mode, the DeepION data augmentation strategy is specifically designed to facilitate the identification of colocalized ion images from different molecules. The ISO mode is designed for istope detection. We adapted the following Python implementation of DeepION: https://github.com/gankLei-X/DeepION.

In the main paper, we propose a new colocalized image clustering workflow, called Moran-HOG clustering. Although our Moran-HOG clustering follows a similar approach as the previously mentioned workflows (*i.e.*, spatial feature extraction followed by clustering with a spatially-agnostic clustering algorithm), it is more computationally efficient. Indeed, rather than use a convolutional neural network for spatial feature extraction, our workflow applies Histogram-of-Oriented-Gradients (HOG) to the MQM of each ion image. Clustering of the Moran-HOG features is done using the $k$-means clustering algorithm. In the main paper, we compare our Moran-HOG workflow to both the NRDC and DeepION workflows.

### 2 Supplementary Methods

#### 2.1 Higher-order spatial weights matrix

In the main paper, we define the binary spatial weights matrix $\boldsymbol{C}$. It is called a $1^{\text{st}}$-order contiguity matrix because it formalizes the spatial connectivity of $1^{\text{st}}$-order neighboring pixels. In Figure S1, the $1^{\text{st}}$-order neighborhood is defined as per the Queen-contiguity criterion. In order to quantify the spatial dependence between a given pixel and the neighbors of its neighbors, we need a $2^{\text{nd}}$-order contiguity matrix. A spatial weights matrix can be considered as a connectivity matrix of a network where the spatially-localized observations are nodes and a neighborhood relationship corresponds to a link [24]. A first-order contiguity matrix shows all paths of length one that exist in the network, and longer paths encode higher-order contiguity. Higher-order spatial weights are defined recursively by applying the spatial weights matrix to a lower-order lagged variable: a $k^{\text{th}}$-order neighbor of a pixel is a $1^{\text{st}}$-order neighbor of a $(k-1)^{\text{th}}$-order neighbor of that pixel [24, 25]. There is therefore a need to correct for duplication. For example, when computing the $3^{\text{rd}}$-order spatial lag, $1^{\text{st}}$-order neighbors should not be counted 3 times. Refer to [26] for a detailed discussion of the construction of higher-order spatial lag operators without redundant paths. We define the spatial weights matrix of order $k$ as $\boldsymbol{C}^k$ [24, 27]. For example, the $2^{\text{nd}}$-order spatial weights matrix is $\boldsymbol{C}^2$. Figure S1 illustrates the difference between pure higher-order contiguity, which excludes lower-order neighbors, and cumulative higher-order contiguity, which includes lower-order neighbors.

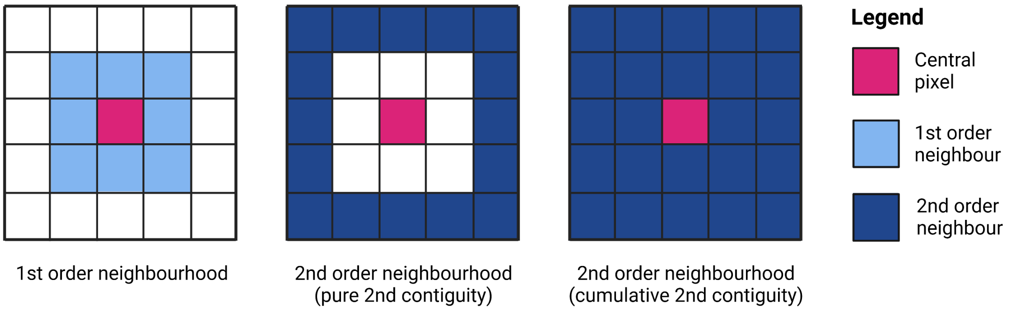

**Fig. S1**: Diagram of a $1^{\text{st}}$ and $2^{\text{nd}}$-order pixel neighborhood according to the Queen-contiguity criterion. The $1^{\text{st}}$-order neighbors share a border and/or vertex with the central pixel. In order to quantify the spatial dependence between a given pixel and the neighbors of its neighbors, we need a $2^{\text{nd}}$-order contiguity matrix. The $2^{\text{nd}}$-order neighborhood may exclude $1^{\text{st}}$-order neighbors (pure higher-order contiguity) or include them (cumulative higher-order contiguity).

#### 2.2 Moran scatterplot schematic

In the main paper, we demonstrate that Moran's $I$ is formally equivalent to the regression coefficient in a linear least squares regression of the spatial lag $\boldsymbol{Wx}$ on the mean-centered variable $\boldsymbol{x}$ [24, 28, 29]. The Moran scatterplot involves plotting each pixel's spatial lag $\mathbf{L}_{\boldsymbol{W}} x_i$ versus its mean-centered (ion) intensity $x_i$. The slope of the linear regression of the spatial lag ($y$-axis) on the mean-centered (ion) intensity ($x$-axis) is equal to Moran's $I$. In Figure S2, the Moran scatterplot is divided into four quadrants that represent four different types of local spatial autocorrelation (SAC). Since the plot is centered on the means of the (ion) intensities and spatial lag values, we can refer to an (ion) intensity value as high or low depending on whether the ($x_i$, $\mathbf{L}_{\boldsymbol{W}} x_i$)-pair is to the left or to the right of the vertical axis respectively, and we can refer to a spatial lag value as high or low depending on whether the ($x_i$, $\mathbf{L}_{\boldsymbol{W}} x_i$)-pair is above or below the horizontal axis respectively [28, 30, 31]. The upper-right and lower-left quadrants correspond to positive local SAC ($I_i > 0$): high (ion) intensity values that cluster together appear in the upper-right (high-high) quadrant, and low (ion) intensity values that cluster together appear in the lower-left (low-low) quadrant. The lower-right and upper-left quadrants correspond to negative local SAC ($I_i < 0$): spatial outlier pixels that have a higher (ion) intensity than their neighbors are in the lower-right (high-low) quadrant, whereas spatial outliers that have a lower (ion) intensity than their neighbors are in the upper-left (low-high) quadrant.

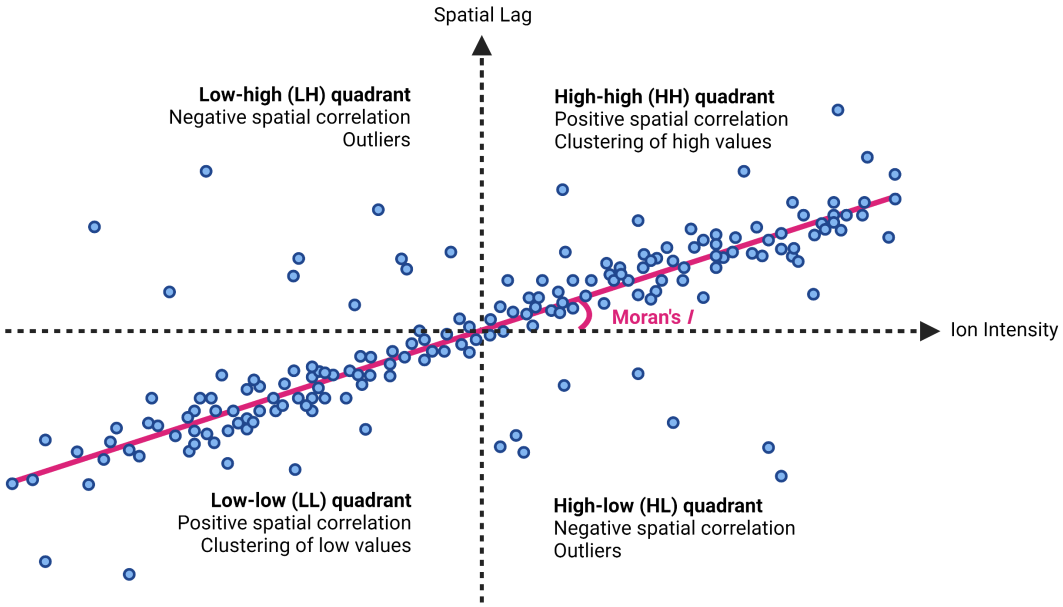

**Fig. S2**: Diagram of a Moran scatterplot obtained by doing a linear least squares regression of the spatial lag $\boldsymbol{Wx}$ onto the mean-centered ion intensity $\boldsymbol{x}$. Assuming row-standardization of the spatial weights matrix, the slope of the regression line is equal to Moran's $I$ of the image. The dotted lines correspond to the means of the spatial lag values and the ion intensities. Each pixel is classified as belonging to one of four quadrants: the high-high and low-low quadrants correspond to positive local spatial autocorrelation (SAC) (spatial clusters), whereas the high-low and low-high quadrants correspond to negative local spatial autocorrelation (SAC) (spatial outliers).

#### 2.3 Spatial filtering

Spatial filtering seeks to transform a spatially-dependent variable (*e.g.*, an ion image) into a spatially-independent variable by removing the spatial patterns embedded into it. In doing so, the original variable is partitioned into two synthetic variables: a spatial and a non-spatial component. Spatial filtering adds a new perspective to the exploratory spatial analysis of IMS data by enabling the removal of systematic, sample-wide, spatial patterns in an ion image. We follow Griffith's approach to spatial filtering, namely Moran eigenvector spatial filtering (MESF), which was first developed in [32], with its first rigorous linear regression treatment appearing in [33]. MESF is based on the eigendecomposition of the spatial weights matrix. The latter must be symmetric, which is why we use a binary contiguity spatial weights matrix $\boldsymbol{C}$ rather than the row-standardized spatial weights matrix $\boldsymbol{W}$. Given that we aim to use MESF to remove spatial dependency from ion images with thousands to hundreds of thousands of pixels, we present an implementation of MESF that is computationally efficient for large regular square tessellations.

##### 2.3.1 Spatial eigenfunction analysis

Following Griffith's approach to spatial filtering [33–35], we redefine Moran's $I$ using the $n$-by-$n$ binary spatial weights matrix $\boldsymbol{C}$ in Equation 1. This alternative definition of Moran's $I$ is written $I_C$ to differentiate it from $I$, which is defined in Equation 5 of the main paper using the row-standardized, and therefore asymmetric, spatial weights matrix $\boldsymbol{W}$. The algebraic properties of $\boldsymbol{C}$, which is a real, non-negative, symmetric, and hollow matrix, have important implications for spatial filtering. Note that different applications call for different notions of locational similarity, as a result of which Moran's $I$ can have as many definitions as there are spatial weights matrices.

$$I_C = \frac{n-q}{\sum_{i=1}^{n}\sum_{j=1}^{n} c_{i,j}} \frac{\sum_{i=1}^{n}\sum_{j=1}^{n} c_{i,j}x_i x_j}{\sum_{i=1}^{n} x_i^2} = \frac{n-q}{\sum_{i=1}^{n}\sum_{j=1}^{n} c_{i,j}} \frac{\boldsymbol{x}^T\boldsymbol{C}\boldsymbol{x}}{\boldsymbol{x}^T\boldsymbol{x}} \tag{1}$$

In section 2.3, we assume a regular square tessellation with $n$ pixels and no isolates: $q = 0$. Furthermore, we use a $n$-by-$n$ centering matrix $(\boldsymbol{I} - \boldsymbol{1}\boldsymbol{1}^T/n)$ in order to express $I_C$ in terms of the vectorized ion image

$\boldsymbol{z}$ rather than the mean-centered vectorized ion image $\boldsymbol{x}$ [34, 35]. $\boldsymbol{I}$ is the $n \times n$ identity matrix and $\mathbf{1}$ is a $n \times 1$ vector of ones. Matrix $(\boldsymbol{I} - \mathbf{1}\mathbf{1}^T/n)$ centers $\boldsymbol{z}$ such that $\left(\boldsymbol{I} - \mathbf{1}\mathbf{1}^T/n\right)\boldsymbol{z} = \boldsymbol{x}$. Given that the centering matrix is symmetric and idempotent, we have $(\boldsymbol{I} - \mathbf{1}\mathbf{1}^T/n)^T = (\boldsymbol{I} - \mathbf{1}\mathbf{1}^T/n)$ and $(\boldsymbol{I} - \mathbf{1}\mathbf{1}^T/n)^2 = (\boldsymbol{I} - \mathbf{1}\mathbf{1}^T/n)$. Equation 1 therefore simplifies to Equation 2.

$$I_C = \frac{n}{\mathbf{1}^T \boldsymbol{C} \mathbf{1}} \frac{\boldsymbol{z}^T \left(\boldsymbol{I} - \frac{\mathbf{1}\mathbf{1}^T}{n}\right) \boldsymbol{C} \left(\boldsymbol{I} - \frac{\mathbf{1}\mathbf{1}^T}{n}\right) \boldsymbol{z}}{\boldsymbol{z}^T \left(\boldsymbol{I} - \frac{\mathbf{1}\mathbf{1}^T}{n}\right) \boldsymbol{z}} \tag{2}$$

We then perform the eigendecomposition of the modified spatial weights matrix in the numerator of Equation 2: $(\boldsymbol{I} - \mathbf{1}\mathbf{1}^T/n)\boldsymbol{C}(\boldsymbol{I} - \mathbf{1}\mathbf{1}^T/n)$ is obtained by pre- and post-multiplication of the binary spatial weights matrix by the centering matrix. The eigenvalues $\lambda_j$ are the $n$ roots of the characteristic polynomial of $(\boldsymbol{I} - \mathbf{1}\mathbf{1}^T/n)\boldsymbol{C}(\boldsymbol{I} - \mathbf{1}\mathbf{1}^T/n)$, and are therefore obtained by solving Equation 3. The eigenvalues are arranged in descending order: $\lambda_1$ denotes the largest eigenvalue, which is called the principal eigenvalue, and $\lambda_n$ denotes the smallest eigenvalue. Associated with each eigenvalue $\lambda_j$ is an eigenvector $\boldsymbol{e}_j$ that satisfies Equation 4. Given that the binary spatial weights matrix $\boldsymbol{C}$ is real and symmetric, $(\boldsymbol{I} - \mathbf{1}\mathbf{1}^T/n)\boldsymbol{C}(\boldsymbol{I} - \mathbf{1}\mathbf{1}^T/n)$ is also real and symmetric. Therefore, the eigenvalues $\lambda_j$ are real numbers and the eigenvectors $\boldsymbol{e}_j$ are real and mutually orthogonal: $\boldsymbol{e}_j^T \boldsymbol{e}_k = 0$ for $j \neq k$ [34, 36]. In order to facilitate the upcoming computations, the eigenvectors are normalized: each eigenvector has a norm of one and $\boldsymbol{e}_j^T \boldsymbol{e}_j = 1$.

$$\det \left[ \left(\boldsymbol{I} - \frac{\mathbf{1}\mathbf{1}^T}{n}\right) \boldsymbol{C} \left(\boldsymbol{I} - \frac{\mathbf{1}\mathbf{1}^T}{n}\right) - \lambda \boldsymbol{I} \right] = 0 \tag{3}$$

$$\left[ \left(\boldsymbol{I} - \frac{\mathbf{1}\mathbf{1}^T}{n}\right) \boldsymbol{C} \left(\boldsymbol{I} - \frac{\mathbf{1}\mathbf{1}^T}{n}\right) - \lambda_j \boldsymbol{I} \right] \boldsymbol{e}_j = \mathbf{0} \;\; \text{for} \;\; j = 1, 2, 3, ..., n \tag{4}$$

Given that $(\boldsymbol{I} - \mathbf{1}\mathbf{1}^T/n)\boldsymbol{C}(\boldsymbol{I} - \mathbf{1}\mathbf{1}^T/n)$ is real, square, and symmetric, the spectral theorem states that it is diagonalizable as per Equation 5. The result of the eigendecomposition of the modified spatial weights matrix is written using the two following $n \times n$ matrices: the orthogonal matrix $\boldsymbol{E}$, whose columns correspond to the $n$ eigenvectors $\boldsymbol{e}_j$, and the diagonal matrix $\boldsymbol{\Lambda}$, whose diagonal elements are the $n$ eigenvalues $\lambda_j$ (for $j = 1, 2, ..., n$) [34, 36]. In addition to being mutually orthogonal, the eigenvectors of the modified spatial weights matrix in the numerator of $I_C$ are also mean-centered, and therefore mutually uncorrelated [34, 37]. Indeed, orthogonality guarantees pairwise uncorrelatedness if and only if the eigenvectors are mean-centered. Refer to section 3 of [38] for the corresponding proof. Having mutually uncorrelated eigenvectors is necessary for spatial filtering in section 2.3.2.

$$\left(\boldsymbol{I} - \frac{\mathbf{1}\mathbf{1}^T}{n}\right) \boldsymbol{C} \left(\boldsymbol{I} - \frac{\mathbf{1}\mathbf{1}^T}{n}\right) = \boldsymbol{E}\boldsymbol{\Lambda}\boldsymbol{E}^T = \sum_{j=1}^{n} \lambda_j \boldsymbol{e}_j \boldsymbol{e}_j^T \tag{5}$$

Each eigenvector of the modified spatial weights matrix $(\boldsymbol{I} - \mathbf{1}\mathbf{1}^T/n)\boldsymbol{C}(\boldsymbol{I} - \mathbf{1}\mathbf{1}^T/n)$ has a corresponding spatial map pattern. Each eigenvector is a $n \times 1$ vector whose elements have a one-to-one correspondence with a pixel in the image under study. Each eigenvector $\boldsymbol{e}_j$ can therefore be visualized as an image whose Moran's $I$ value is determined by eigenvalue $\lambda_j$ [32, 34, 35, 37]. Equation 6 relates the $I_C$ statistic of the spatial pattern determined by eigenvector $\boldsymbol{e}_j$ to its corresponding eigenvalue $\lambda_j$. Equation 6 follows from the definition of Moran's $I_C$ in Equation 2, and from the eigendecomposition of the modified spatial weights matrix in the numerator of Moran's $I_C$ in Equation 5. Refer to section 2 of [39] for the proof of Equation 6. The first eigenvector, $\boldsymbol{e}_1$ is the set of numerical values that has the largest Moran's $I$ achievable by any set for the spatial arrangement defined by the spatial weights matrix $\boldsymbol{C}$. The second eigenvector $\boldsymbol{e}_2$ is the set of values that has the largest achievable Moran's $I_C$ by any set that is uncorrelated with $\boldsymbol{e}_1$. The third eigenvector is the set of values that has the largest achievable Moran's $I_C$ by any set that is uncorrelated with $\boldsymbol{e}_1$ and $\boldsymbol{e}_2$, and so on. This sequence of eigenvectors continues through $\boldsymbol{e}_n$, which is the set of values that has the smallest Moran's $I_C$ achievable by any set that is uncorrelated with the preceding $n - 1$ eigenvectors. In conclusion, the $n$ eigenvectors of $(\boldsymbol{I} - \mathbf{1}\mathbf{1}^T/n)\boldsymbol{C}(\boldsymbol{I} - \mathbf{1}\mathbf{1}^T/n)$ represents a kaleidoscope of all the spatial patterns possible for a given geometric surface partitioning [33].

$$I_C(\boldsymbol{e}_j) = \frac{n}{\mathbf{1}^T\boldsymbol{C}\mathbf{1}} \frac{\boldsymbol{e}_j^T\left(\boldsymbol{I}-\frac{\mathbf{1}\mathbf{1}^T}{n}\right)\boldsymbol{C}\left(\boldsymbol{I}-\frac{\mathbf{1}\mathbf{1}^T}{n}\right)\boldsymbol{e}_j}{\boldsymbol{e}_j^T\left(\boldsymbol{I}-\frac{\mathbf{1}\mathbf{1}^T}{n}\right)\boldsymbol{e}_j} = \frac{n}{\mathbf{1}^T\boldsymbol{C}\mathbf{1}} \frac{\boldsymbol{e}_j^T\left(\lambda_j\boldsymbol{e}_j\boldsymbol{e}_j^T\right)\boldsymbol{e}_j}{\boldsymbol{e}_j^T\boldsymbol{e}_j} = \lambda_j \frac{n}{\mathbf{1}^T\boldsymbol{C}\mathbf{1}} \tag{6}$$

Equation 6 demonstrates why the extreme eigenvalues of the modified spatial weights matrix, $\lambda_1$ and $\lambda_n$, respectively define the maximum and minimum values of the Moran's $I_C$ global SAC statistic: $(\lambda_1, \boldsymbol{e}_1)$ and $(\lambda_n, \boldsymbol{e}_n)$ optimize its Rayleigh quotient [32, 34, 40]. Equation 7 follows from Equation 6. Refer to [41] for the corresponding proof.

$$I_{C,\max} = \frac{n}{\mathbf{1}^T\boldsymbol{C}\mathbf{1}}\lambda_1 \qquad I_{C,\min} = \frac{n}{\mathbf{1}^T\boldsymbol{C}\mathbf{1}}\lambda_n \tag{7}$$

We can express the range of Moran's $I_C$ as a function of the eigenvalues of the binary spatial weights matrix $\boldsymbol{C}$, rather than the eigenvalues of the modified spatial weights matrix $(\boldsymbol{I}-\mathbf{1}\mathbf{1}^T/n)\boldsymbol{C}(\boldsymbol{I}-\mathbf{1}\mathbf{1}^T/n)$. Note that, unlike the extreme eigenvalues of the row-standardized spatial weights matrix $\boldsymbol{W}$ (refer to the main paper's Methods section 4.1), those of $\boldsymbol{C}$ do not fall within a set range and must therefore be computed for each new instance of $\boldsymbol{C}$ [41, 42]. Assuming a large $n$, the pre- and post-multiplication of $\boldsymbol{C}$ by $(\boldsymbol{I}-\mathbf{1}\mathbf{1}^T/n)$ replaces the principal eigenvector of $\boldsymbol{C}$ with a vector that is proportional to vector $\mathbf{1}$ (specifically, $1/\sqrt{n}\,\mathbf{1}$ to verify normalization) and whose corresponding eigenvalue is zero [32, 38, 42]. If the non-principal eigenvectors of $\boldsymbol{C}$ are centered and renormalized, then as $n$ increases they converge upon the eigenvectors of $(\boldsymbol{I}-\mathbf{1}\mathbf{1}^T/n)\boldsymbol{C}(\boldsymbol{I}-\mathbf{1}\mathbf{1}^T/n)$. Accordingly, all the eigenvalues of $\boldsymbol{C}$, except for its principal eigenvalue, approximately equal and asymptotically converge upon one of the eigenvalues of $(\boldsymbol{I}-\mathbf{1}\mathbf{1}^T/n)\boldsymbol{C}(\boldsymbol{I}-\mathbf{1}\mathbf{1}^T/n)$ [32, 38, 42]. As a result, for a large $n$, the extreme eigenvalues of the modified spatial weights matrix (*i.e.*, $\lambda_1$ and $\lambda_n$) are approximately equal to the smallest and second-largest eigenvalues of the binary spatial weights matrix [40]. Refer to section 3 of [38] for the corresponding proofs.

##### 2.3.2 Moran eigenvector spatial filtering

MESF uses the eigendecomposition of the modified spatial weights matrix $(\boldsymbol{I}-\mathbf{1}\mathbf{1}^T/n)\boldsymbol{C}(\boldsymbol{I}-\mathbf{1}\mathbf{1}^T/n)$ to model, and potentially remove, latent spatial patterns from a spatially-localized variable, for example a vectorized ion image. Griffith frames MESF as an ordinary least squares regression problem whose aim is to model the spatial structure of the spatially-localized variable by a linear combination of spatial eigenvectors [33, 37, 43]. In Equation 8, MESF decomposes $\boldsymbol{z}$ into three variables: a trend ($\gamma_0$), a spatially structured variable called a spatial filter ($\boldsymbol{\gamma}_k$), and a non-spatial residual ($\boldsymbol{\varepsilon}$). In section 3.2, we compare the ion intensity distribution of several ion images ($\boldsymbol{z}$) to that of their respective non-spatial residuals ($\boldsymbol{\varepsilon}$).

$$\boldsymbol{z} = \mathbf{1}\gamma_0 + \boldsymbol{E}_k\boldsymbol{\gamma}_k + \boldsymbol{\varepsilon} \tag{8}$$

When relating the set of $n$ normalized eigenvectors of the modified spatial weights matrix to a spatially-referenced variable $\boldsymbol{z}$, the eigenvector $1/\sqrt{n}\,\mathbf{1}$ relates to the mean response ($\gamma_0 \approx \mu_z$), and the remaining $n-1$ eigenvectors relate to distinct map patterns [32, 37, 39]. As we will see in section 3.2, the smaller the eigenvalue of an eigenvector, the more fragmented its spatial pattern is. The spatial patterns that are embedded in $\boldsymbol{z}$ can usually be well approximated by a limited number of spatial eigenvectors, $k$, which is far less than $n$ ($k \ll n$). Furthermore, eigendecomposition does not tend to scale well to large datasets such as ion images due to its computational complexity of $O(n^3)$. It is therefore neither necessary nor practical to include all $n$ eigenvectors into the linear regression, and that is why it is the $n$-by-$k$ matrix $\boldsymbol{E}_k$, rather than $\boldsymbol{E}$, that is used in Equation 8. How to select a suitable subset of $k$ spatial eigenvectors is not a trivial question, and the answer ultimately depends on one's application and computational resources [37, 43]. Given that ion images exhibit positive global SAC, we can exclude from $\boldsymbol{E}_k$ all spatial eigenvectors whose corresponding Moran's $I_C$ is either negative or zero. We further reduce the set of spatial eigenvectors by only including eigenvectors whose spatial patterns have a positive Moran's $I_C$ above a predefined threshold [37, 43, 44]. We set this threshold to include only spatial eigenvectors with high positive global SAC. Specifically, we include a given eigenvector $\boldsymbol{e}_j$ into matrix $\boldsymbol{E}_k$ if $I_C(\boldsymbol{e}_j) > I_C(\boldsymbol{e}_1)/2$, where $I_C(\boldsymbol{e}_1)$ is the most extreme Moran's $I_C$ value (corresponding to the principal eigenvalue and eigenvector of the modified spatial weights matrix).

Finally, note that it is the need for uncorrelated spatial eigenvectors (*i.e.*, uncorrelated variables in the ordinary least squares regression problem) that motivates the use of the binary and symmetric spatial weights matrix $\boldsymbol{C}$, rather than the row-standardized and asymmetric matrix $\boldsymbol{W}$ [1].

[1] It is not recommended to do MESF on the basis of the eigendecomposition of $(\boldsymbol{I} - \mathbf{1}\mathbf{1}^T/n)\boldsymbol{W}(\boldsymbol{I} - \mathbf{1}\mathbf{1}^T/n)$ because the row-standardized spatial weights matrix $\boldsymbol{W}$ is asymmetric. As a result, the spatial eigenvectors would not be orthogonal, and would therefore not satisfy the assumptions of the MESF linear regression (Equation 8). We can rewrite the main paper's Equation 3 as $\boldsymbol{W} = \boldsymbol{D}^{-1}\boldsymbol{C}$, where $\boldsymbol{D}$ is a $n$-by-$n$ diagonal matrix whose $(i,i)^{\text{th}}$ entry is $\sum_j c_{i,j}$. If we were to do spatial filtering using $\boldsymbol{W}$, we could define the symmetric matrix $\boldsymbol{W}^{\star} = \boldsymbol{D}^{-1/2}\boldsymbol{C}\boldsymbol{D}^{-1/2}$ such that $\boldsymbol{W}^{\star}$ has the same eigenvalues as those of $\boldsymbol{W}$ (algebraic similarity). The eigendecomposition of $\boldsymbol{W}^{\star}$ could therefore be used as a basis for MESF [37, 45, 46].

### 3 Supplementary Results

#### 3.1 Moran-HOG clustering

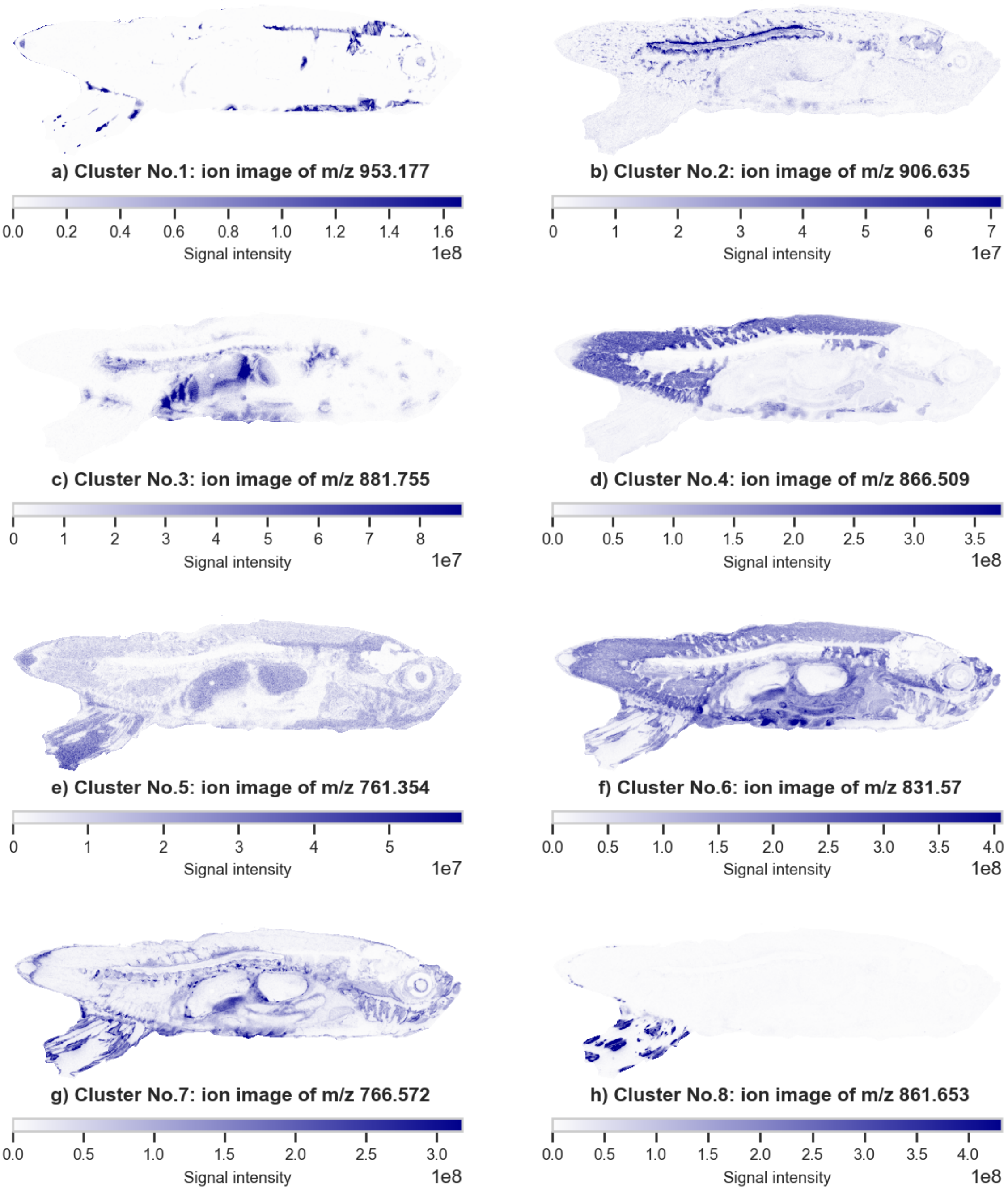

**Fig. S3**: Overview of the 8 different clusters used in the main paper's colocalization-based image clustering case study. One representative ion image from IMS dataset n$^o$2 is presented per cluster.

In section 2.3 of the main paper, we evaluate our Moran-HOG workflow against two deep-clustering workflows. We designed the clustering case study as follows: we manually selected 174 ion images from IMS dataset n$^o$2 and assigned them to one of 8 clusters. One representative ion image per cluster is presented in Figures S14, S15, S16, S17, S18, S19, S20, and S21. Each ion image is characterized by its mass-to-charge ratio ($m/z$). Some clusters are associated with a specific anatomical structure: cluster n$^o$2 corresponds to nervous tissue (*e.g.*, brain and spinal cord) and cluster n$^o$4 corresponds to muscle. The clusters have varying sizes: the smallest cluster is made up of only 6 ion images, whereas the largest cluster is made up of 45 ion images. Each ion image in these more detailed figures is presented together with its MQM and the HOG images of both the (scaled) ion image and the MQM. The MQMs are obtained by computing the local Moran statistics of each ion image using a cumulative 5$^{\text{th}}$-order Queen-contiguity spatial weights matrix.

#### 3.2 Moran eigenvector spatial filtering

Whereas the main paper focuses on methods to extract and utilize spatial-statistical information, the purpose of section 3.2 is to demonstrate how spatial patterns can be removed from IMS data using Moran eigenvector spatial filtering (MESF). IMS dataset n$^o$4 is a negative ionization mode HuBMAP dataset, obtained from a section of human kidney tissue from 43-year-old Caucasian male donor. The dataset's corresponding HuBMAP sample ID is VAN0030-LK-1-34 and its DOI is 10.35079/HBM252.SRFF.799. The data can be downloaded from the HuBMAP portal: https://portal.hubmapconsortium.org/browse/dataset/b0ec8f348a3725034359c911a3fe5037.

We start by defining a binary, and therefore symmetric, 1$^{\text{st}}$-order Queen-contiguity spatial weights matrix $\boldsymbol{C}$. We proceed to the eigendecomposition of the modified spatial weights matrix $(\boldsymbol{I} - \boldsymbol{11}^T/n)\boldsymbol{C}(\boldsymbol{I} - \boldsymbol{11}^T/n)$ as per Equation 5. Given that the ion images of IMS dataset n$^o$4 share the same spatial configuration because they were acquired from the same tissue section with the same pixel size, they all share the same spatial eigenvectors. Each ion image is a 364-by-230 rectangle of pixels, with a total of $n = 83720$ pixels. Including all $n = 83720$ spatial eigenvectors into a spatial filter would severely distort the resulting ion images. Furthermore, the high computational cost of spatial eigenvector computation (*i.e.*, time complexity of $O(n^3)$) makes it unfeasible to compute all $n = 83720$ spatial eigenvectors. We therefore need to select $k$ spatial eigenvectors out of a total of $n$. Given that the ion images exhibit statistically-significant levels of positive global SAC, we define our spatial filter using the $k = 2000$ spatial eigenvectors with the largest eigenvalues (and the highest Moran's $I_C$ statistics). The map patterns of some of these eigenvectors are presented in descending order of eigenvalue in Figure S4. In Figure S4, the principal eigenvector, written $\boldsymbol{E}_1$, has an eigenvalue of $\lambda_1 = 7.999$, which is the largest eigenvalue, and its map pattern has the maximum possible Moran's $I_C$ statistic of $I_C = 1.005$. The Moran's $I_C$ statistic of a given spatial eigenvector is equal to the product of $n/\boldsymbol{1}^T\boldsymbol{C}\boldsymbol{1} \approx 0.126$ with its eigenvalue (refer to Equation 6). The scale of the spatial patterns encoded by the spatial eigenvector maps decreases with their Moran's $I_C$ statistic. Note that, although our work demonstrates the utility of spatial eigenvectors for spatial filtering, spatial eigenvectors have many other applications, such as missing value imputation [47].

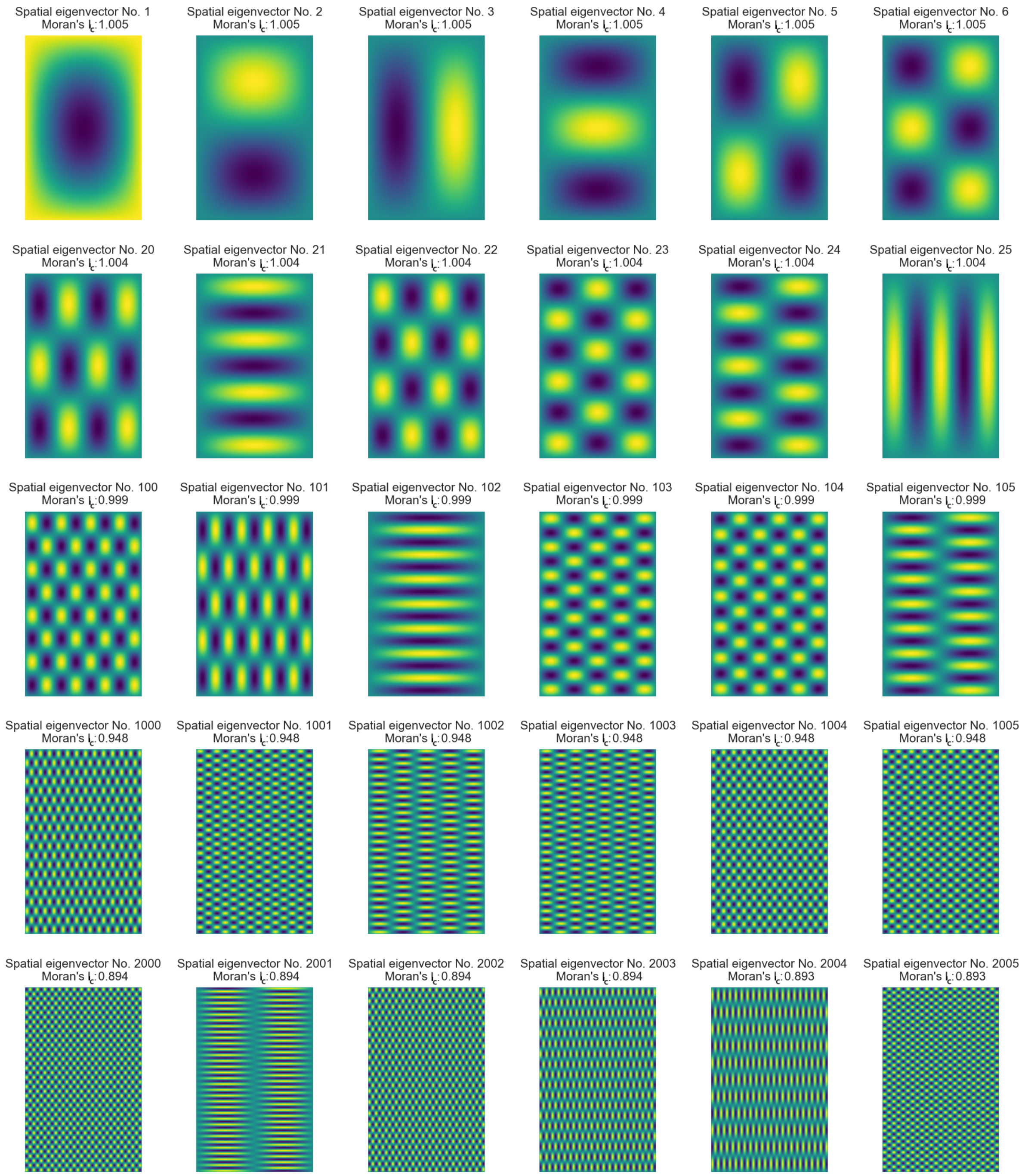

**Fig. S4**: Map patterns of the spatial eigenvectors of IMS dataset n$^o$4 obtained by eigendecomposition of its binary 1$^{\text{st}}$-order Queen-contiguity spatial weights matrix. The map patterns are arranged, from top-left to bottom-right, in descending order of global SAC as measured by Moran's $I_C$. As the degree of global positive SAC decreases, the map patterns describe increasingly high spatial frequency content.

Spatial filtering of all 212 ion images making up IMS dataset n$^o$4 is done by linear regression of the vectorized ion image $\boldsymbol{z}$ on the matrix $\boldsymbol{E}_k$ of spatial eigenvectors (refer to Equation 8). The different scales of the spatial patterns captured by the eigenvectors $\boldsymbol{E}_k$ allows for the removal of global, regional, and local spatial patterns from each ion image. The spatial filtering residual is defined as $\boldsymbol{\varepsilon} = \boldsymbol{z} - \mu_z - \boldsymbol{E}_k\boldsymbol{\gamma}_k$ (refer to Equation 8). The utility of spatial filtering is application dependent: some applications may call for the removal of global trends, in order to analyze local spatial patterns, other applications may require isolating spatial effects from non-spatial effects. As illustrated by Figures S5, S6, and S7, our approach effectively removes the spatial patterns embedded in ion images. We compare the Moran's $I_C$ statistic of the original ion image to that of the residual for all 212 ion images. We observe a median Moran's $I_C$ reduction rate of 82%.

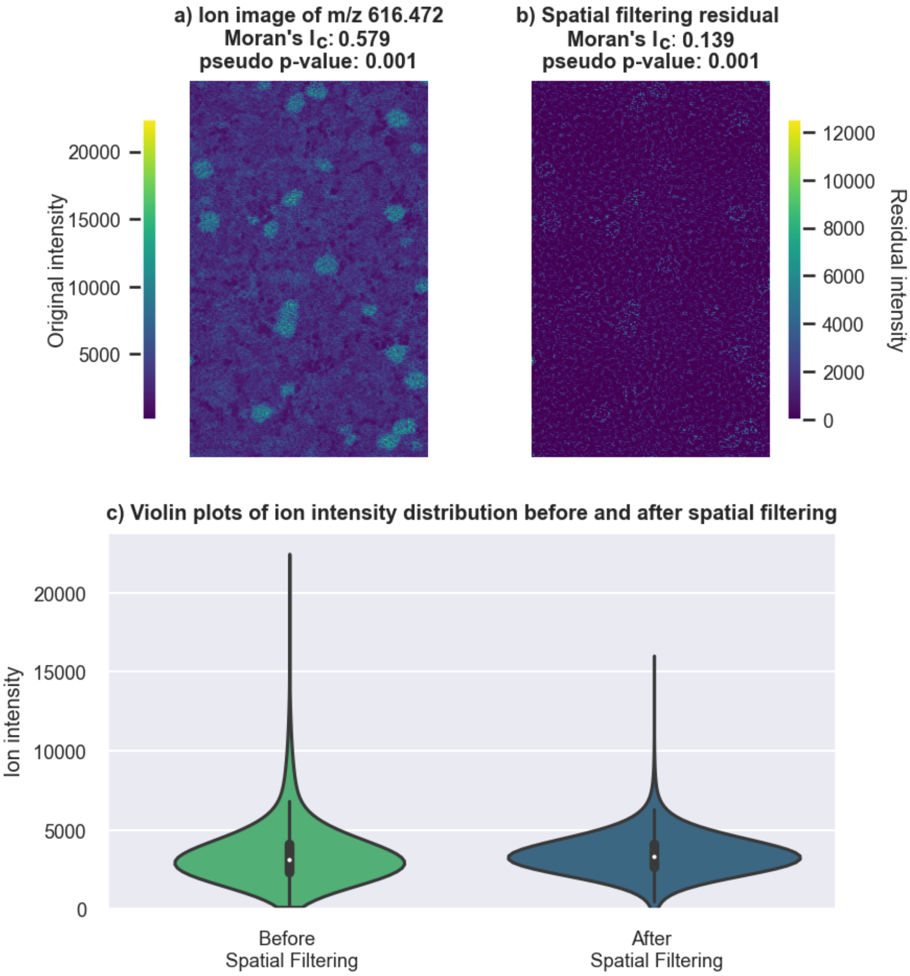

**Fig. S5**: Moran eigenvector spatial filtering of the ion image of $m/z$ 616.472 of IMS dataset n$^o$4. Moran eigenvector spatial filtering (with 2000 spatial eigenvectors) reduces the Moran's $I_C$ from 0.579 to 0.139. It also reduces the variance by 52.12%, the skew by 73.98%, and the kurtosis by 58.40%.

We compare the variance, skew, and kurtosis of the ion intensity distribution before and after MESF. MESF reduces the variance of the ion intensity distribution for all 212 ion images. The median variance reduction rate is 27% across all ion images. Furthermore, 10% of the ion images see a variance reduction of more than 55%. These results align with our expectation that positive global SAC increases variance [40]. However, to our knowledge, we are the first to quantify the effect of positive global SAC on the distribution of molecular imaging data. When comparing the skew and kurtosis of the ion intensity distribution before and after MESF, we observe that most of the ion images see a reduction in skew and kurtosis. The median skew reduction rate is 20% and the median kurtosis reduction rate is 12%. We believe that, as the spatial resolution of IMS data increases, correctly accounting for spatial dependencies between the mass spectra of neighboring pixels will become more important. Indeed, IMS achieves increasingly smaller pixel sizes (*e.g.*, subcellular pixel sizes [48]), and smaller pixels tend to exhibit stronger spatial dependencies [49].

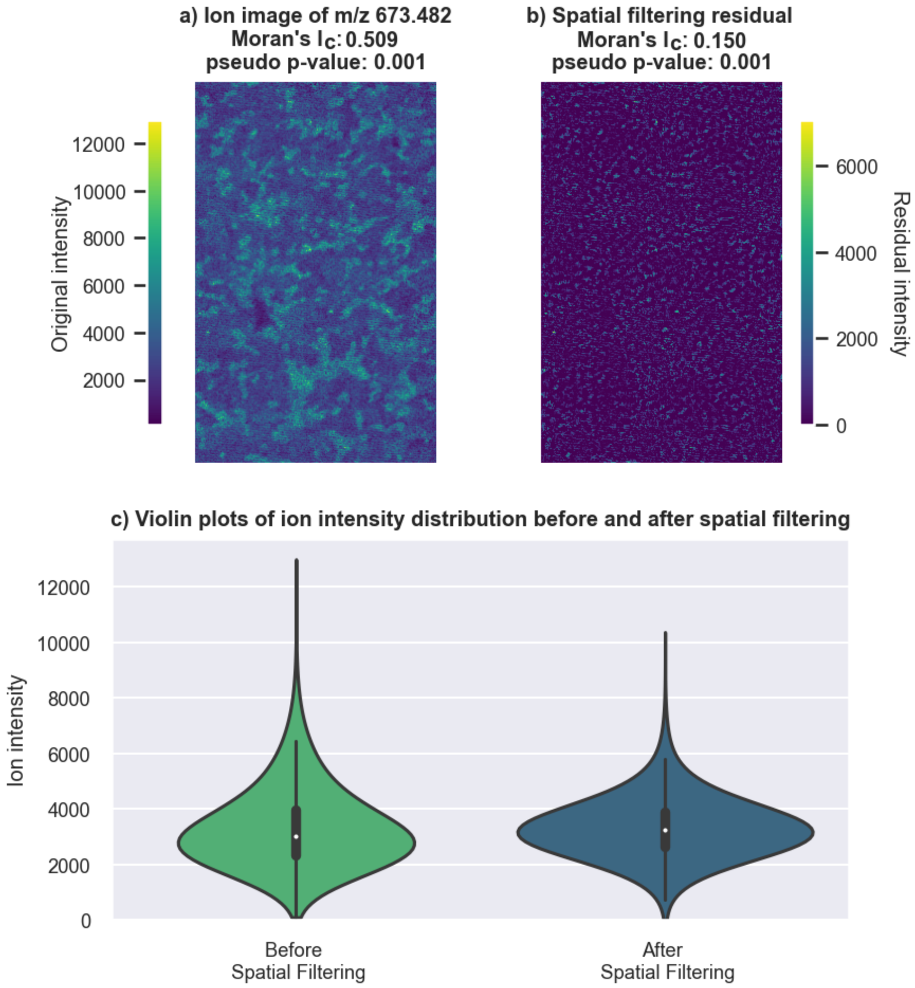

**Fig. S6**: Moran eigenvector spatial filtering of the ion image of $m/z$ 673.482 of IMS dataset n$^o$4. Moran eigenvector spatial filtering (with 2000 spatial eigenvectors) reduces the Moran's $I_C$ from 0.509 to 0.150. It also reduces the variance by 43.66%, the skew by 52.54%, and the kurtosis by 18.63%.

In Figures S5, S6 and S7, we compare the ion image, and the corresponding ion intensity distribution, of three molecular species, namely $m/z$ 616.472, $m/z$ 673.482, and $m/z$ 480.309, before and after MESF. MESF significantly reduces the global spatial dependencies of these ion images: when comparing the original ion images (Figures S5a, S6a, S7a), to the MESF residual (Figures S5b, S6b, S7b), we observe that the Moran's $I_C$ statistic of the ion image of $m/z$ 616.472 goes from 0.579 to 0.139, the Moran's $I_C$ statistic of the ion image of $m/z$ 673.482 goes from 0.509 to 0.150, and the Moran's $I_C$ statistic of the ion image of $m/z$ 480.309 goes from 0.780 to 0.347. We also observe that spatial filtering makes the distribution of these ion intensity values resemble a Gaussian distribution by reducing variance, skew, and kurtosis. The post-filtering data distributions are compatible with many statistical models and signal processing algorithms that assume that the input data (or residuals) follow a Gaussian distribution. In Figure S5c, MESF reduces the variance by 52%, the skew by 73.98%, and the kurtosis by 58.40%. In Figure S6c, MESF reduces the variance by 43.66%, the skew by 52.54%, and the kurtosis by 18.63%. Finally, in Figure S7c, MESF reduces the variance by 69.09%, the skew by 61.01%, and the kurtosis by 27.55%.

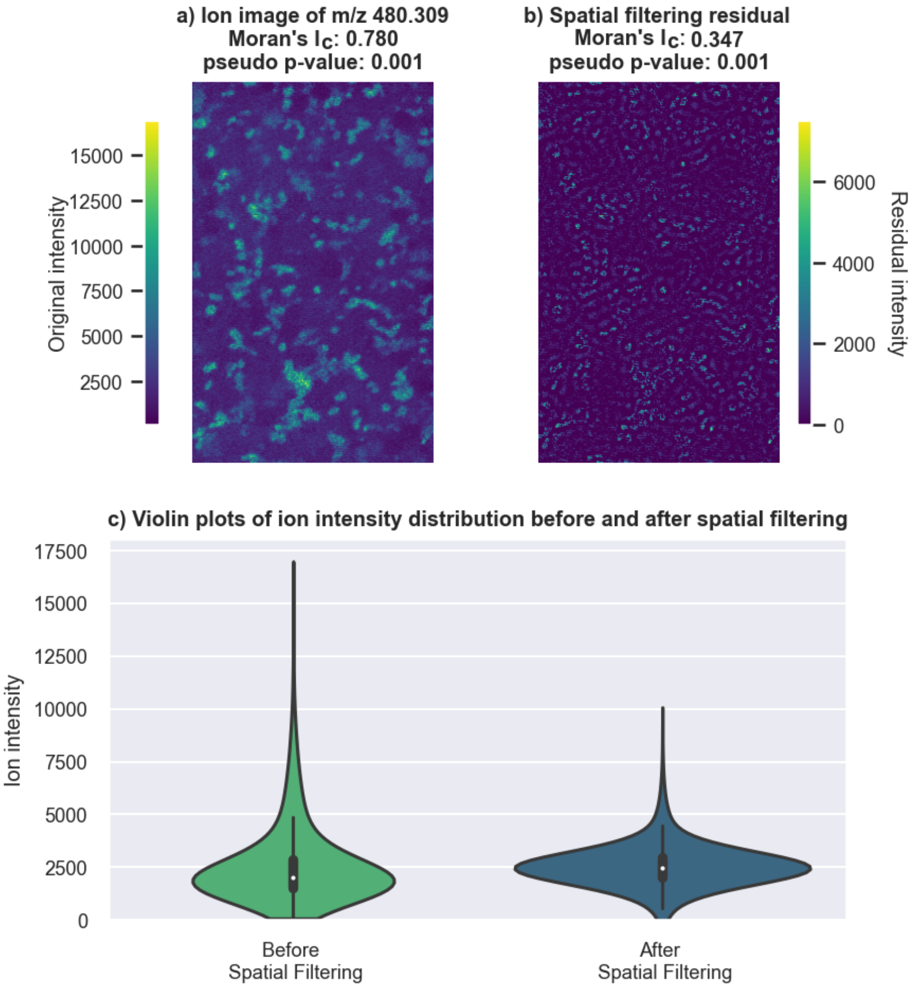

**Fig. S7**: Moran eigenvector spatial filtering of the ion image of $m/z$ 480.309 of IMS dataset n$^o$4. Moran eigenvector spatial filtering (with 2000 spatial eigenvectors) reduces the Moran's $I_C$ from 0.780 to 0.347. It also reduces the variance by 69.09%, the skew by 61.01%, and the kurtosis by 27.55%.

#### 3.3 Moran Imaging Python toolbox

All our results were generated using a new open-source Python package, called `moran_imaging`, that we developed specifically for the spatial analysis of molecular imaging data. The `moran_imaging` package is maintained on our research group's GitHub profile: https://github.com/vandeplaslab/Moran_Imaging. Our code is licensed under the BSD License, ensuring free use, modification, and redistribution with attribution. It is released via the Python Package Index (PyPI) software repository: https://pypi.org/project/Moran-Imaging/. The `moran_imaging` package is user-friendly and has minimal dependencies: it relies upon basic scientific computing and plotting Python packages (*e.g.*, `numpy`, `scipy`, `scikit-learn`, and `matplotlib`). As a result, it allows for easy installation and easy interfacing with other Python packages. It provides the statistical tools to identify, visualize, and quantify spatial patterns in IMS data, as well as other types of molecular imaging data, such as multiplexed immunofluorescence data or imaging mass cytometry data. Our Python package is computationally efficient thanks to `numba` and `joblib`. Its scalability addresses the need for spatio-molecular analysis of increasingly large molecular imaging datasets. Note that, unlike spatial statistics methods implemented by spatial data analysis libraries such as `libpysal` and `esda` [50], `moran_imaging` supports missing data, and can therefore be applied to molecular imaging data with off-tissue pixels. For example, IMS datasets n$^o$1, n$^o$2, and n$^o$3 have off-tissue pixels because they were acquired from non-rectangular tissue samples, whereas the MxIF dataset and IMS dataset n$^o$4 do not have any off-tissue pixels. We hope that our Python package and the particular adaptations it provides will facilitate the adoption of spatial statistics within the molecular imaging community.

To facilitate reproducibility and code reuse, we share both code and data. Three datasets can be downloaded from Zenodo: https://zenodo.org/records/17399931. The first dataset is a subset of 100 ion images of IMS dataset n$^o$1, the second dataset is the UMAP embedding of IMS dataset n$^o$2 that was used for Moran-Felsenszwalb segmentation, and the third dataset is a subset of 174 ion images of IMS dataset n$^o$2 that was used for Moran-HOG clustering. These datasets are licensed under the Creative Commons Attribution-NonCommercial-NoDerivatives 4.0 International (CC BY-NC-ND) license, which permits sharing with proper attribution, but prohibits commercial use and modifications. The tutorial Jupyter notebooks available on GitHub demonstrate how to reproduce the following results:

- Figures related to the spatially-informed exploratory data analysis of IMS dataset n$^o$1: Figures 1 and 2 of the main paper and Figures S9 and S10 of the Supplementary Information.
- Figures related to Moran-Felsenszwalb segmentation of IMS dataset n$^o$2: Figure 4 of the main paper and Figure S11 and Figure S12 of the Supplementary Information.
- Figures related to the Moran-HOG clustering of IMS dataset n$^o$2: Figures S14, S15, S16, S17, S18, S19, S20, S21 of the Supplementary Information.

The following hardware specifications should be considered when interpreting the runtimes reported in sections 2.2 (Moran-Felzenszwalb segmentation) and 2.3 (Moran-HOG clustering) of the Results of the main paper. The computations were performed on a Dell workstation featuring an Intel Xeon Platinum 8270 CPU  2.70 GHz with 26 physical cores, 52 logical processors, and 1.5 TB of RAM. The workstation is also equipped with an NVIDIA Quadro RTX 5000 GPU featuring 16 GB of VRAM, supported by NVIDIA's CUDA 12.5 platform, which is optimized for parallel computations.

### 4 Supplementary Experimental Protocols

We studied five multiplexed molecular imaging datasets: four IMS datasets, which we refer to as IMS datasets n$^o$1, n$^o$2, n$^o$3 and n$^o$4, and one set of three MxIF datasets obtained from diseased human brain tissue. As per the case study overview of the main paper (section 4.7), IMS datasets n$^o$1, n$^o$2, n$^o$3, and n$^o$4 are matrix-assisted laser desorption/ionization (MALDI) IMS datasets from Vanderbilt University (Nashville, TN, USA). A Fourier-transform ion cyclotron resonance (FT-ICR) mass analyzer was used for IMS dataset n$^o$1, whereas IMS datasets n$^o$2, n$^o$3, and n$^o$4 were acquired using a trapped ion mobility spectrometry (TIMS) mass analyzer. Datasets n$^o$1 and n$^o$3 were obtained from a coronal and sagittal section of murine brain tissue, respectively. Dataset n$^o$2 is obtained from a whole-body section of a zebra fish, and Dataset n$^o$4 is obtained from non-diseased human kidney tissue as part of the HuBMAP initiative.

#### 4.1 IMS Dataset n$^o$1

**Materials**: The MALDI matrix 2,5-dihydroxyacetphenone (DHA, Sigma-Aldrich Chemical CO., St. Louis, MO, USA) was applied using a TM Sprayer (HTX Technologies, Carrboro, NC, USA) and rehydrated as described in [51, 52].

**Sample Preparation**: The brain tissue sections of the case study were collected from a rat Parkinson's disease model. Parkinson's disease is characterized by a degeneration of dopaminergic neurons (*i.e.*, neurons that synthesize dopamine) and the formation of Lewy bodies (*i.e.*, protein aggregates) in the substantia nigra. The substantia nigra is a region of the basal ganglia, which are a group of nuclei (*i.e.*, clusters of neurons) located beneath the cerebral cortex. Dopamine is the primary neurotransmitter (*i.e.*, chemical messenger) of the basal ganglia. The loss of dopaminergic neurons severely impairs the brain's motor function, leading to movement disorders. The dopaminergic neurons of a rat's left brain hemisphere were destroyed to make it resemble a diseased brain. The right hemisphere was used as a control. The rat model was obtained as follows: the nigrostriatal dopaminergic neurons of the left hemisphere were destroyed to make it resemble a diseased brain, whereas the right hemisphere was used as a control. An adult male Sprague Dawley rat was deeply anesthetized with isoflurane, pretreated with desmethylimipramine (12.5 $mg/kg$, ip) and placed in a stereotaxic frame. After an incision of the dorsal surface of the skull and placement of a burr hole, the animal was injected with despramine. 10 min later 1.5:L of 6-hydroxydopamine HBr (6-OHDA; 4.0:$g/\mu L$, free base) was unilaterally injected into the substantia nigra (AP:-5.4; L:2.3; DV:-8.4) to selectively destroy nigrostriatal dopaminergic neurons. Although there is a crossed-nigrostriatal pathway, it is quite small, contributing well under 5% of the dopamine content to the contralateral striatum, and thus the contralateral striatum is usually referred to as the intact (control) side. All in-house animal experiments were performed with approval by the Vanderbilt Institutional Animal Care and Use Committee. Brain tissue was harvested, snap frozen using liquid nitrogen, and stored at -80$^oC$ until use. Frozen brain tissue was sectioned in the coronal plane at 10$\mu m$ using a cryostat (-20$^oC$, Leica CM3050S; Buffalo Grove, IL, USA) and thaw mounted onto conductive Indium-tin-oxide coated glass slides (Delta Technologies, Loveland, CO, USA). Samples were washed to remove interfering lipids and salts in sequential washes of 70% ethanol (30 s), 100% ethanol (30 s), Carnoy's fluid (6:3:1 ethanol:chloroform: acetic acid) (2 min), 100% ethanol (30 s), water with 0.2% TFA (30 s), and 100% ethanol (30 s) [52].

**Imaging Mass Spectrometry**: MALDI IMS images were collected using a 15 T FT-ICR mass spectrometer (Bruker Daltonics, Billerica, MA, USA) with a spatial sampling resolution of 75 $\mu m$ (laser spot size $\approx$ 50 $\mu m$) and a mass resolving power of 50,000 ($m$/FWHM) at $m/z$ 5000. The molecular images focused on a $m/z$ range of 1300 to 23,000 with a total of $\approx$ 20,000 pixels. The instrument was tuned for protein imaging as described in [51].

**Data Processing**: Data were imported into MATLAB 2015b (The Mathworks Inc., Natick, MA) for further data analysis, where they were normalized to TIC and peak picked, resulting in a total of 2611 peaks. Refer to [53] for additional information.

#### 4.2 MxIF Dataset Cohort

**Sample Preparation**: The frontal cortex of a human brain donor with severe Alzheimer's disease and severe cerebral amyloid angiopathy was frozen in liquid nitrogen, stored in a -80$°C$ freezer, embedded with 15% fish gelatin, and then stored again in a -80$°C$ freezer. Tissue was thawed and cryosectioned at 10 $\mu m$ thickness using a CM3050 S cryostat (Leica Biosystems, Wetzlar, Germany). The sections were thaw-mounted onto indium tin oxide (ITO) coated glass slides (Delta Technologies, Loveland, CO).

**Immunofluorescence Microscopy**: After the collection of the post-IMS autofluorescence image, stain microscopy was performed. For the brain tissue, the matrix was removed with a series of ethanol (EtOH) washes (70% to 90% EtOH). The tissue was then fixed with a 4% PFA solution for 15 minutes. The PFA was removed with a series of increasing sugar solutions (10% to 30% sucrose). The tissue was then placed in a sealed petri dish with 1X TBS and photobleached for 48 hours with an LED lamp at 4$°C$ (BESTVA DC Series 1200W LED Grow Light Full Spectrum). Sections were then washed with 100 $mM$ glycine/1X TBS/0.1% Triton X-100 buffer for 30 minutes. Sections were blocked for 60 minutes at 37$°C$ in a solution containing 10% normal donkey serum in 1X TBS/0.1% bovine serum albumin. Primary antibodies for Collagen IV, $\alpha$SMA, and thiazine red were used to stain the tissue. Tissue was washed 4 times for 10 minutes each in 1X TBS/0.05% Tween-20 then coverslipped with a DAPI fluoromount. Fluorescence microscopy images of the samples were then collected with standard DAPI, eGFP, DSRed (thiazine red), cy5 (Collagen IV) and cy7 ($\alpha$SMA) fluorescent filters for the tissue samples using a Zeiss AxioScan.Z1 slide scanner (Carl Zeiss Microscopy GmbH, Oberkochen, Germany), equipped with a Colibri7 LED light source.

#### 4.3 IMS Dataset n$^o$2

**Ethics statement**: Zebrafish studies were approved by the Institutional Animal Care and Use Committees (IACUC) of Vanderbilt University Medical Center (protocol number M2200064) in accordance with the Public Health Service Policy on the Human Care and Use of Laboratory Animals under the United States of America National Institutes of Health (NIH) Office of Laboratory Animal Welfare (OLAW).

**Zebrafish husbandry and animal work**: The zebrafish line used in this study had an AB background and were maintained on a 14:10 h light:dark cycle in a recirculating aquaculture system. A single Adult (>90 days post fertilization) male zebrafish was euthanized according to IACUC protocol.

**Materials**: Acetonitrile, isopentane, and gelatin from cold water fish were purchased from Sigma-Aldrich (St. Louis, MO). The matrix 4-(dimethylamino) cinnamic acid (DMACA) with 99% purity was bought from Thermo Scientific (Waltham, MA).

**Sample preparation**: Whole-body adult male zebrafish were frozen using a dry ice/isopentane freezing method. Samples were then embedded in 10% fish gelatine followed by a second embedding of the swim bladders. Cryosectioning was done at a 10$\mu m$ thickness using a CM3050 S cryostat (Leica Biosystems, Wetzlar, Germany) and mounted onto indium tin oxide-coated glass slides (Delta Technologies, Loveland, CO, USA). Matrix application of DMACA at a concentration of $2mg/mL$ was applied using an in-house sublimation apparatus. Autofluorescence image was collected on a Zeiss Axio Scan.Z1 (Carl Zeiss Microscopy GmbH, Oberkochen, Germany) prior to MALDI imaging.

**Imaging Mass Spectrometry**: Experiment was carried out on a prototype MALDI timsTOF Pro mass spectrometer (Bruker Daltonics, Bremen, Germany) in positive ionization mode at a spatial resolution of 10$\mu m$ (30% laser power at 10 $kHz$, 200 shots per pixel).

#### 4.4 IMS Dataset n$^o$3

**Materials**: DHA and ammonium sulfate, were purchased from Sigma-Aldrich (St. Louis, MO, USA). High-performance liquid chromatography-grade ethanol and water were purchased from Fisher Scientific; control rat brain was purchased from BioIVT (Westbury, NY, USA).

**Sample Preparation**: Control rat brain was cryosectioned to $10\mu m$ thickness using a CM3050 S cryostat (Leica Biosystems) and thaw-mounted onto conductive indium tin oxide-coated glass slides (Delta Technologies). Matrix deposition protocols were based on a published procedure [54]: 2',5'-DHA with $62.5\mu M$ ammonium sulfate in 60% ethanol–water was sprayed for a final matrix density of $1.23\mu g/mm^2$. The matrix was applied using a robotic sprayer-M5 Sprayer (HTX Technologies) equipped with a sample heating tray.

**Imaging Mass Spectrometry**: MALDI TIMS IMS was performed using a prototype MALDI timsTOF Pro mass spectrometer (Bruker Daltonics) [55]. Data (71940 pixels) were collected in negative ionization mode with a spatial resolution of $20\mu m$ (14$\mu$m beam scan) over $m/z$ 1000 - 3000. The following instrument parameters were used: ESI dry gas temperature -100°C; ion transfer time -120$\mu s$; pre-pulse storage time -12$\mu s$; collision RF -3500 $Vpp$; TIMS funnel 1 RF - 450 $Vpp$; TIMS funnel 2 RF -400 $Vpp$; multipole RF -400 $Vpp$, collision cell entrance voltage $--200V$; MALDI deflection plate: -90$V$; source pressure: $\sim 2.35mBar$; TIMS ramp time -550$ms$; $1/K_0$ range 2.05 -2.17 $V \cdot s/cm^2$; Scan rate: 0.03 $V \cdot m/s$.

**Histology, Microscopy, and Tissue Annotation**: Following the IMS experiments, the matrix was removed from the sample using 100% ethanol (10$s$) and rehydrated with 70% ethanol (10$s$). Tissues were stained using a modified cresyl violet stain [56]. Brightfield microscopy was obtained using a Zeiss AxioScan Z1 slide scanner (Carl Zeiss Microscopy GmbH, Oberkochen, Germany). Annotations of anatomical regions were made on histologically stained tissue sections with the help of The Allen Mouse Brain Atlas [57] and The Rat Brain in Stereotaxic Coordinates [58].

#### 4.5 IMS Dataset n$^o$4

**Materials**: Acetone, isopentane, tetrahydrofuran (THF), acetonitrile, and methanol were purchased from Fisher Scientific (Pittsburgh, PA, USA). 1,5-Diaminonaphthalene (DAN) and carboxymethylcellulose were purchased from Sigma-Aldrich Chemical Company (St. Louis, MO, USA).

**Sample Preparation**: Human kidney tissue was surgically removed during a full nephrectomy, and remnant tissue was processed for research purposes by the Cooperative Human Tissue Network at Vanderbilt University Medical Center. Remnant biospecimens were collected in compliance with the Cooperative Human Tissue Network standard protocols and the National Cancer Institute's Best Practices for the procurement of remnant surgical research material. Participants consented to remnant tissue collection in accordance with institutional IRB policies. Half of the excised tissue was flash-frozen over an isopentane-dry ice slurry, embedded in carboxymethylcellulose, and stored at -80 $°C$ until use. Kidney tissues were cryosectioned to a 10 $\mu m$ thickness and thaw mounted onto indium tin-oxide (ITO) coated glass slides (Delta Technologies, Loveland, CO, USA) for IMS analysis or regular glass slides for histological staining. Slides were stored at -80 $°C$ and returned to $\approx$20$°C$ within a vacuum desiccator prior to further processing.

**Imaging Mass Spectrometry**: Samples for IMS analysis were coated with a 20 $mg/mL$ solution of DAN dissolved in THF using a TM Sprayer M3 (HTX Technologies, LLC, Chapel Hill, NC, USA) yielding a 1.67 $mg/cm^2$ coating (0.05 $mL/hr$, 4 passes, 40 $°C$ spray nozzle). Tissue samples were imaged immediately after matrix deposition. MALDI IMS was performed on timsTOF fleX MS system (Bruker Daltonics, Bremen, Germany). The ion images were collected in negative ion modes at 10 $\mu m$ pixel size with the beam scan set to 6 $\mu m^2$ using 150 laser shots per pixel and 18.6% laser power (30% global attenuator and 62% local laser power) at 10 $kHz$. B Data were collected from $m/z$ 150-2000 for lipid analysis.

#### 4.6 Data Preprocessing

MALDI IMS data of Dataset n$^o$2, n$^o$3, and n$^o$4 were processed using in-house developed software. Each dataset was first converted into a custom binary format optimized for storage and speed of analysis of IMS data. Each spectrum was $m/z$ aligned using between 6 and 10 alignment peaks that were automatically selected. The alignment peaks are selected by their frequency of occurrence through the dataset to ensure that several anchor points are available for each spectrum. This is in contrast to using a pre-defined set of peaks that might not be present in every spectrum, hindering the alignment process. The alignment process was accomplished using the Python msalign library (v0.2.0). Subsequently, the spectra were calibrated using four calibration points. Outlier-insensitive total ion current (TIC) normalization factors were calculated for each dataset. The normalizations were obtained by accumulating data between the 0.05 and 0.95 quantiles of each mass spectrum (5/95% TIC). Normalization aims to counteract noise factors and to project all mass spectra onto a consensus intensity scale so that intensities can be compared between spectra. We then calculate the average mass spectrum for each dataset, which was subsequently peak-picked. A separate peak list was used for each dataset to extract ion centroid data. Each ion image was obtained by integrating the area under a curve within a window of approximately $\pm 5$ ppm. This value varies slightly depending on the bin spacing in the specific mass spectrum.

### 5 Supplementary Figures

#### 5.1 IMS Dataset n$^o$1

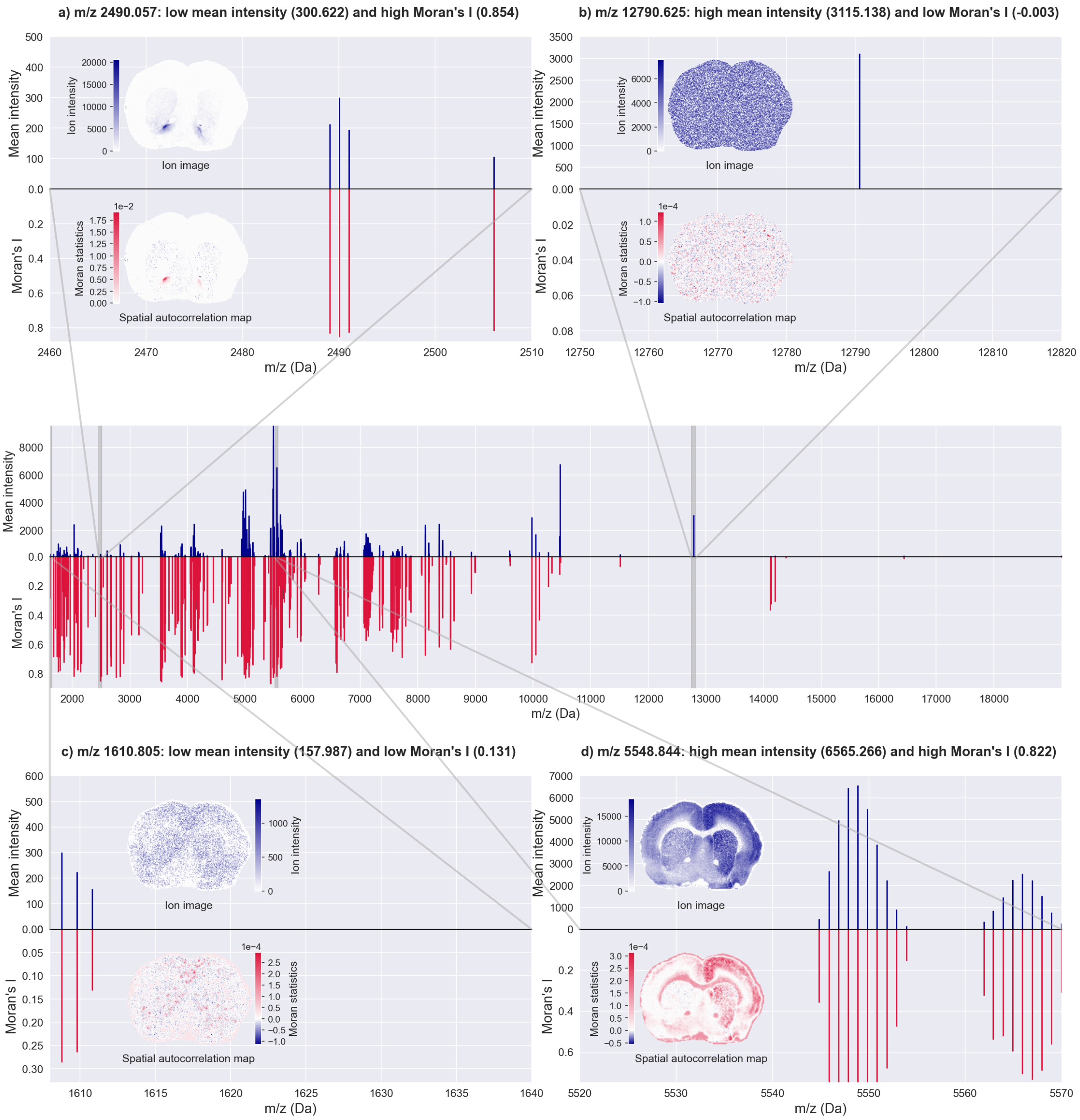

**Fig. S8**: Mean ion intensity versus global spatial autocorrelation: comparison of the mean intensity spectrum of IMS dataset n$^o$1 with a pseudo-spectrum of each $m/z$ bin's Moran's $I$ statistics. We observe that biologically informative $m/z$ bins (or peaks) with spatially-structured ion images may have low or high mean ion intensity. In contrast, noisy ion images have a low (near-zero or negative) Moran's $I$. There are four scenarios: low mean intensity and high global SAC, high mean intensity and low global SAC, low mean intensity and low global SAC, and high mean intensity and high global SAC. For each of these scenarios, the ion image is compared to its spatial autocorrelation map.

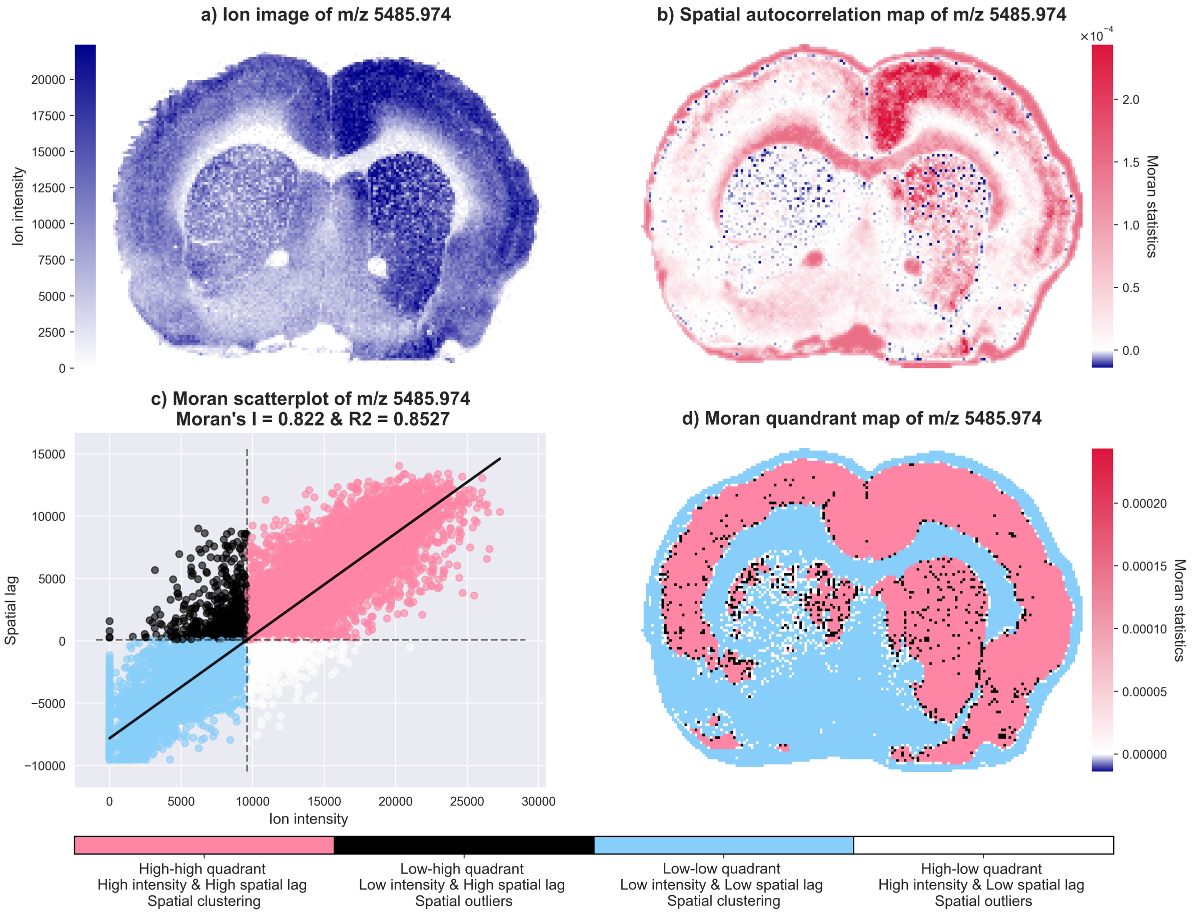

**Fig. S9**: Spatial analysis of the ion image of $m/z$ 5485.974, which has a Moran's $I$ of 0.822. The Moran scatterplot represents Moran's $I$ as the slope of a linear regression line of the spatial lag $\boldsymbol{Wx}$ on the ion intensity $\boldsymbol{x}$. It is centered on the mean ion intensity (9601.87) and the mean spatial lag (76.99). Given that the coefficient of determination is $R^2 = 0.8527$, the spatial patterns of $m/z$ 5485.974 are spatially stationary. Furthermore, 89.97% of pixels are located in the high-high and low-low quadrants of the Moran scatterplot. The asymmetry between the left and right hemispheres of the rat brain is due to the destruction of the nigrostriatal dopaminergic neurons in the left hemisphere, which is a model of Parkinson's disease (refer to section 4.1).

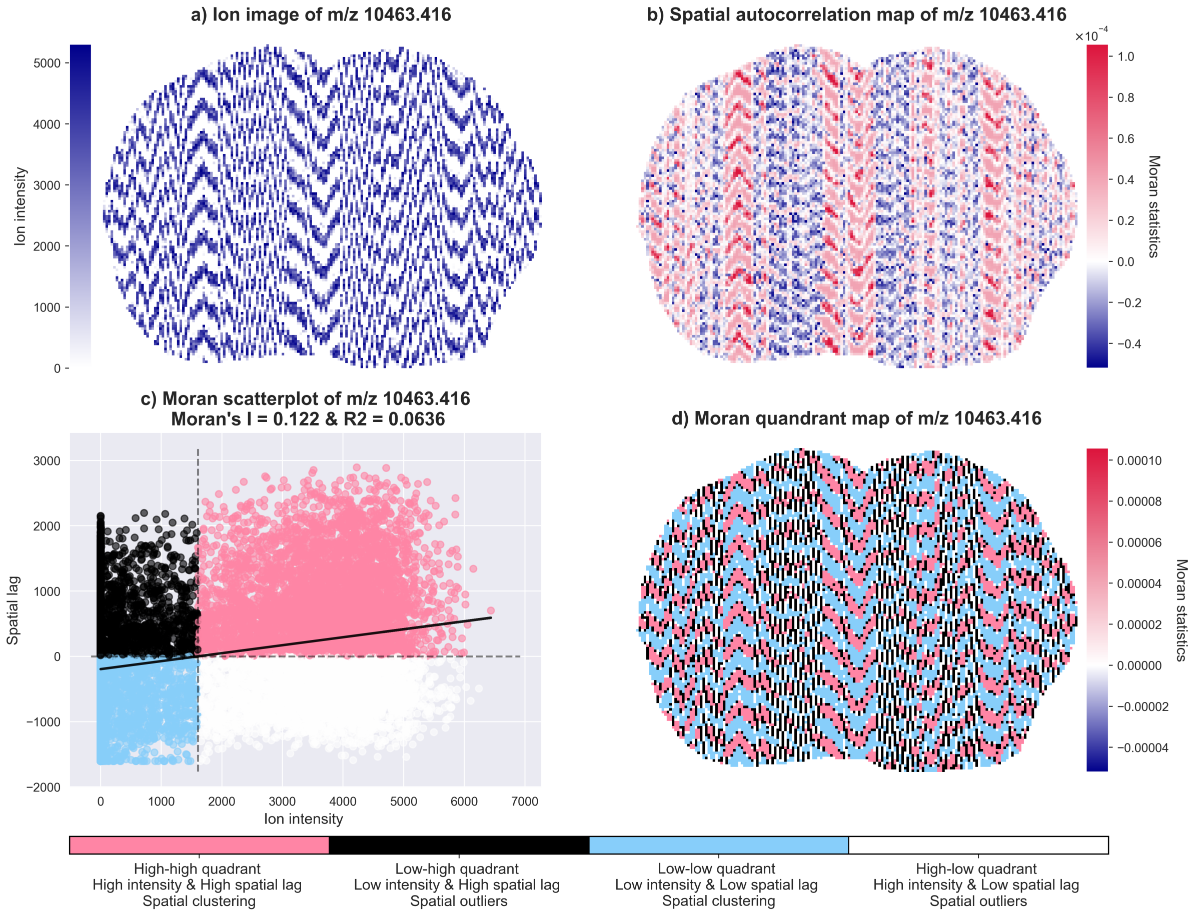

**Fig. S10**: Spatial analysis of the ion image of $m/z$ 10463.416, which has a Moran's $I$ of 0.122. The Moran scatterplot represents Moran's $I$ as the slope of a linear regression line of the spatial lag $\boldsymbol{Wx}$ on the ion intensity $\boldsymbol{x}$. It is centered on the mean ion intensity (1605.1757) and the mean spatial lag (0.1657). Given that the coefficient of determination is $R^2 = 0.0636$, the spatial patterns of $m/z$ 10463.416 are not spatially stationary. Furthermore, only 57.76% of pixels are located in the high-high and low-low quadrants of the Moran scatterplot.

#### 5.2 IMS Dataset n$^o$2

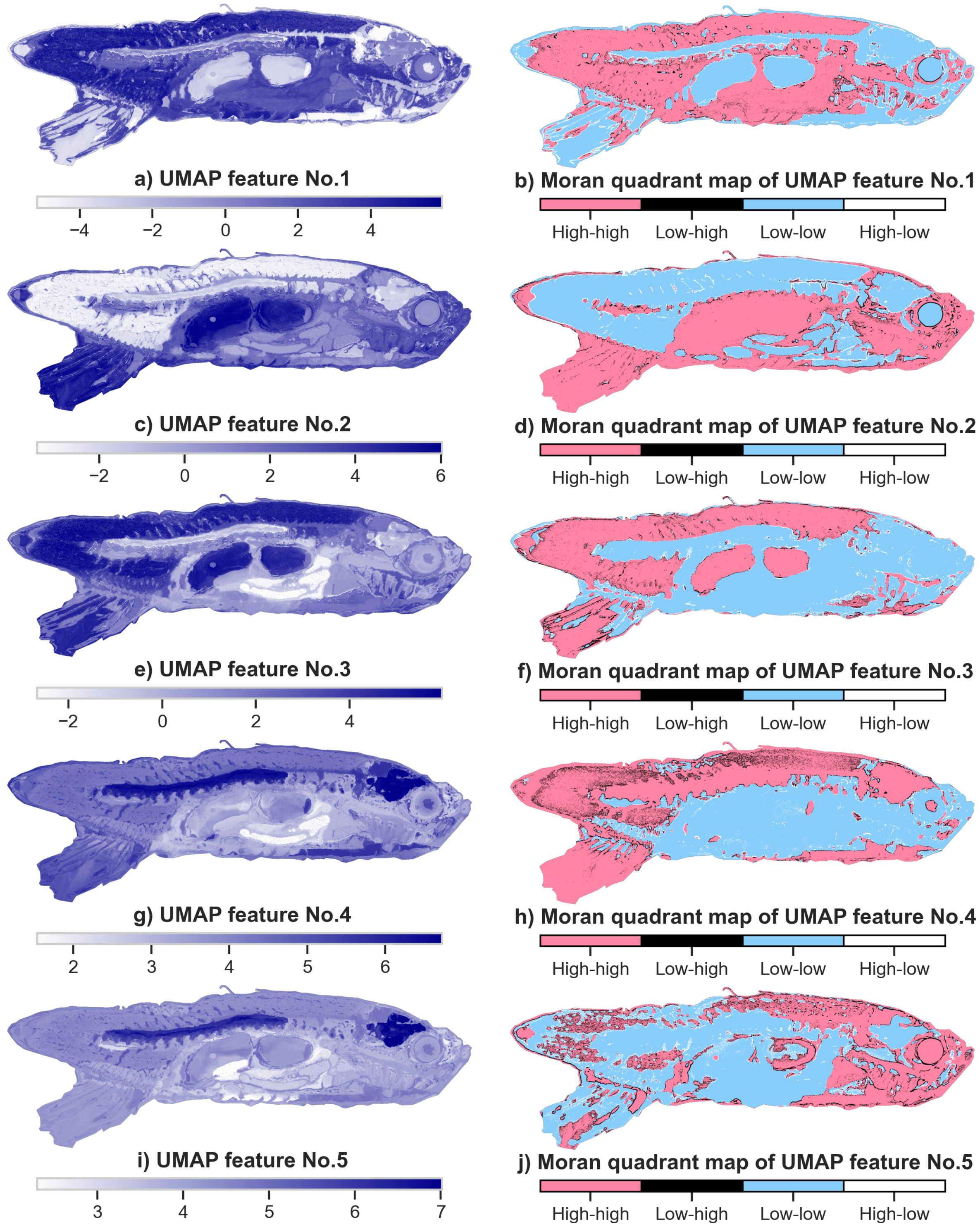

**Fig. S11**: Dimensionality reduction and processing prior to the segmentation of IMS dataset n$^o$2: top-5 UMAP features and their corresponding MQMs. The MQMs are obtained by computing the local Moran statistics of each UMAP feature map using a cumulative 5$^{\text{th}}$-order Queen-contiguity spatial weights matrix. Each Moran quadrant map facilitates the identification of different anatomical structures: **b**, spinal cord, gills, and swim bladders; **d**, brain, gastrointestinal tract, liver, and eye; **h**, vertebrae, brain, swim bladders, lens of the eye; **j**, heart, gills, and eye.

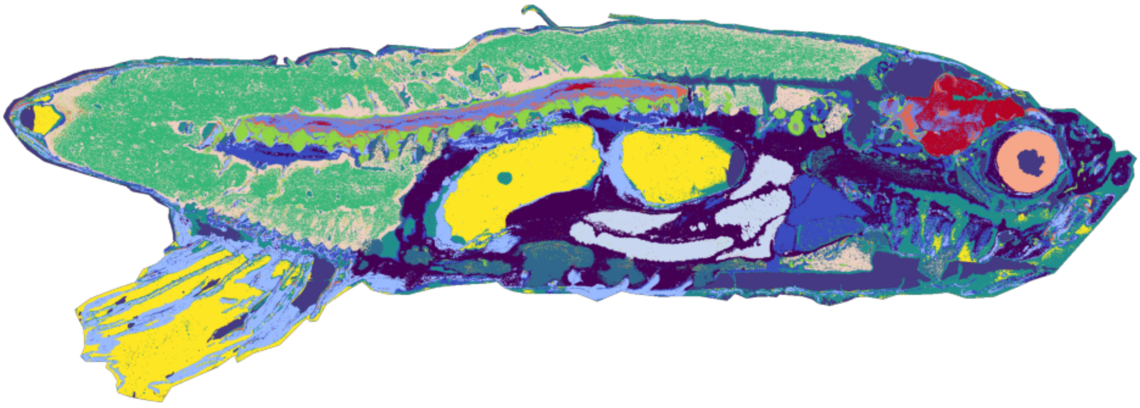

a) Result of non-spatial k-means clustering with k=13

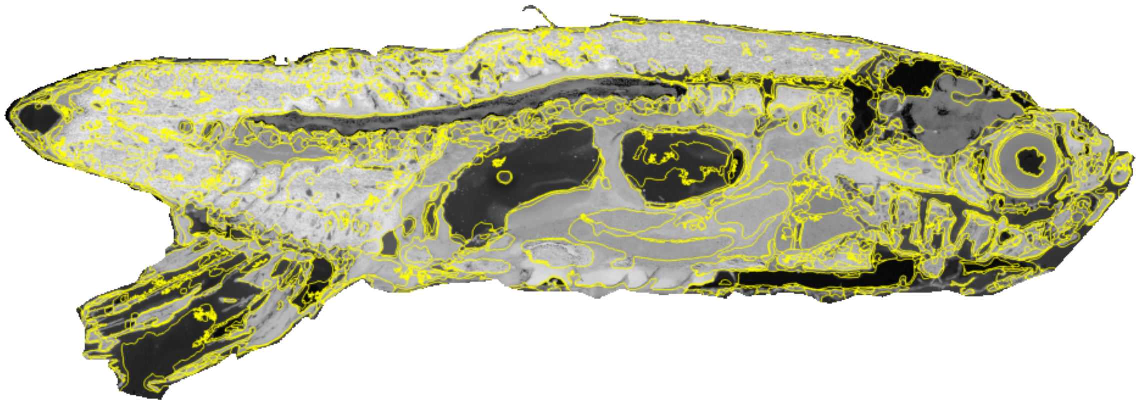

b) Felzenszwalb segmentation of Moran quadrant maps: super-pixel boundaries

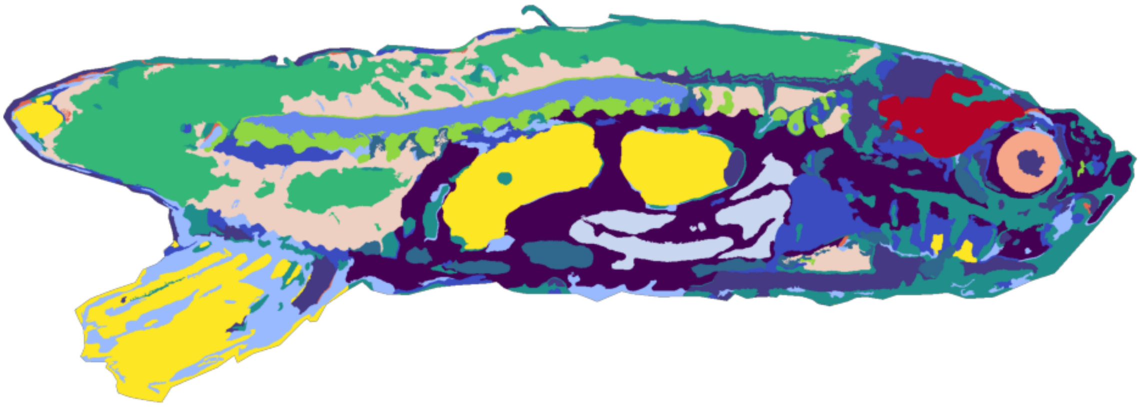

c) Felzenszwalb segmentation of Moran quadrant maps: super-pixel labeling

**Fig. S12**: Moran-Felzenszwalb segmentation of IMS dataset n$^o$2. The super-pixels are obtained by applying Felsenszwalb's segmentation algorithm to the 5 MQMs of the top-5 UMAP feature maps. The super-pixels are labeled by majority voting: the label assigned to each super-pixel is the mode of the $k$-means clustering labels of the pixels making up that super-pixel.

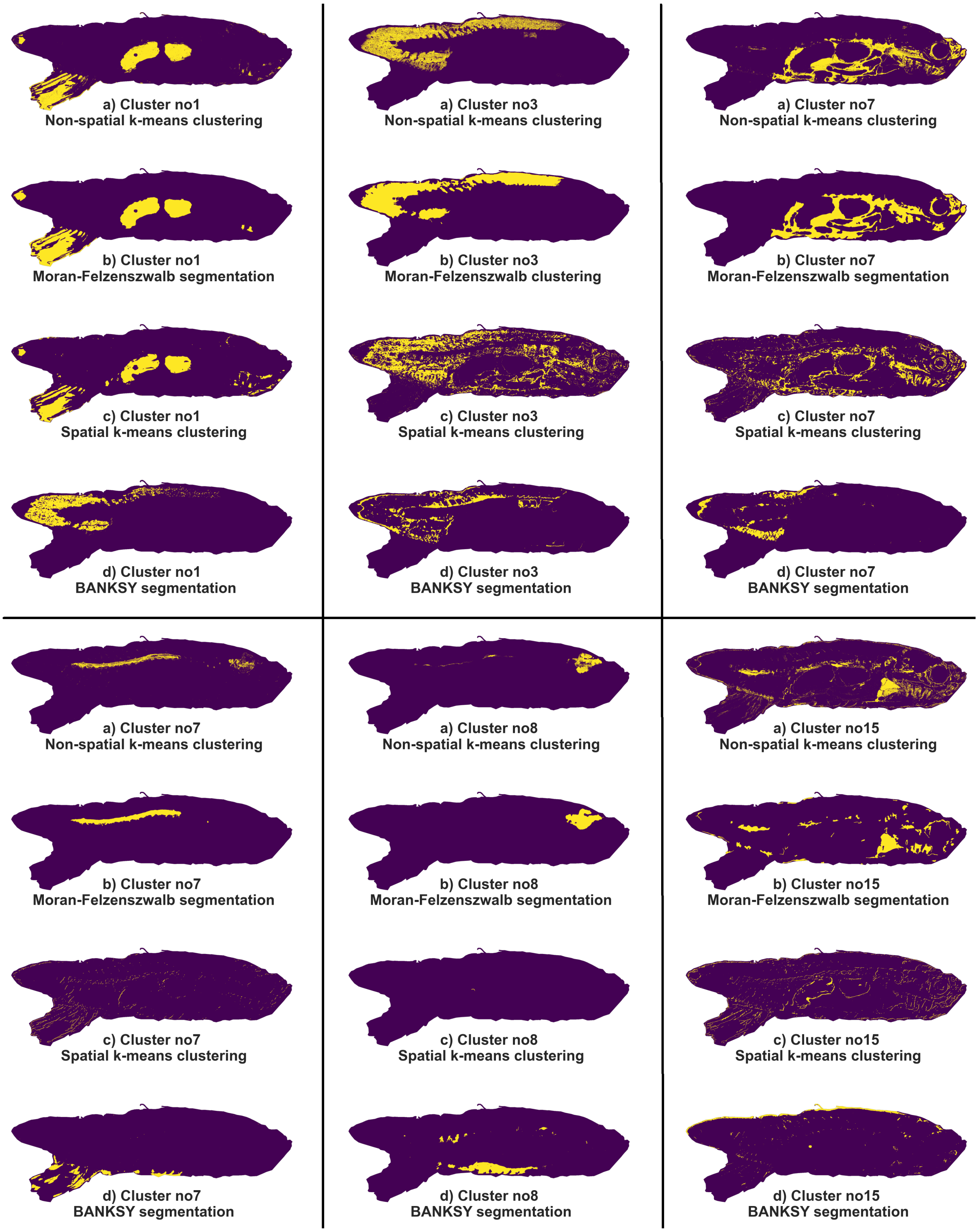

**Fig. S13**: Moran-Felzenszwalb segmentation of IMS dataset n$^o$2. The dataset is partitioned into spatially-coherent clusters, many of which correspond to anatomical regions. Cluster n$^o$1 identifies the swim bladders and caudal fin, cluster n$^o$3 identifies muscle, cluster n$^o$7 identifies the spinal cord, cluster n$^o$8 identifies the brain, and cluster n$^o$15 identifies the heart. The Moran-Felzenszwalb segmentation results are compared to the results of non-spatial $k$-means clustering, spatial $k$-means clustering, and BANKSY segmentation.

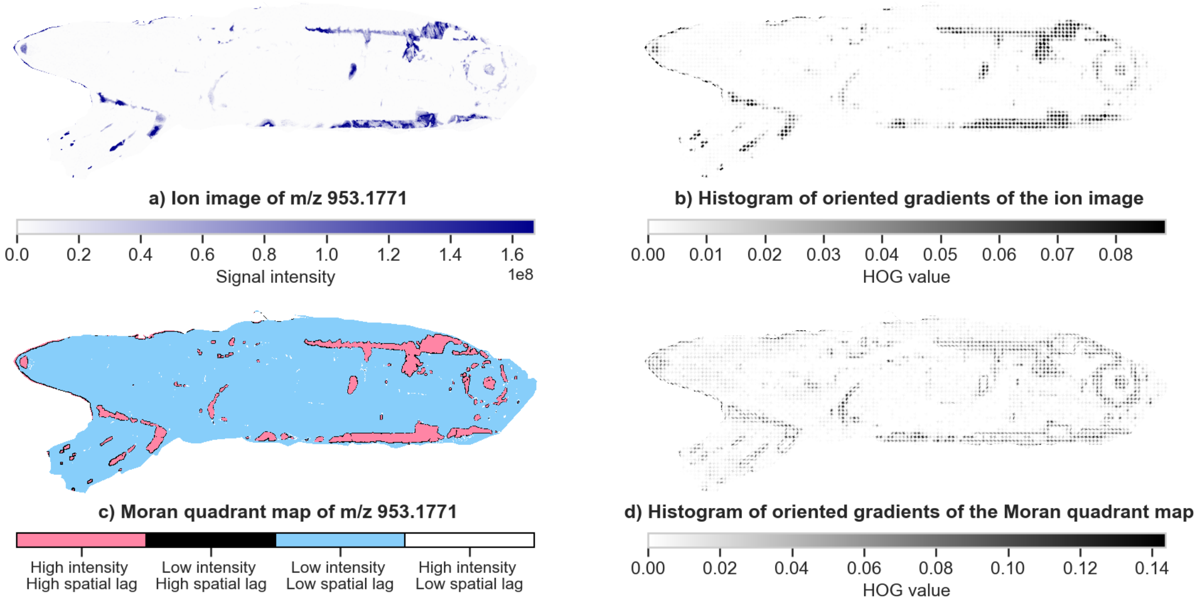

**Fig. S14**: Ion image of $m/z$ 953.1771, representative of cluster n$^o$1. The HOG features of the Moran quadrant map, rather than the HOG features of the ion image, were provided to the $k$-means clustering algorithm.

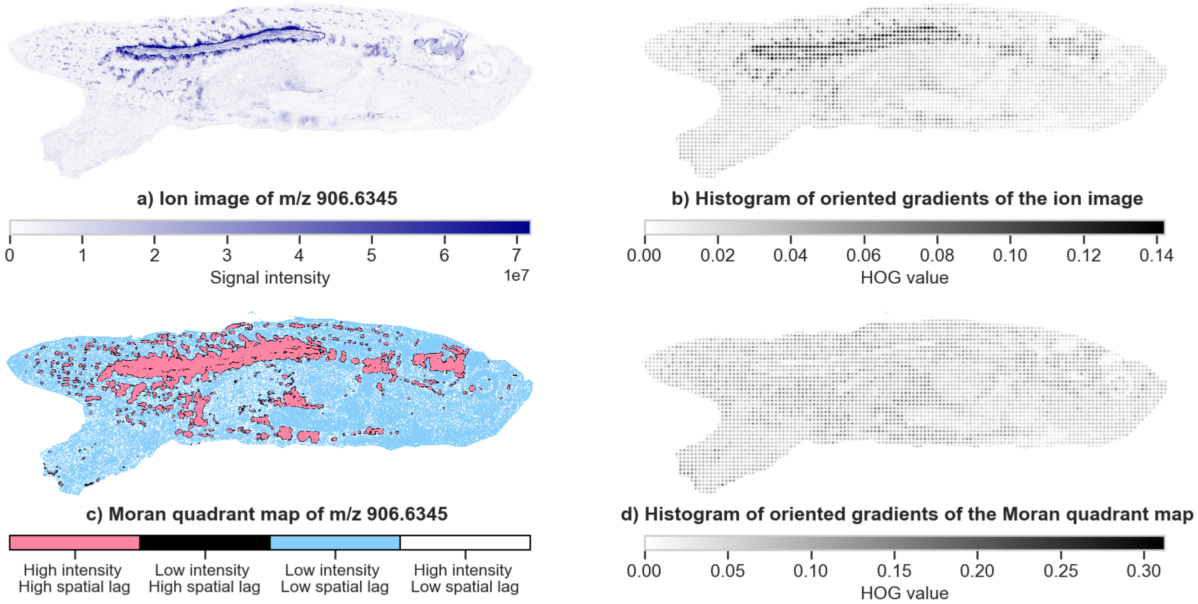

**Fig. S15**: Ion image of $m/z$ 906.6345, representative of cluster n$^o$2. The HOG features of the Moran quadrant map, rather than the HOG features of the ion image, were provided to the $k$-means clustering algorithm.

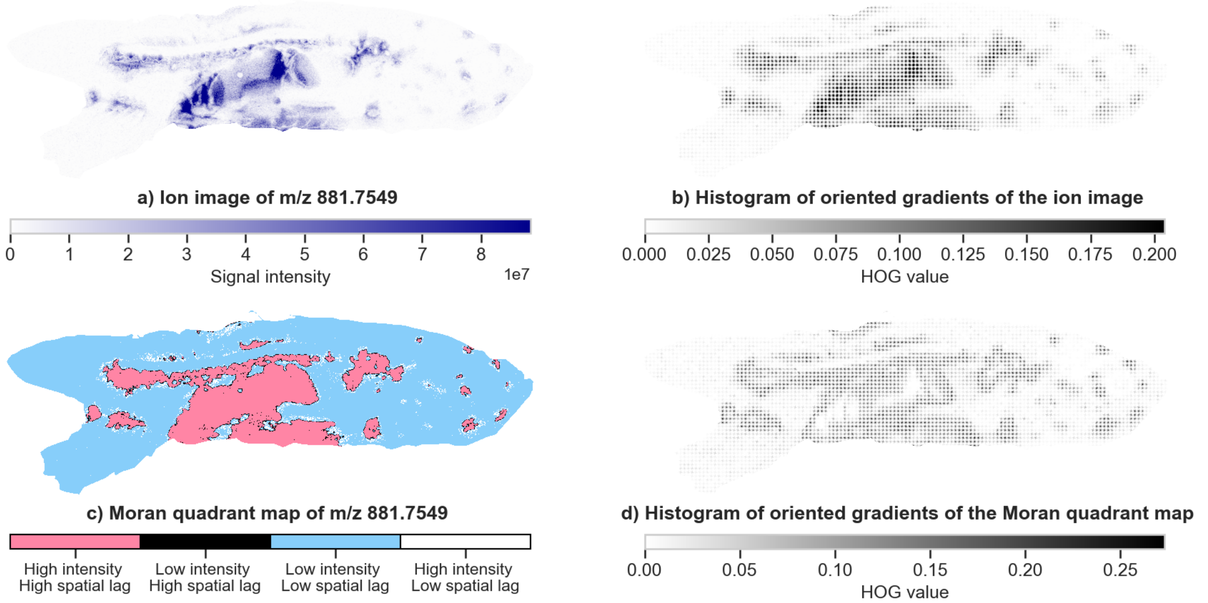

**Fig. S16**: Ion image of $m/z$ 881.7549, representative of cluster n$^o$3. The HOG features of the Moran quadrant map, rather than the HOG features of the ion image, were provided to the $k$-means clustering algorithm.

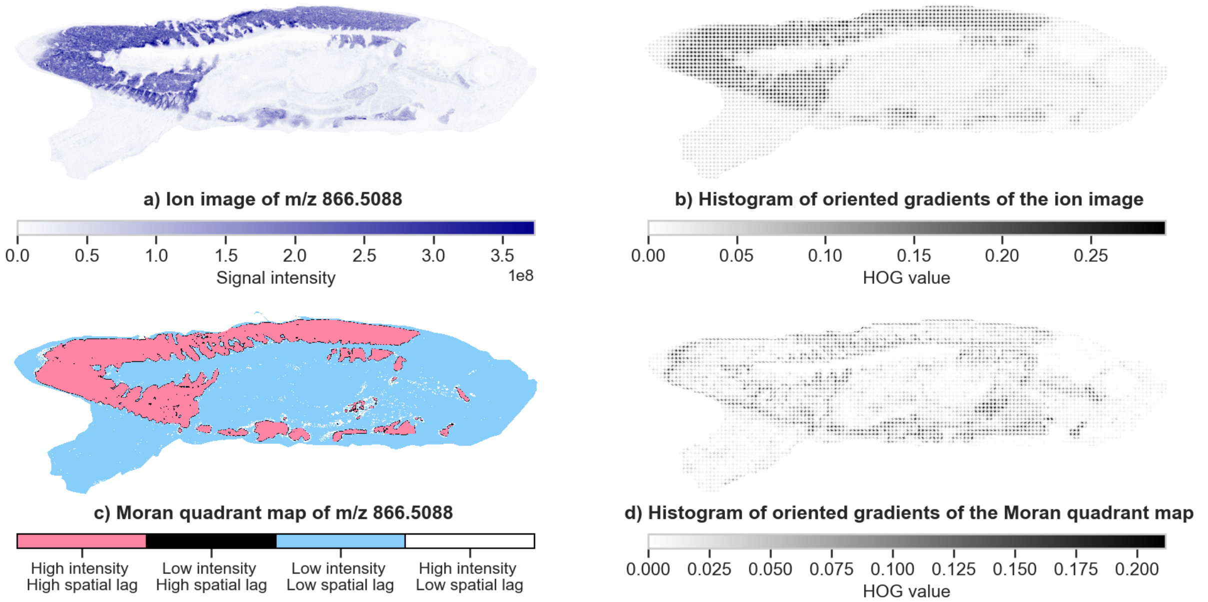

**Fig. S17**: Ion image of $m/z$ 866.5088, representative of cluster n$^o$4. The HOG features of the Moran quadrant map, rather than the HOG features of the ion image, were provided to the $k$-means clustering algorithm.

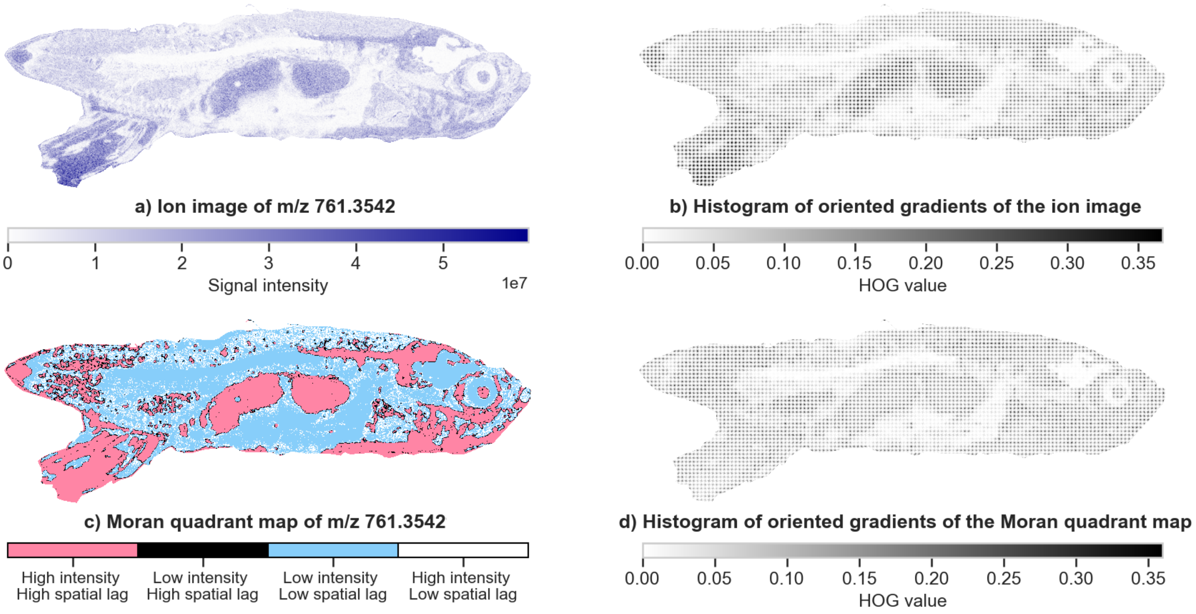

**Fig. S18**: Ion image of $m/z$ 761.3542, representative of cluster n$^o$5. The HOG features of the Moran quadrant map, rather than the HOG features of the ion image, were provided to the $k$-means clustering algorithm.

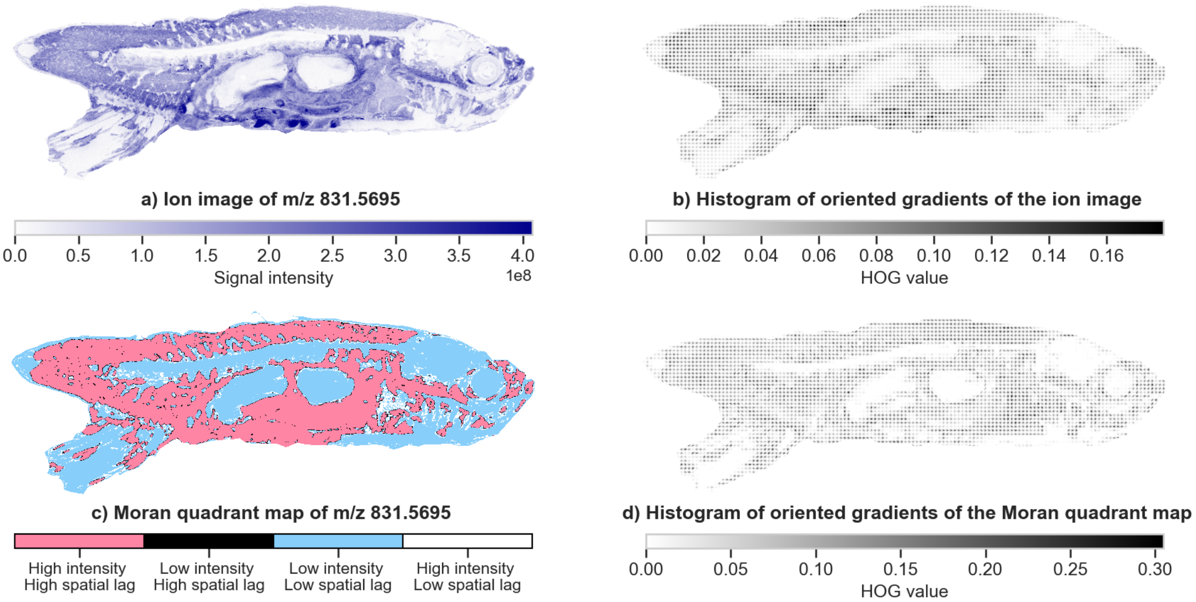

**Fig. S19**: Ion image of $m/z$ 831.5695, representative of cluster n$^o$6. The HOG features of the Moran quadrant map, rather than the HOG features of the ion image, were provided to the $k$-means clustering algorithm.

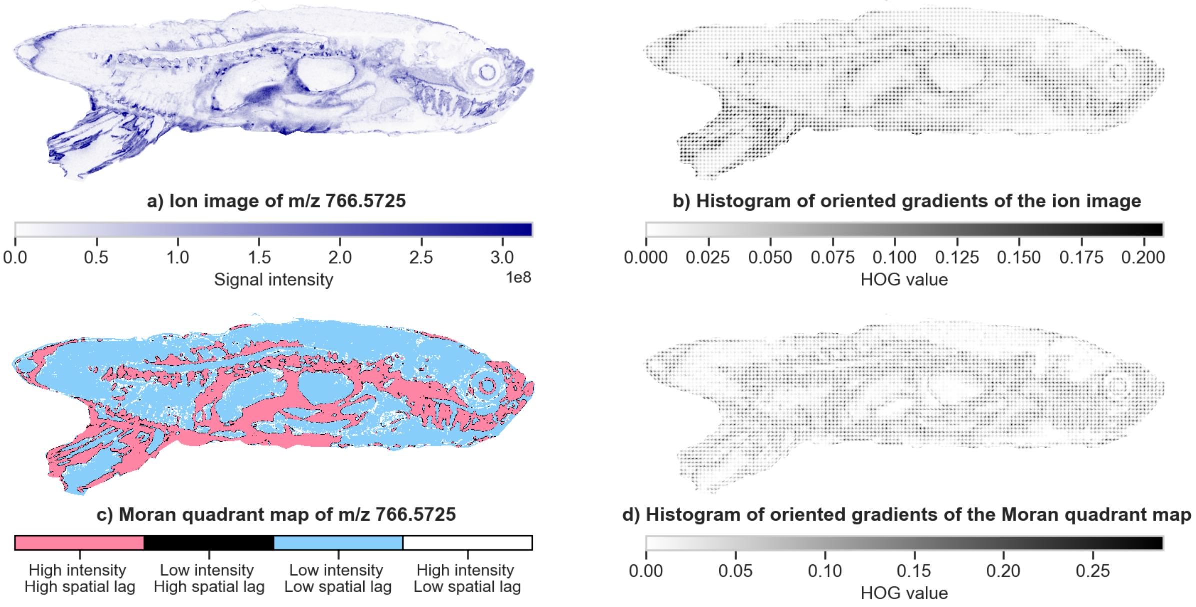

**Fig. S20**: Ion image of $m/z$ 766.5725, representative of cluster n$^o$7. The HOG features of the Moran quadrant map, rather than the HOG features of the ion image, were provided to the $k$-means clustering algorithm.

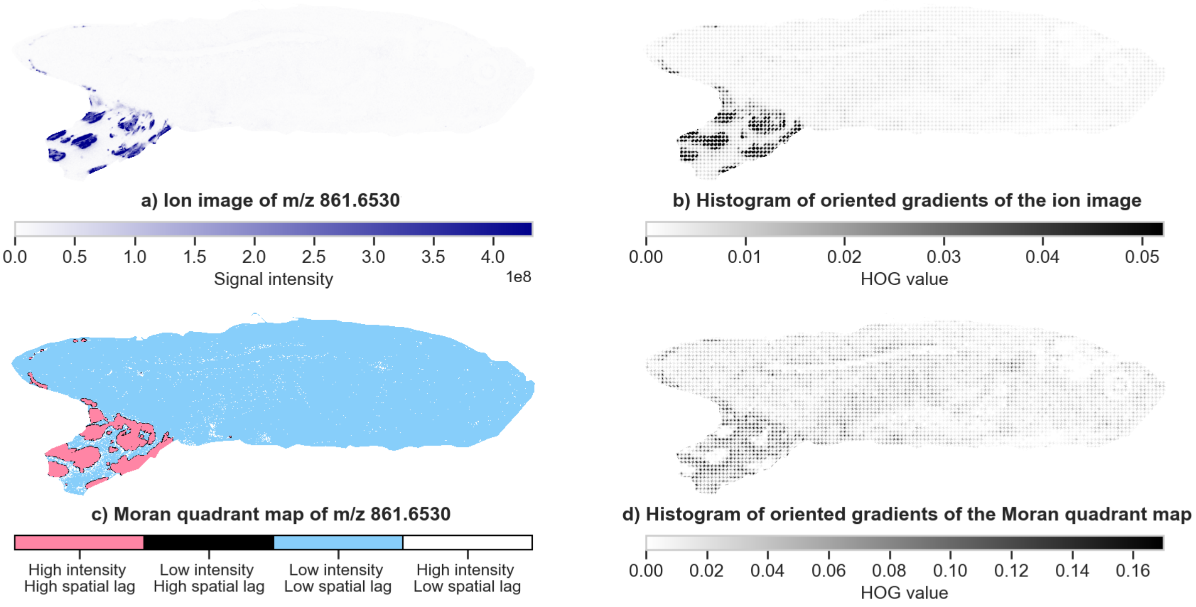

**Fig. S21**: Ion image of $m/z$ 861.6530, representative of cluster n$^o$8. The HOG features of the Moran quadrant map, rather than the HOG features of the ion image, were provided to the $k$-means clustering algorithm.

#### 5.3 IMS Dataset n$^o$3

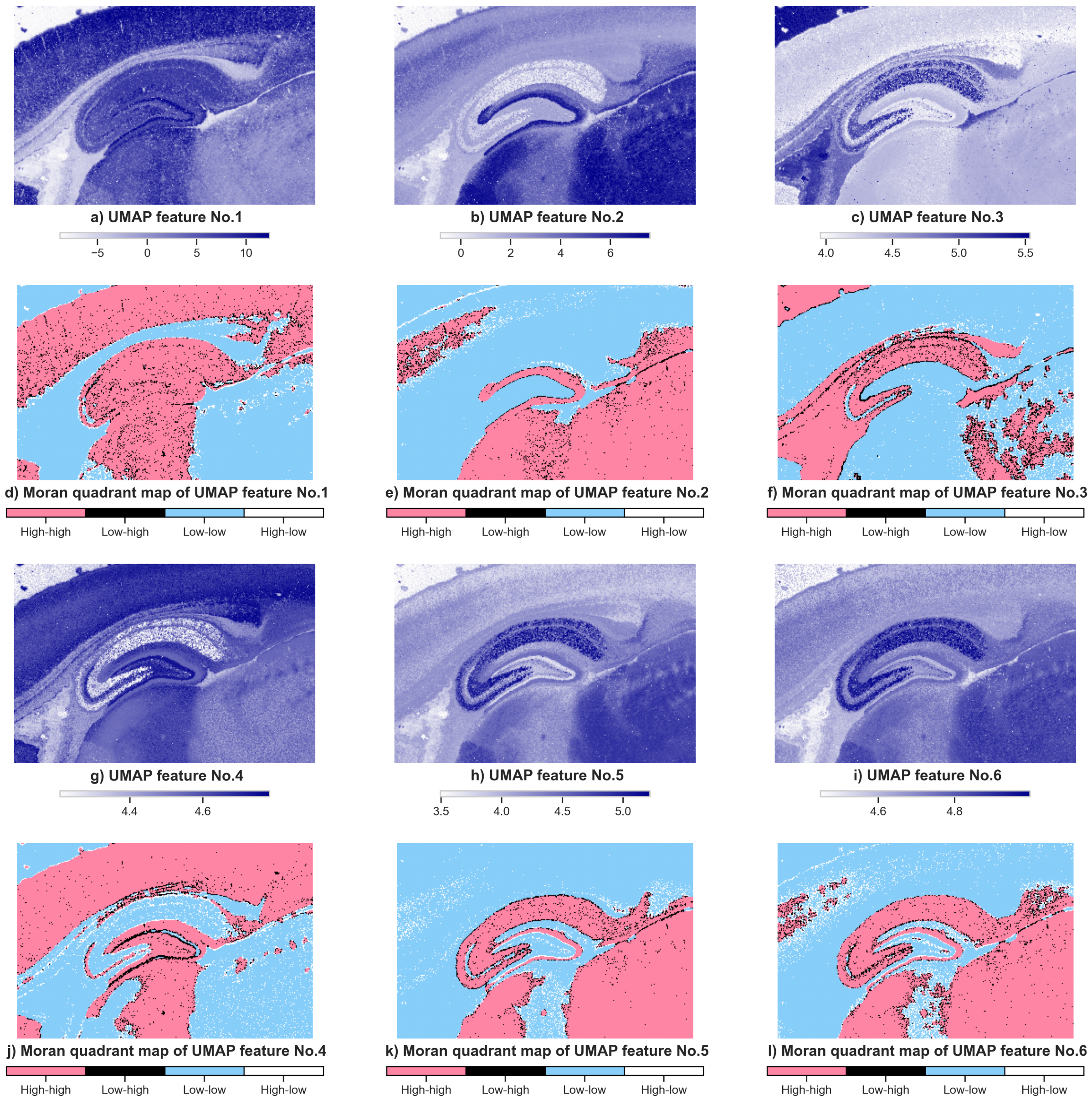

**Fig. S22**: Dimensionality reduction and processing prior to the segmentation of IMS dataset n$^o$3: the top-six UMAP features and their corresponding MQMs. The MQMs are obtained by computing the local Moran statistics of each UMAP feature using a cumulative 2$^{\text{nd}}$-order Queen-contiguity spatial weights matrix.

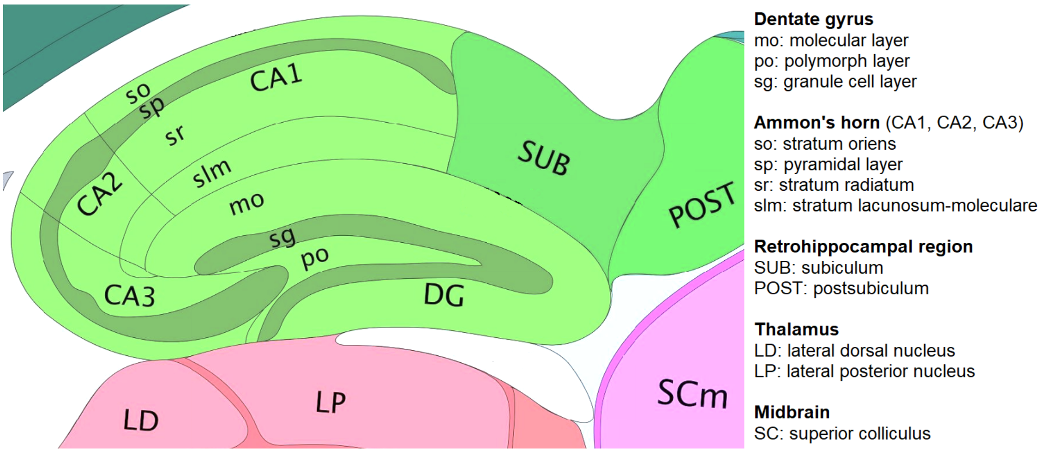

**Fig. S23**: Anatomical atlas of the sagittal section of a mouse hippocampus. The figure was adapted from the Allen adult mouse brain anatomical reference atlas [57].

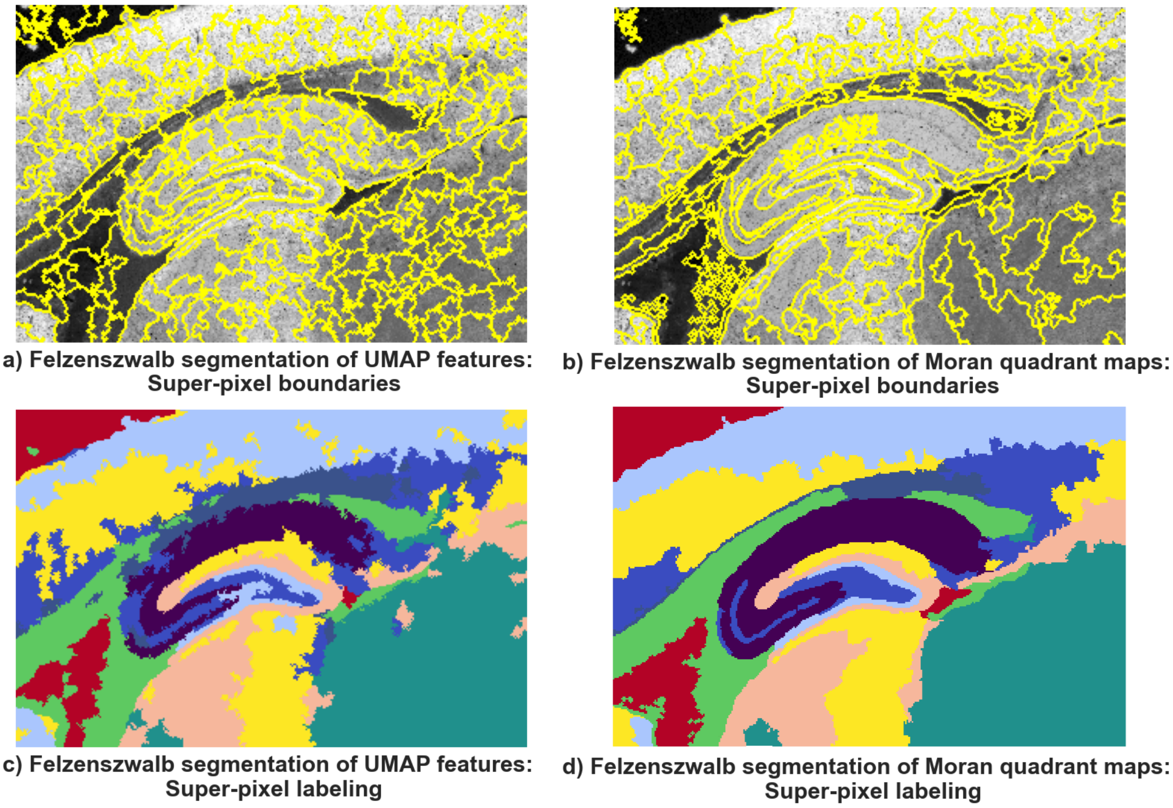

**Fig. S24**: Felzenszwalb segmentation versus Moran-Felzenszwalb segmentation of the top twelve UMAP features of dataset n$^o$3. **a** & **c**, The super-pixels are obtained by applying Felsenszwalb's segmentation algorithm to the top twelve UMAP features. **b** & **d**, The super-pixels are obtained by applying Felsenszwalb's segmentation algorithm to the MQMs of the top twelve UMAP features. The same hyper-parameters were used for the segmentation of the UMAP features and the MQMs. The MQMs were computed using a cumulative 2$^{nd}$-order Queen-contiguity spatial weights matrix. The super-pixels are labeled by majority voting: the label assigned to each super-pixel is the mode of the labels of the pixels that make up that super-pixel. The super-pixels obtained on the MQMs of the UMAP features are more faithful to the biological structure of the sample.

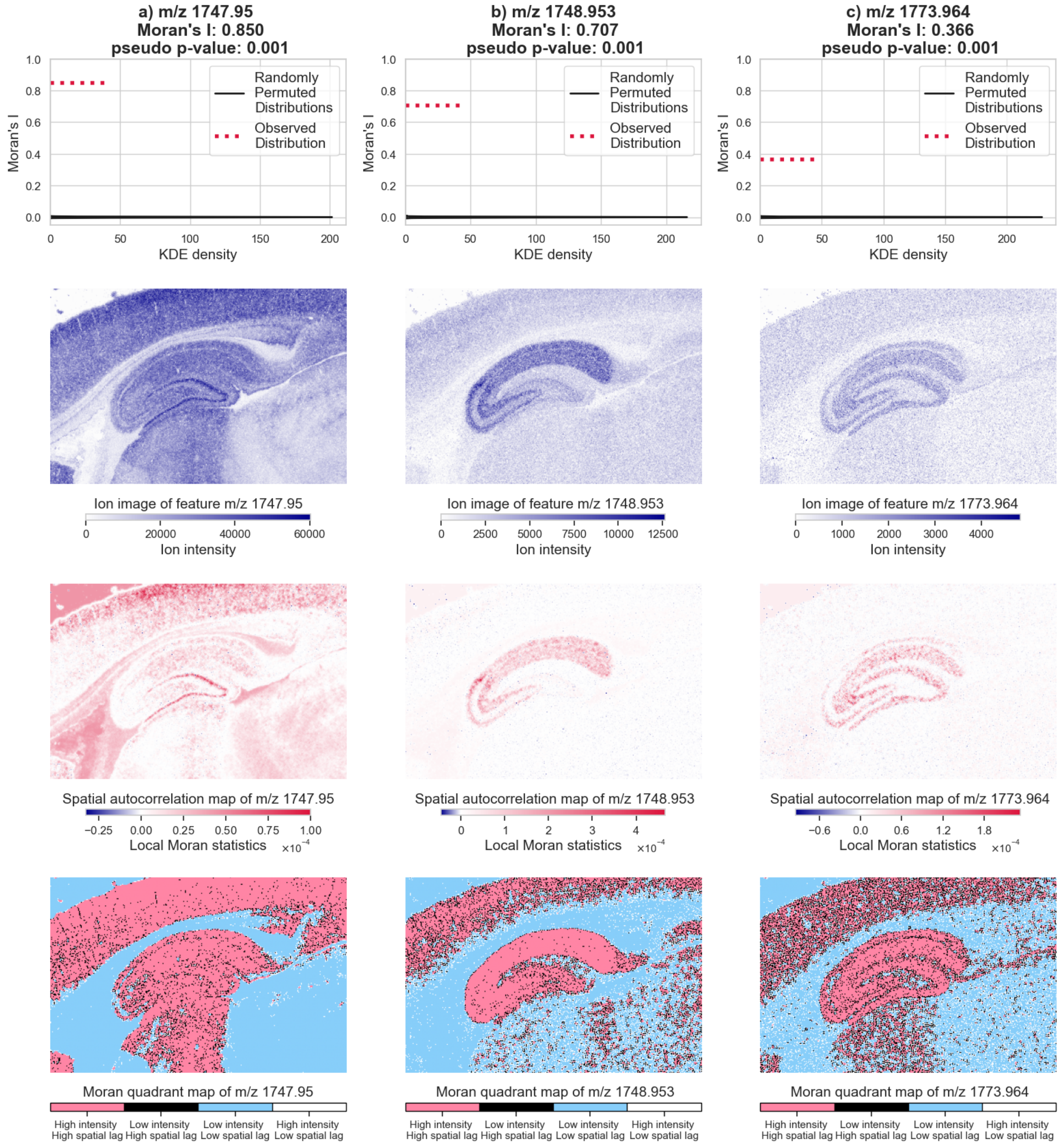

**Fig. S25**: Spatial analysis of three ion images from dataset n$^o$3 with a 1$^{st}$-order Queen-contiguity spatial weights matrix. The molecular species are ordered from left to right in descending order of global SAC, as measured by Moran's $I$. We observe that noisy ion images have low Moran's $I$ statistic. The statistical significance of each ion image's Moran's $I$ statistic is assessed in the first row: given that all three pseudo $p$-values are 0.001, we can conclude that the ion images of $m/z$ 1836.968, $m/z$ 1877.976, and $m/z$ 1776.981 are not spatially random. The second, third, and fourth rows present the ion images, spatial autocorrelation maps, and MQMs of the three ion images. The MQMs could serve as a basis for segmentation.

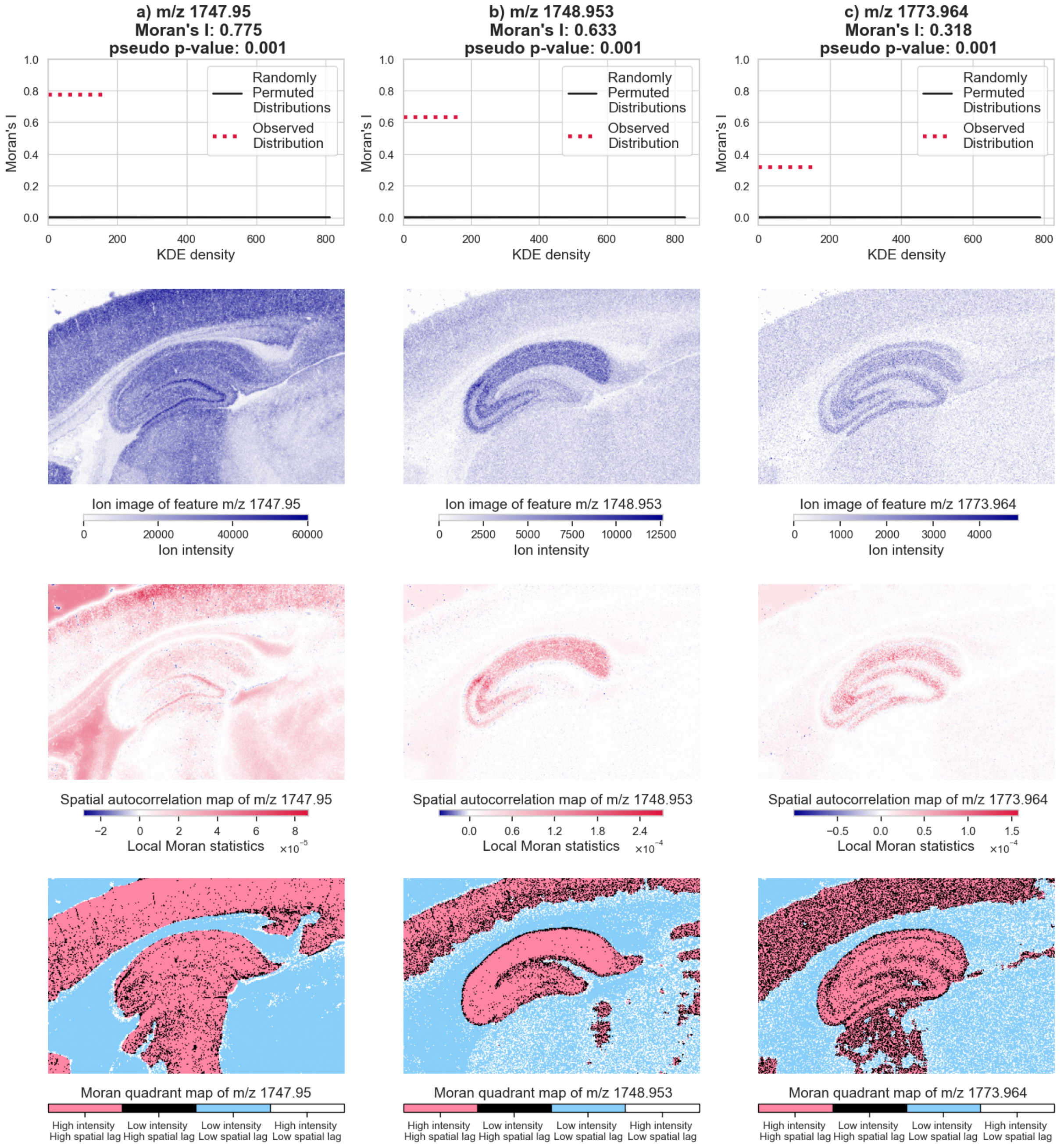

**Fig. S26**: Spatial analysis of three ion images from dataset n$^o$3 with a cumulative 5$^{th}$-order Queen-contiguity spatial weights matrix. The neighborhood of each pixel is defined as the union of its 1$^{st}$, 2$^{nd}$, 3$^{rd}$, 4$^{th}$, and 5$^{th}$-order neighbors as per the Queen-contiguity criterion, which is equivalent to defining a 5$^{th}$-order neighbor as a pixel that is located at a distance of either 1, 2, 3, 4, or 5 pixels away from the central pixel. In accordance with Tobler's first law of geography, we observe that global spatial autocorrelation decreases as the neighborhood order increases. We also observe that the MQMs of $m/z$ 1836.968, $m/z$ 1877.976, and $m/z$ 1776.981 that are obtained with a cumulative 5$^{th}$-order Queen-contiguity spatial weights matrix are less noisy than the corresponding MQMs obtained with a pure 5$^{th}$-order Queen-contiguity spatial weights matrix.

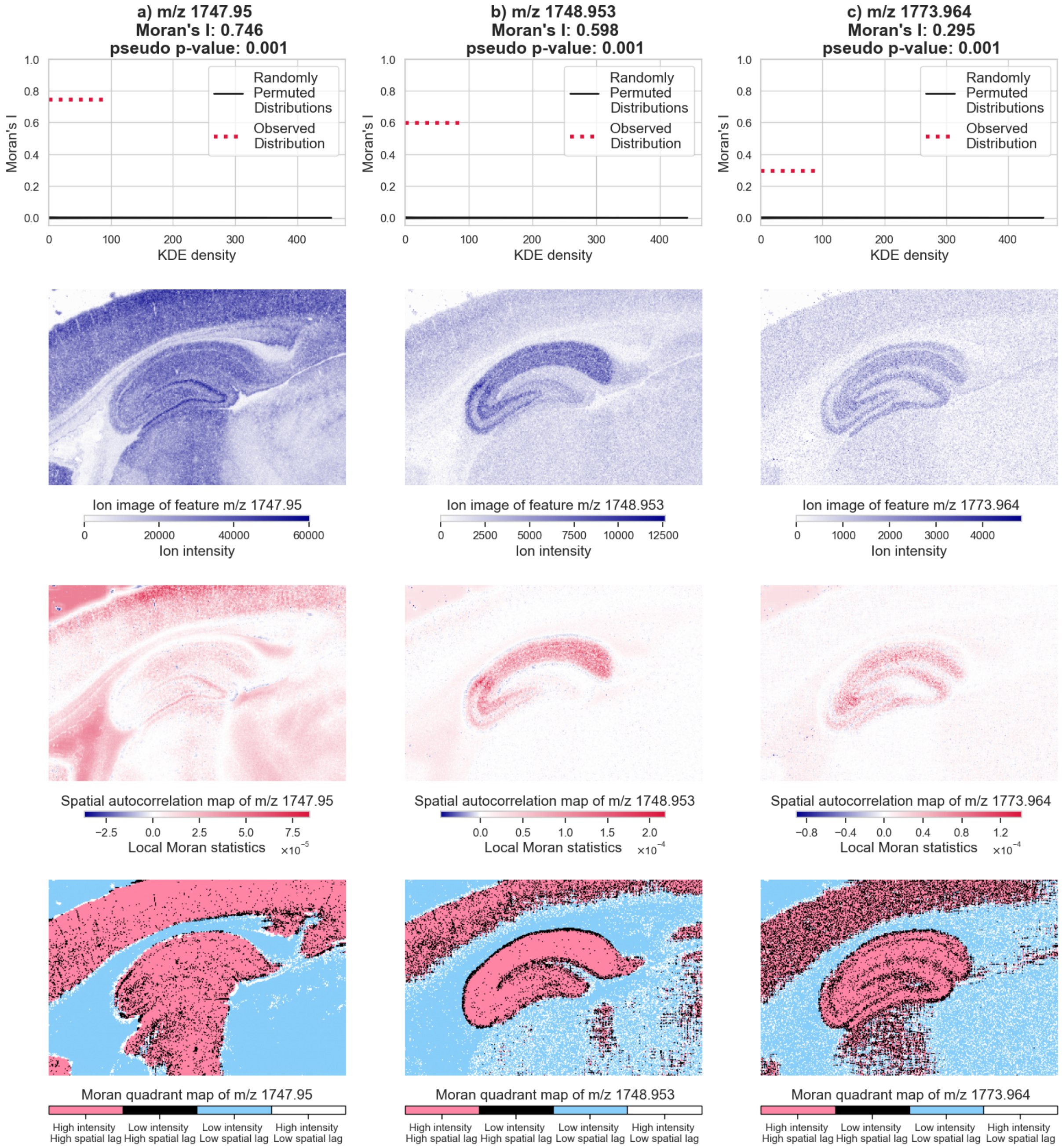

**Fig. S27**: Spatial analysis of three ion images from dataset n$^o$3 with a pure 5$^{th}$-order Queen-contiguity spatial weights matrix. Unlike the cumulative 5$^{th}$-order neighborhood used in Figure S26, a pure 5$^{th}$-order does not include lower-order neighbors (it only includes 5$^{th}$-order neighbors).
